# BromoCatch: a self-labelling tag platform for protein analysis and live cell imaging

**DOI:** 10.1101/2025.04.07.647551

**Authors:** Maria Rodriguez-Rios, Conner Craigon, Mark A. Nakasone, Adam G. Bond, Mark Dorward, Anthony K. Edmonds, Mark C. Norley, Robert E. Arnold, Paul M. Wood, Stephen J. Reynolds, Joel O. Cresser-Brown, Graham P. Marsh, Hannah J. Maple, Alessio Ciulli

## Abstract

Visualizing and manipulating proteins in live cells is crucial for studying complex biological processes. Self-labelling protein (SLP) tags such as HaloTag and SNAP-tag can be fused to genes of interest to allow protein labelling in cells. Limitations including size of the tag and suboptimal fitness of reactivity motivate development of improved tools to enable rapid, specific and stable protein labelling. We present BromoCatch, a novel SLP platform based on a small ∼13 kDa bromodomain (BD) engineered with a nucleophilic cysteine for covalent ligand engagement. A structure-based designed library of 16 “bumped” binders bearing diverse electrophilic warheads was screened against two different cysteine mutants using differential scanning fluorimetry and intact protein mass spectrometry to monitor covalent complex formation. The para-acrylamide bumped derivative MR116 and the Brd4-BD2 double mutant L387A,E438C formed the most potent and stable adduct, and its binding mode through covalent modification was confirmed by an X-ray cocrystal structure solved to 1.3 Å of resolution. BromoCatch exhibited potent and irreversible target engagement in cells through nanoBRET and residence time assays. Practicality and scope are further demonstrated through the design and proof-of-concept application of a biotinylated conjugate, PROTAC tag degraders, and fluorescent probes of both full-on and switch-on types for ex-cellulo and live-cell imaging. Together, we qualify BromoCatch as a novel, versatile and efficient protein labelling tool and technology platform. Its advantageous design features and kinetic fitness, and its modular design enabling diverse functionalities, are anticipated to usher a range of future applications and witness broad utility.

## Introduction

The ability to visualize and manipulate proteins in live cells is critical for studying biology and support drug development. Multiple technology platforms have been developed for this purpose. Self-labelling protein tags (SLP) have emerged as powerful, versatile and technically accessible tools that have been widely adopted for both *in vitro* and *in vivo* protein research^1^. The principle behind SLPs is the combination of a protein tag, expressed as a fusion to the protein-of-interest, and a complementary small molecule ligand (‘tag ligand’) that potently (typically covalently) and selectively reacts with the protein tag (Figure 1). The key advantage of the technology is the intrinsic versatility of SLPs due to the small molecule ligand component, which can be modified to incorporate a wide variety of handles including fluorophores, biotin, click-chemistry handles, E3 ligase ligands for targeted degradation, and orthogonal domain-binding ligands for induced proximity applications (**Figure 1A**). SLPs have a broad range of applications, including fluorescent labelling for live-cell imaging, Förster resonance energy transfer (FRET) assays for studying protein-protein interactions, protein turnover and degradation studies, biotinylation, and affinity purification^2^. Their utility even extends to *in vivo* imaging, making them powerful and versatile tools for both basic research and translational applications.^3–8^

**Figure 1.**
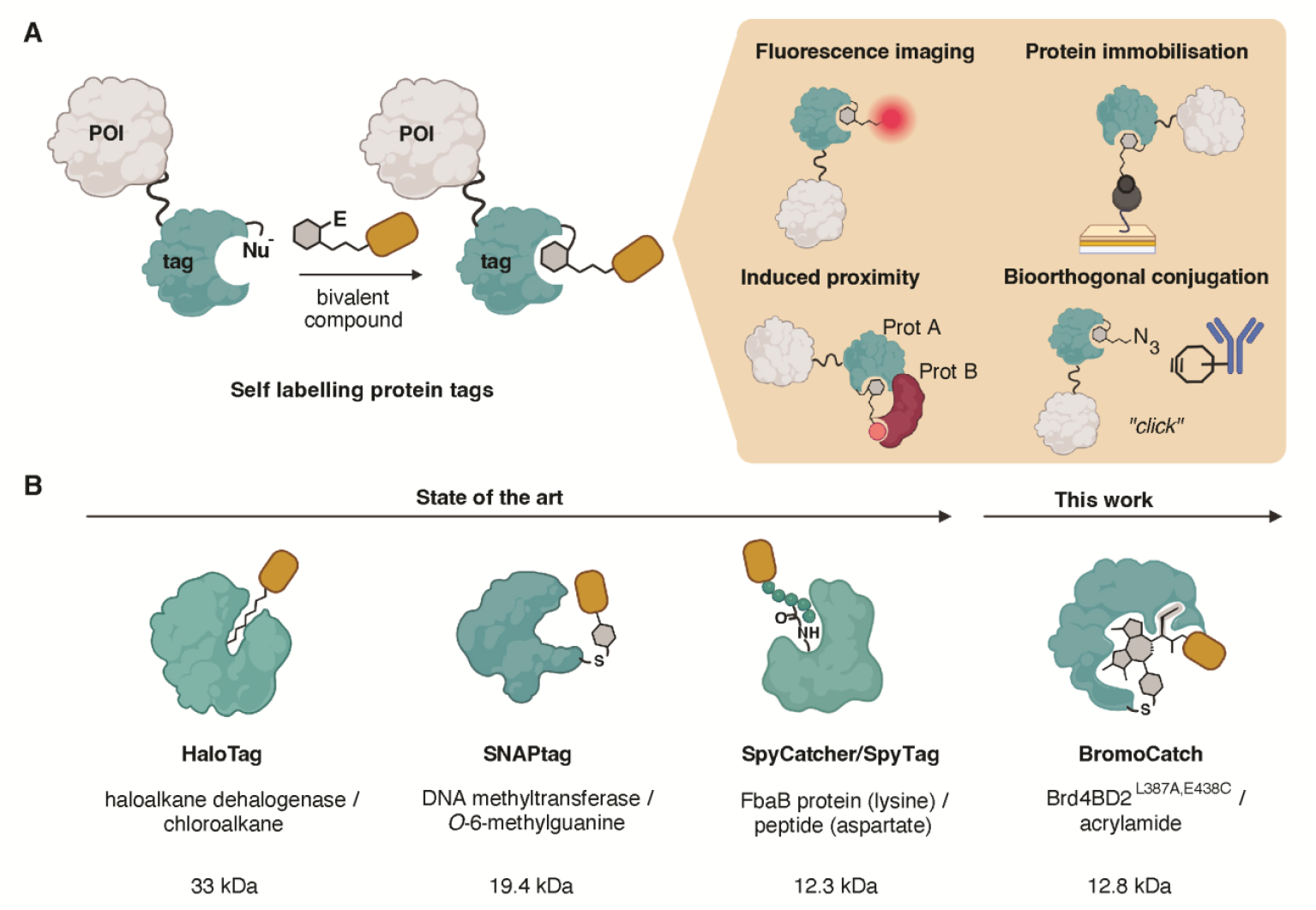
Self labelling tag platforms for protein analysis and manipulation. A) Principle of self labelling tags principle. B) Current self-labelling protein tags prior art and introducing BromoCatch described herein.

An important application of SLPs is in live-cell imaging experiments, enabled through pairing tag ligands with fluorophores. This approach offers several benefits over creation of fusion proteins with fluorescent proteins such as GFP for the same application, primarily the superior photophysical properties offered by synthetic fluorophores (specifically brightness and photostability), and the ability to switch-on the fluorescent signal with temporal control upon simple addition of compound. Breakthrough advances in imaging technology and instrumentation now place greater demands on the fluorophore used and render fluorescent proteins fundamentally sub-optimal for many advanced and super resolution microscopy techniques. Further, it is well documented that the relatively large molecular weight of GFP (27kDa, often larger than the fusion partner protein of interest) can confound experiments by altering the interactions, localization and/or function of the protein partner. ^9–13^

The most popular SLPs currently in use are HaloTag^14,15^, SNAP-tag^16,17^ and CLIP-tag^18^ (**Figure 1B**). HaloTag developed by Promega is engineered from the bacterial haloalkane dehalogenase (DhaA) from Rhodococcus. Through mutagenesis, Wood et al. ^14,19^ engineered DhaA (H272F) to irreversibly trap the covalent ester intermediate within the enzyme mechanism at the nucleophilic residue D106, by deactivating the second step in the mechanism through the catalytically inactive phenylalanine at position 272. Despite its relatively large size (33 kDa), HaloTag’s net negative charge at pI=5 enables it to function as a solubility tag in protein purification with reduced tendency towards aggregation. SNAP-tag, developed in Kai Johnsson’s lab, is derived from human O6-alkylguanine-DNA alkyltransferase (hAGT). Through directed evolution and random mutagenesis, Johnsson et al. developed the SNAP-tag, which has high specificity for O6-benzylguanine-equipped ligands. At a size of 19.4 kDa, SNAP-tag is less likely to cause less perturbation of tagged proteins compared to HaloTag^20^. A more recent derivation of SNAP-Tag is CLIP-tag^16^. Through mutagenesis of eight amino acids in SNAP-tag, CLIP-tag was created, reacting with high speed and selectivity with O2-benzylcytosine derivatives. The creation of CLIP-tag was motivated by the need for a tag compatible with SNAP-tag, allowing for multiplexing of two proteins in the same cell, given the limited number of SLPs available at the time^16^. The covalent TMP-tag^21^ has recently gained popularity following the optimization of its originally reversible version^22^. This system utilizes *Escherichia coli* dihydrofolate reductase (eDHFR; 18 kDa) as a fusion tag and employs the folate analog trimethoprim (TMP) for labeling.

The HaloTag and SNAP/CLIP-tag platforms have thus emerged as gold standard and are an integral part of the chemical biologist’s toolbox. There are limitations, however, associated with both systems. HaloTag is a relatively large (33kDa) protein tag, and it is expected that fusion of such a large domain may perturb the protein of interest ^21,23,24^. Indeed, altered or impaired protein function following incorporation of the HaloTag7 protein has been reported following conjugation to the CB2 receptor,^25^ AGO2, ^26^ COX8A and ATP5ME^21^The SNAP-tag exhibits slower labelling kinetics and exhibits reduced fluorescence with rhodamine substrates compared to HaloTag, additionally the benzyl guanine ligand is not optimally cell permeable, requiring typically higher ligand concentrations and additional washing steps. We note that recent improvements have been made to this platform to try to address some of these limitations.^27^ The TMP-tag is the smallest non-peptide-based SLP, but its covalent reaction depends on NADPH, the native cofactor of DHFR, which limits its use to intracellular applications. Finally, the SLP ligands typically used are not considered ‘drug-like’ and are associated with poor pharmacokinetic properties, specifically rapid clearance, which can present challenges during *in vivo* application.^28^

In this work we aimed to address the limitations of existing SLPs and expanding the toolbox by developing an orthogonal platform for multiplexed tag applications (**Figure 1B**). Inspired by our previous ‘bump-and-hole’ approach applied to Brd4-BD2 to generate a functionally silent, ‘hole-containing’ point mutant (Brd4-BD2^L387A^) and a highly specific and potent small molecule ligand (ET-JQ1-OMe) ^29,30^ we sought to modify this system to introduce covalency, while simultaneously reducing the molecular weight of the protein tag. Our lab’s BromoTag degron system,^31^ based on the bump and hole approach, enables selective targeting of a mutant Brd4-BD2 L387A bromodomain using a complementary bumped non-covalent PROTAC^31^. This platform facilitates potent, selective, and rapid degradation of tagged proteins, proving valuable in targeted protein degradation (TPD)^32–36^ and for induced proximity approaches beyond TPD^37^. Here, we describe our development of BromoCatch, a self-labeling tag platform expanding applications beyond degradation. By engineering Brd4-BD2^L387A^ to introduce a nucleophilic cysteine amino acid (Brd4-BD2^L387A, E438C^) at a suitable location and incorporating an electrophilic warhead into the bumped ligand, we created a covalent and highly specific SLP system.

## Results and Discussion

The initial design of the system focused on finding residues in or around the binding pocket of the ‘hole containing’ protein mutant Brd4-BD2^L387A^ that could be mutated to incorporate a cysteine that oriented near a suitable vector of the ET-JQ1-OMe (**Figure 2A**).^31,38^ To evaluate the Brd4-BD2 binding pocket, we used an existing co-crystal structure of Brd2-BD2^L383V^ bound to ET-JQ1-OMe (PDB code: 6YTM). Brd2-BD2 has exquisite structural homology and sequence identity at the ligand binding site compared to Brd4-BD2, thus offering appropriate surrogate to structurally interrogate the Brd4-BD2 binding pocket (**Supplementary Figure 1**). Residues Met442 and Glu438 in Brd4-BD2 are located within 6 Å of the pendant aryl ring in ET-JQ1-OMe and were therefore identified as attractive sites for introducing cysteine mutations. We rationalized that the distance to the pendant ring (6 Å) would be sufficient to enable nucleophilic attack of a suitable electrophilic warhead, introduced into the phenyl ring. (**Figure 2B**). To test this hypothesis, two Brd4-BD2 mutants were designed, keeping the ‘hole’ (L387A) and incorporating a nucleophilic cysteine by additional mutation of E438C (**Figure 2C**) or M442C (**Figure 2D**). Complementarily, a series of 16 electrophilic derivatives of the ET-JQ1-OMe were designed that maintain the ethyl ‘bump’ and incorporate an electrophilic warhead either at the *para* or *meta* positions of the pendant aryl ring (**Figure 2E**).

**Figure 2.**
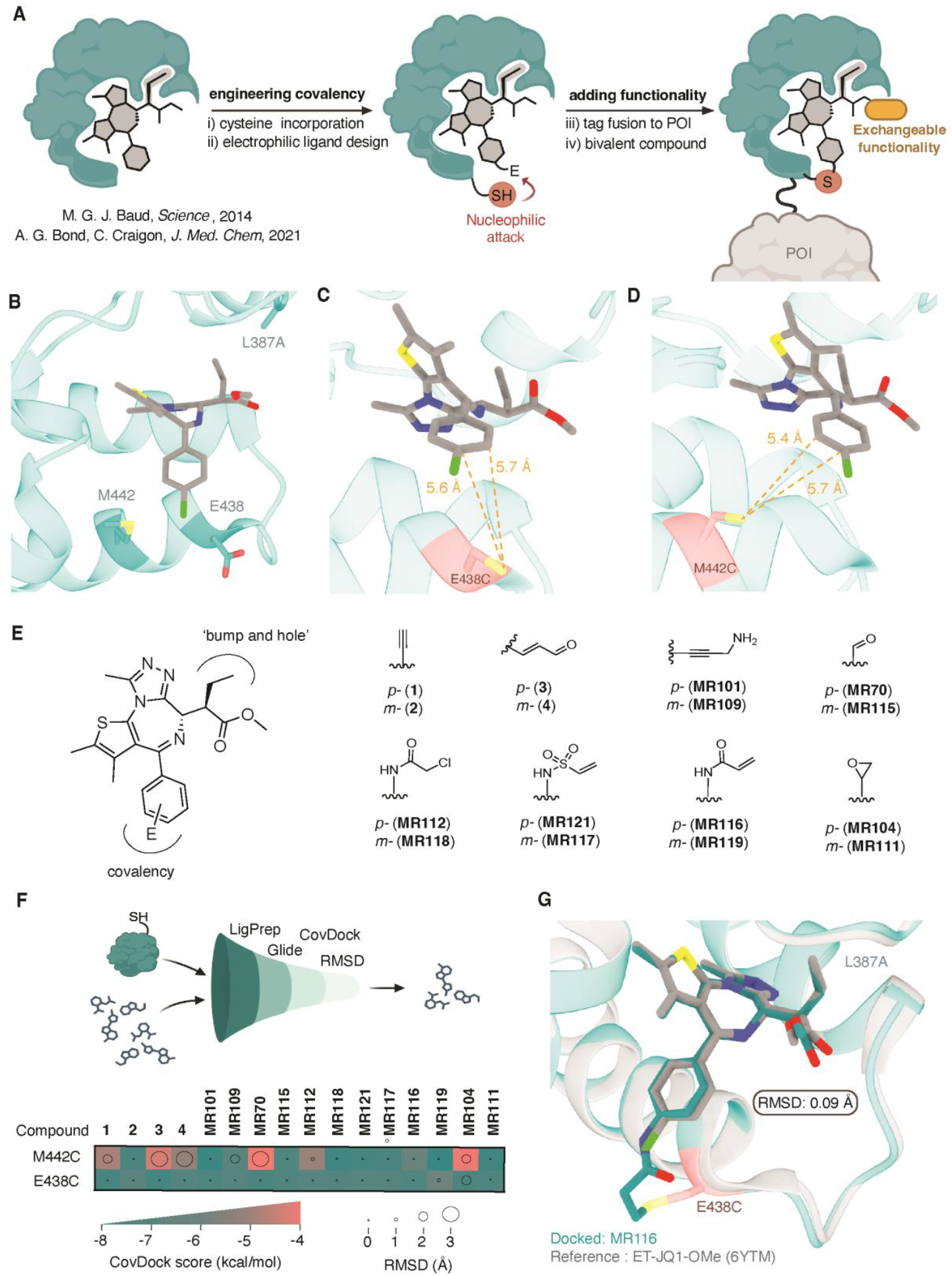
BromoCatch system design, development and covalent docking. A) Rational design of the system B) Brd4-BD2 identified amenable residues M442 and E438. C) Cysteine mutant distances for the E438C mutant to *meta* and *para* positions of the aromatic ring. D) Cysteine mutant distances for the M442C mutant to *meta* and *para* positions of the aromatic ring. E) Library of *meta* and *para* electrophilic compounds. F) *In silico* evaluation workflow and heatmap summarizing covalent docking scores and RMSDs for E438C and M442C mutants. G) Covalent docking of MR116 (teal carbons) as predicted bound to Brd2-BD2^L383A,D434C^ and 3D alignment with non-covalently bound Et-JQ1-OMe (grey carbon) in co-crystal structure 6YTM (RMSD CS 0.09 Å).

To gain an indication of the system viability, the designed protein mutants and the library of ligands were prepared *in silico* (Schordinger Suite, LigPrep) and evaluated by reversible (Glide)^39^ and covalent docking (CovDock)^40^. Starting from the ET-JQ1-OMe : Brd2-BD2^L383V^ co-crystal structure (PDB code: 6YTM), both L387A,E438C and L387A,M442C double mutants were generated and screened *in silico* against the entire library of *meta* and *para* ligands. The results of this screen are shown in **Figure 2F**, where CovDock scores and RMSD over the bound ligand atoms are monitored to predict to likelihood of a given system to form strong covalent binding while retaining the known favourable ligand-binding mode and non-covalent interactions. The most striking trend observed in the covalent docking screen is that the E438C mutation (D434C in Brd2-BD2, **Supplementary Table 1**) resulted in better covalent binding scores and much lower RMSD values than M442C, suggesting that the cysteine residue position/orientation in the binding pocket is important and supporting E438C as the preferred mutant. Of note, the differential predicted reactivity for the two cysteines, based on calculated pKa values of E438C = 8.4 and M442C = 8.9, may contribute to the trend observed. When comparing the *meta* or *para* position for a given electrophilic warhead (e.g. MR116 vs MR119) we did not observe major differences or trends between the two, and both analogues yielded highly favorable energy scores and very low RMSD especially for the E438C mutant (**Figure 2F**), suggesting that the electrophilic warhead can react from either positions. Overall, the covalent docking screen suggests that both engineered cysteine mutants should react well with most of the designed electrophilic ligands, with the E438C predicted as preferred. As an example of the promising system design, the docked acrylamide **MR116** showed exquisite overlap in binding mode when superposed with the reference non-covalent ligand (RMSD_CS_ 0.09) and a high CovDock score of -7.29 kcal/mol (**Figure 2G**).

With the promising results from the *in silico* studies, the newly designed mutant proteins were taken forward for recombinant protein expression. To achieve our goal of minimizing tag molecular weight, we removed 17 amino acids of the N-terminus before the bromodomain (333-350) in a region that was expected to have no detrimental effect on stability while maintaining all the structural regions of Brd4-BD2 intact (**Supplementary Figure 2**). The short mutants Brd4-BD2^L387A,E438C^ (351-459) and Brd4-BD2^L387A,M442C^ (351-459) were successfully expressed and purified from *E. coli* and their stability tested compared to the longer Brd4-BD2^WT^ and Brd4-BD2^L387A^ (333-459). The shortening of the sequence and introduction of the double mutation (L387A and M442C or E438C) had no apparent detrimental effect on overall stability, with the mutant showing relatively similar melting temperature (Tm) values to that of the Brd4-BD2^WT^ and Brd4-BD2^L387A^ (**Supplementary Figure 2**). Based on these results, both double mutants Brd4-BD2^L387A,E438C^ and Brd4-BD2^L387A,M442C^ were taken forward.

**Scheme 1.**
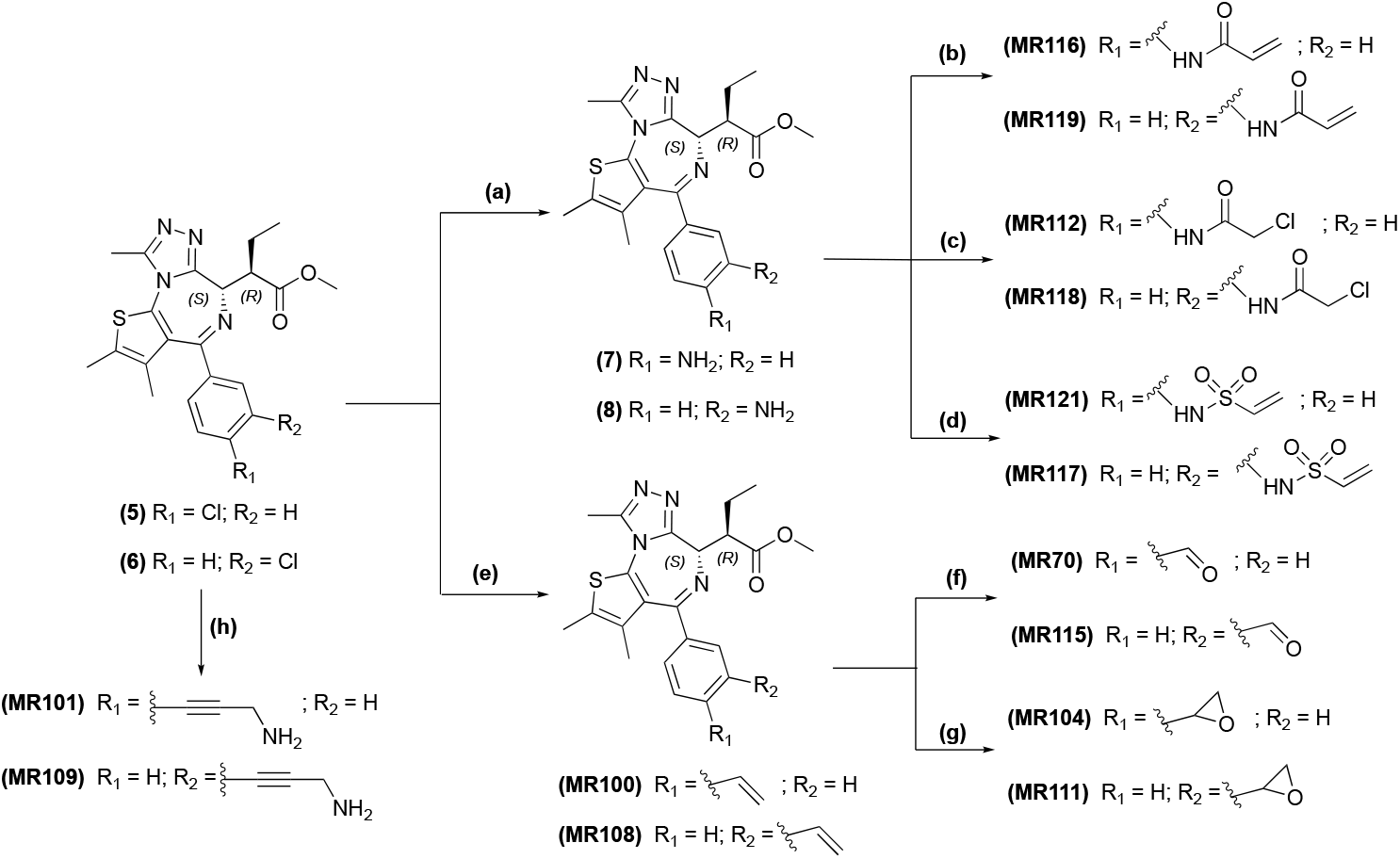
Synthesis of electrophilic bumped ligand library. Reagents & Conditions. (a) (i) benzophenone imine, Pd_2_(dba)_3_, *t*-BuXPhos, K_3_PO_4_, 1,4-dioxane, 85 °C; (ii) 1M HCl (aq.), THF, RT; (7, 50%) (8, 50%); (b) acryloyl chloride, DIPEA, DCM, RT; (MR116, 71%), MR119, 59%); (c) chloroacetyl chloride, DIPEA, DCM, RT; (MR112, 67%) (MR118, 71%); (d) vinyl sulfonyl chloride, pyridine, DCM, 0 °C; (MR121, 30%) (MR117, 54%); (e) potassium vinyltrifluoroborate, XPhos Pd G2, DIPEA, DMF/H_2_O (1:1), 90 °C; (MR100, quant.) (MR108, 91%) (f) OsO_4_, NaIO_4_, acetone/H_2_O (5:1), RT; (MR70, quant.) (MR115, quant) (g) *m*CPBA, DCM, RT; (MR104, 10%) (MR111, 7%) (h) propargyl amine, XPhos Pd G2, Cs_2_CO_3_, THF, 90 °C (MR101, 20%; MR109, 13%)

A library of the *in silico*-designed electrophilic ligands was synthesized, functionalising the ET-JQ1-OMe scaffold by incorporating a range of electrophilic handles. The complementary diazepine scaffolds (5) and (6) were prepared analogously to the protocols of Bond *et al*.,^38^ with subsequent derivatization at the *para-* or *meta-*position of the pendant aryl ring, respectively (**Scheme 1**). Buchwald-Hartwig amination with benzophenone imine and subsequent hydrolysis produced anilines (7) and (8), which could readily be converted to the respective acrylamide (MR116 and MR119), chloroacetamide (MR112 and MR118) and vinyl sulfonamide (MR121 and MR117) ligands in high yield. In similar fashion, a potassium vinyl trifluoroborate-promoted Suzuki-Miyaura coupling was used to access styrene intermediates (MR100) and (MR108), in-turn serving as convenient precursors of epoxides (MR104) and (MR111) and offering facile access to aldehydes (MR70) and (MR115) by means of oxidative cleavage. Finally, Sonogashira cross-coupling with propargyl amine gave rise to the GNE-0011 analogues ^41,42^ (MR101) and (MR109), completing the panel.

To evaluate the reactivity of the different electrophilic ligands towards both engineered cysteine-containing mutants, covalency was systematically assessed using ESI-MS (**Figure 3**). Binding resulted in a shift in LC retention time (280 nm) giving resolvable peaks that could be separately processed to obtain the corresponding mass spectrum, that could be subsequently deconvoluted to calculate the mass of the protein (**Figure 3A**). Ligands that exhibited poor or no covalent binding showed no retention time shift, and no mass changes were detected or were only present in trace amounts (<5%). Excitingly, several electrophiles (acrylamide, chloroacetamide, vinyl sulfonamide and epoxide) showed covalent modification of both cysteine containing mutants Brd4-BD2^L387A,E438C^ and Brd4-BD2^L387A,M442C^ (**Figure 3B, Supplementary Table 2)**. Importantly, the compounds did not covalently modify Brd4-BD2^WT^ or Brd4-BD2^L387A^, indicating high selectivity for the newly introduced cysteines. Brd4-BD2^L387A,E438C^ exhibited excellent reactivity towards both *meta* and *para* electrophilic ligands, likely due to the position of the engineered cysteine 438 in a relatively flexible region of the binding pocket. Conversely, cysteine 442 in Brd4-BD2^L387A,M442C^ was substantially less reactive for many of the compounds tested, possibly because this cysteine is located in a more rigid region (alpha-helix C) and has a slightly higher calculated pKa (8.9) compared to the cysteine 438 in Brd4-BD2^L387A,E438C^ (calculated pKa=8.4). Interestingly, *para*-substituted compounds performed better than *meta*-substituted for Brd4-BD2^L387A,M442C^.

**Figure 3.**
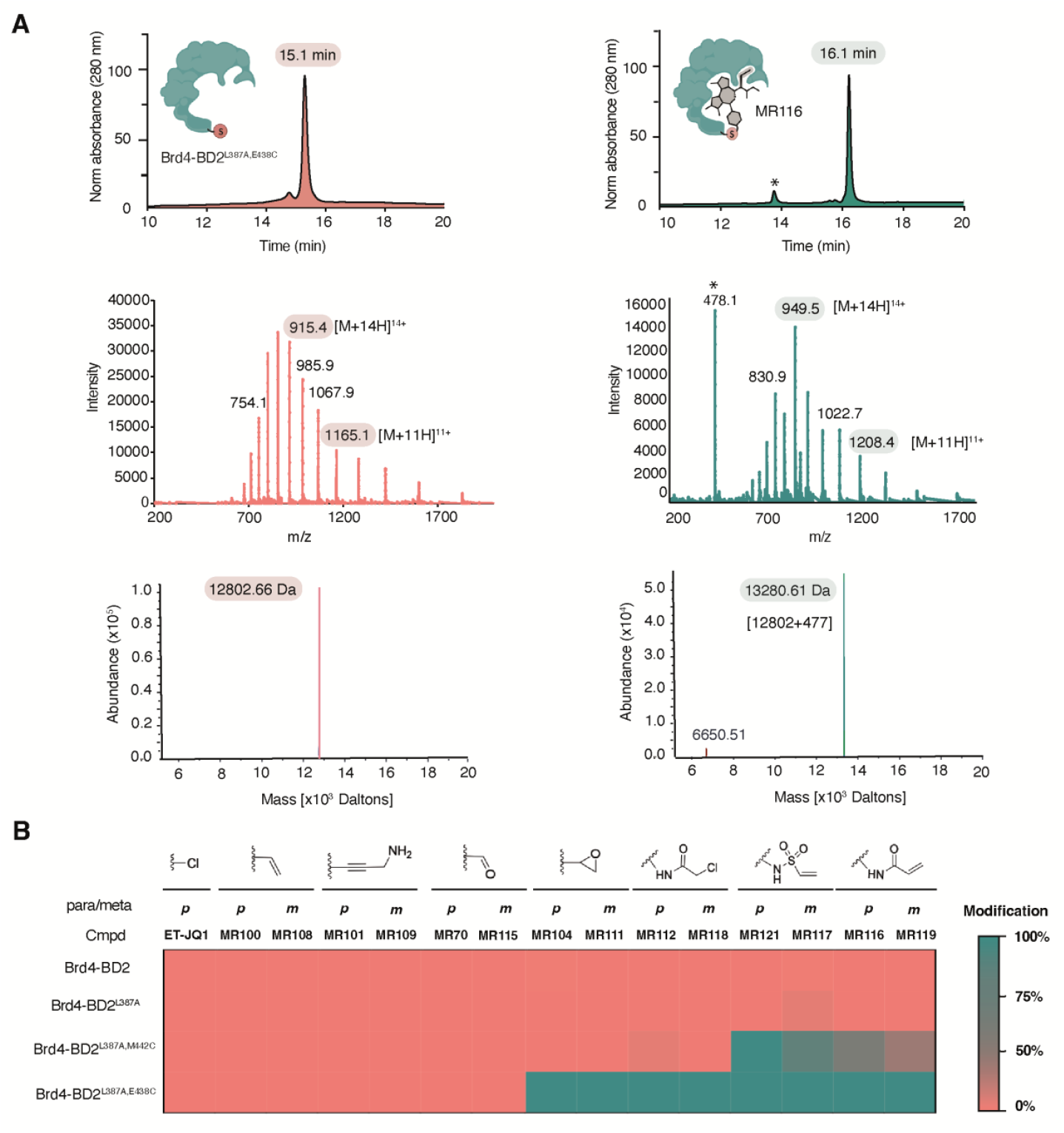
Library screen by intact protein ESI-MS to monitor covalent adduct formation. A). Brd4-BD2^L387A,E438C^ protein was flown on ESI-MS either in the absence (left) or presence (right) of MR116. UV (280 nm) peak shift (top), m/z envelope change (middle) and deconvoluted mass (bottom) are shown. Asterisk indicates the mass of the MR116 ligand. B) Heatmap shows the results of the systematic studies for covalent modification of all the proteins against all synthesized ligands including controls of non-electrophilic compounds (ET-JQ1 and styrene compounds).

To estimate the relative binding affinity of the component of the ligand library, a differential scanning fluorimetry (DSF) screen was performed to measure the compound-induced thermal stabilization of the different variants of the bromodomain protein (**Figure 4**). The increment in melting temperature (ΔT_m_) for the protein in presence of the ligand was measured relative to that of the corresponding unliganded protein (DMSO-only control) - representative DSF traces of four protein variants with compound MR116 are shown in **Figure 4A**. All tested compounds provided poor stabilization of Brd4-BD2^WT^, confirming the expected effect of the ethyl bump abrogating reversible binding to the bromodomain. Most of the compounds maintained the ‘bump and hole’ reversible binding against the Brd4-BD2^L387A^ (ΔTm ∼10 °C), as expected. Crucially, much greater stabilization of the cysteine bearing mutant could be observed for several ligands, with ΔTms measured as high as +28 °C and more prevalently with the Brd4-BD2^L387A, E438C^ mutant (**Figure 4B, Supplementary Table 3**). This augmented thermal shift is consistent with the enhanced stability from covalent adduct formation. Of note, the pattern of extra stabilization observed on ΔTms (**Figure 4B**) correlated remarkably well with the trends in reactivity observed by ESI-MS (**Figure 3B**). The non-electrophilic and poorly electrophilic ligands maintained the ‘bump and hole’ non-covalent binding mode for both cysteine containing mutants without showing any additional stabilization (ΔTm +10 °C), confirming lack of covalent adduct formation. In a similar trend to that observed the ESI-MS analysis, the stabilization effect for Brd4-BD2^L387A,M442C^ was less pronounced, with some of the strong electrophiles showing comparatively poor stabilization (*meta* chloroacetamide and *meta* sulphonamide). Taken together, our structure-activity screens reveal that the mutant Brd4-BD2^L387A,E438C^ in combination with the *para* chloroacetamide (MR112), vinyl sulphonamide (MR121) and acrylamide (MR116) performed as best combinations, providing highly strongest covalent binding.

**Figure 4.**
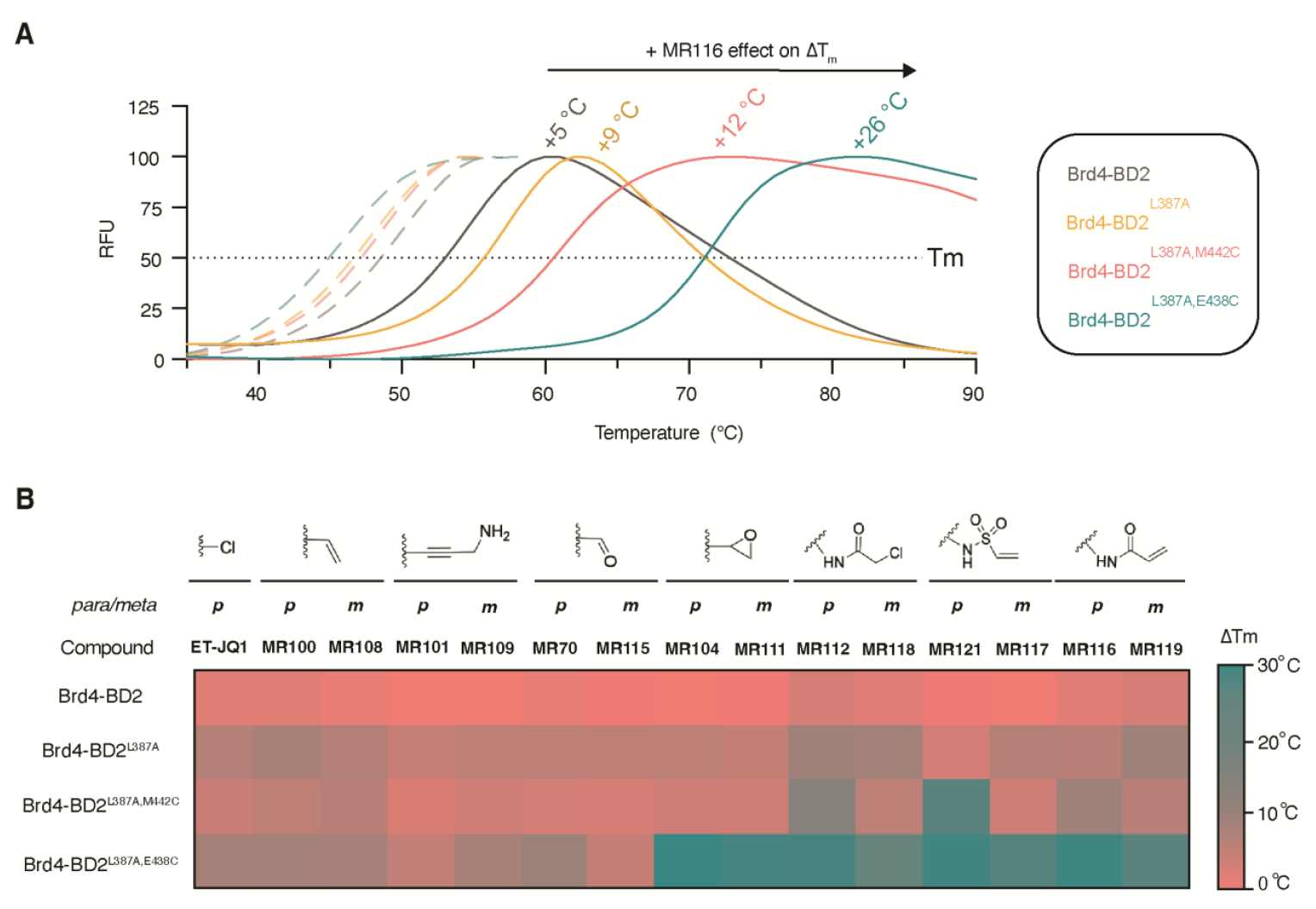
Library screen by Differential Scanning Fluorimetry to monitor ligand-induced protein thermal stabilization. A) DSF melting curves for the different mutant and wild-type proteins in absence or presence of compound MR116. B) Heatmap shows the results of the systematic studies for ΔTm for the different proteins against all synthesized ligands including controls of non-electrophilic compounds (ET-JQ1 and styrene compound MR100).

Based on its high reactivity and superior stabilization with most ligands tested, we elected Brd4-BD2^L387A,E438C^ as our protein tag of choice. The acrylamide MR116 was selected as the specific complementary covalent ligand given the optimal specific covalent modification and superior stabilization observed with Brd4-BD2^L387A,E438C^. Acrylamides have excellent stability *in cellulo*, low cross-reactivities and are widely used in chemical probes,^43,44^ with multiple compounds in clinical development and FDA approved drugs containing acrylamides as covalent warheads.^45,46^ We named the system ‘BromoCatch’, with the Brd4-BD2^L387A,E438C as^ our BromoCatch fusion protein.

To confirm covalency at a molecular level and evidence the expected binding mode, the BromoCatch protein was co-crystallized in the presence of the acrylamide MR116. Since no crystals were obtained for the Brd4-BD2^L387A,E438C^ : MR116 complex despite multiple co-crystallization attempts, we decided to switch to the highly homologous Brd2-BD2^L383A,D434C^, because Brd2-BD2 had previously been successful in our hands in yielding co-crystal structures with bump and hole ligands. ^29,38,47,48^ The screens with the Brd2-BD2^L383A,D434C^ : MR116 complex resulted in high quality crystals within hours. The Brd2-BD2^L383A,D434C^/MR116 co-crystal showed exquisite overlap with the reference crystal structure (Brd4-BD2^L383V^/ET-JQ1-OMe, PDB code: 6YTM) with an RMSD value for the common benzodiazepine below 1 Å, suggesting that the binding mode of the benzodiazepine scaffold is maintained. Inspection of the electron density around the ligand clearly evidence covalent bond between the ligand and the engineering cysteine (**Figure 5B**), as continuous electron density (1sigma 2Fo-Fc) at an atomic level is observed consistent with formation of a thioether bond. Closer inspection of the omit difference map (polder map) suggests two alternate conformations are possible for the thioether bond of the adduct (**Figure 5C, Supplementary Figure 4**).

**Figure 5.**
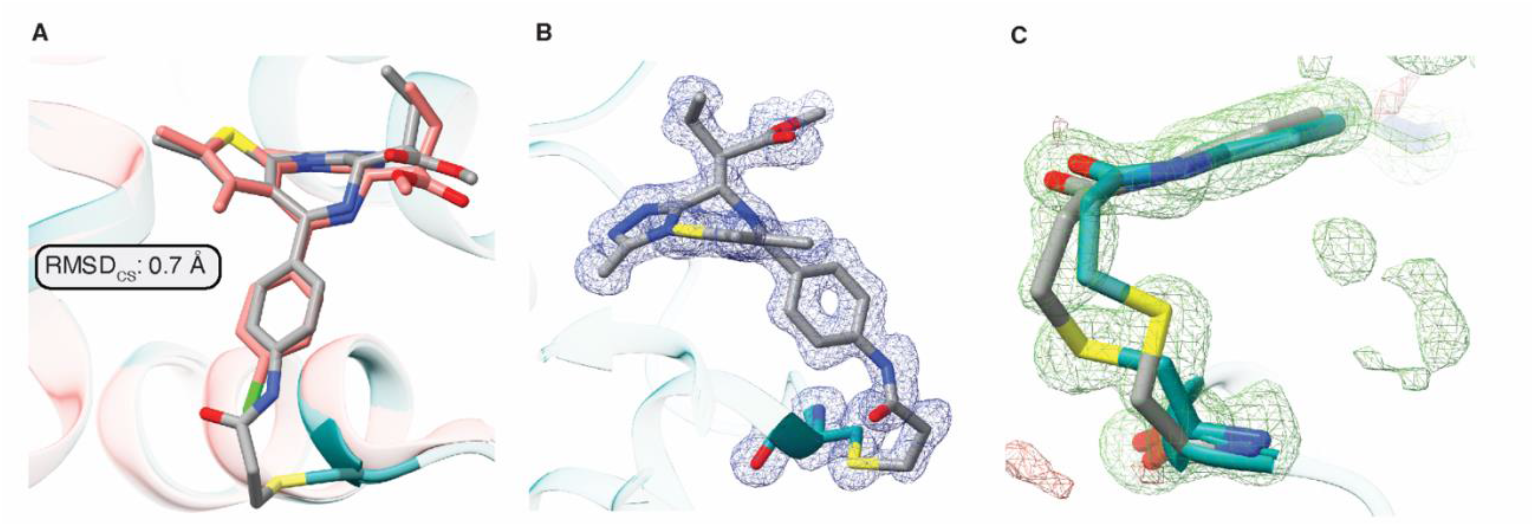
Co-crystal structure of the covalent adduct between MR116 and the bromodomain mutant. **A)** Overlay of the Brd2-BD2^L383V^/ET-JQ1-OMe (6YTM) and the Brd2-BD2^L383A,D434C^/MR116 solved to 1.3 Å of resolution (RMSDcs 0.7 Å). **B)** 2Fo-Fc electron density contoured to 1 sigma around the MR116 ligand and the bonded cysteine in the protein (rotamer 1). C) The polder map (Fo-Fc) around the covalent bond reveals electron density that supports two possible conformations.

Having established and validated the binding mode for the BromoCatch mutant protein, we next moved to assess intracellular target engagement of the BromoCatch tag across our eight most effective covalent compounds using nanoBRET. A BromoCatch-Nanoluciferase (NanoLuc) construct was generated and subsequently transfected into HEK293 cells. We synthesized a reversible tracer molecule, ET-JQ1-PEG3-Bodipy (560/590 nm), which binds to the BromoCatch binding site reversibly and can be competitively displaced by BromoCatch covalent ligands, resulting in a measurable loss of BRET signal. HEK293 cells expressing BromoCatch-NanoLuc were preincubated with ET-JQ1-PEG3-Bodipy (1 µM) before adding the electrophilic compounds. After a 10-minute incubation, the BRET signal was recorded (**Figure 4A**). Among the tested ligands, MR116 exhibited superior intracellular potency, with an IC_50_ of 31 nM, significantly lower than the reversible ligand ET-JQ1-OMe (88 nM) (**Supplementary Figure 5 & Supplementary Table 4**). Given the short incubation time, this result suggests rapid cellular uptake of MR116, reflecting the good cell permeability of the BromoCatch ligand. MR116 also exhibits very rapid covalent binding kinetics, as evidenced by complete conversion to the covalent adduct that was observed by intact MS upon as short as 5 min incubation time at room temperature (**Supplementary Figure 6**). We hypothesized that MR116’s enhanced intracellular activity is due to its covalent binding. To validate this, we next performed a nanoBRET-based residence time experiment. HEK293 cells expressing BromoCatch-NanoLuc were first equilibrated with a saturating concentration of ET-JQ1-OMe (250 nM), MR116 (250 nM) or DMSO for 2 hours. Following this, the ligand was removed, cells were washed in media and ET-JQ1-PEG3-Bodipy tracer (25 µM) was added. An increase in BRET signal would indicate tracer binding to BromoCatch-NanoLuc, occurring only after ligand dissociation, thereby providing a residence time measurement. However, no increase in BRET signal was observed for MR116 over 120 minutes (**Figure 5B**), indicating covalent engagement of MR116 with the BromoCatch tag in live cells.

Having established the best protein-ligand pair, we moved to derivatise the covalent binder MR116 into bifunctional compounds to incorporate additional functionalities. The methyl ester in the JQ1scaffold represents a suitable and widely precedented exit vector to append a diverse range of functional handles.^31,49,50^

We first assessed a common application of SLP’s for click-chemistry based conjugation. We synthesized alkyne based-probe BromoCatch ligand MR155 that can be ‘clicked’ to an azide-bearing molecule using copper catalysed azide-alkyne cycloaddition (CuAAC). Upon incubation of the alkyne MR155 with different protein variants followed by click-reaction the with the sulfonated Cy5 azide, we observed that only BromoCatch but not wild-type or L378A mutant (both lacking the nucleophilic cysteine) could be fluorescently labelled *in vitro* (**Supplementary Figure 7**). Next, we explored the utility for biotin-based protein analysis. ^19,51,52^ To this end, we developed a biotin-functionalised bivalent probe (MR169, Fig 6A). The biotin probe maintained the high stabilising effect for BromoCatch (ΔTm of +20, **Supplementary Figure 8**) and complete covalent modification of the expected Δmass was confirmed by intact protein MS analysis when tested with the recombinant BromoCatch protein (at 2:1 ratio) *in vitro* (Fig 6B). The utility of the probe for protein analysis in cell lysates was evaluated in HEK293-FT cells transfected with a H2B-BromoCatch mammalian expression construct. The H2B-BromoCatch tagged protein was specifically biotinylated by treatment at increasing concentrations of MR169. Western blot analysis shows specific biotinylating of H2B-BromoCatch at all concentrations tested and no cross-biotinylating in WT or in transfected cell lines, even when the probe is at the highest concentrations (**Figure 6C**). The anti-H2B antibody detection demonstrates the successful transfection of the H2B-BromoCatch construct (31 kDa) and the band co-localises with the anti-streptavidin band, confirming the targeted protein identity.

**Figure 6.**
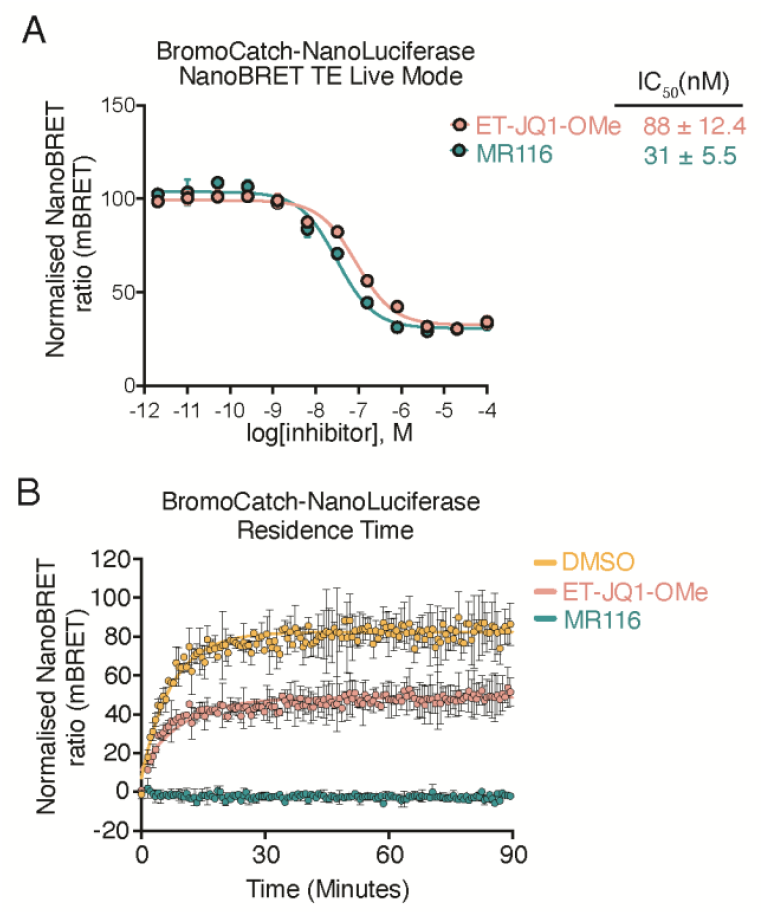
Quantitative analysis and determination of covalency of MR116 ligand binding to NanoLuc-tagged Brd4-BD2^L387A,E438C^ expressed in HEK293 cells. A) Inhibition of the specific binding of 1 µM ET-JQ1-Bodipy to NanoLuc-tagged Brd4-BD2^L387A, E438C^ expressed in HEK293 cells by increasing the concentrations of MR116 and ET-JQ1-OMe. Data are mean ± SE from N4 independent repeats. IC50 values calculated as mean (±S.E.M.) from four independent biological experiments using log(inhibitor) vs. response (three parameters) using Graphpad Prism Version 10.2.3. B) Real-time kinetic analyses of Et-JQ1-Bodipy binding to NanoLuc-tagged Brd4-BD2^L387A,E438C^ expressed in HEK293 cells using 250 nM covalent ligands MR116 and ET-JQ1-OMe. BRET ratios for kinetic studies have been baseline-corrected at time 0. Data are mean ± SD of N2 repeats. Graphpad Prism Version 10.2.3.

Since BromoCatch is directly derived from BromoTag, our inducible degron tag that allows rapid potent and selective degradation of tagged protein upon treatment with degrader AGB1,^31^ we were curious to assess the compatibility of BromoCatch to also be recruited for degradation. To this end, the covalent PROTAC MR170 was synthesized as an AGB1 analogue to investigate the impact of covalent target engagement on PROTAC mediated degradation. MR170 (VHL) is based on MR116 with PEG3 linker-functionalized E3 ligase binders (**Supplementary Figure 9A)**. To test the covalent PROTAC in live cells we generated a HiBiT-BromoCatch-Brd4 HEK293 cell line, and validated it as a heterozygous knock-in for HiBiT-BromoCatch-Brd4 HEK293 via immunoblotting (**Supplementary Figure 9B)**. Following this we subsequently performed a live-cell HiBiT kinetic degradation assay in which MR170 achieving 61% degradation (Dmax) at 2.5 µM over 9 h. In comparison, non-covalent AGB1 PROTAC, originally designed for BromoTag, could degrade >90% BromoCatch-tagged Brd4 as model target at 1 µM over 9 h confirming its effectiveness as a degrader in this system (**Supplementary Figure 9C, Supplementary Table 5**). These findings demonstrate BromoCatch’s compatibility with covalent PROTACs, though full degradation efficiency was not achieved. However, the reversible AGB1 PROTAC retained efficacy in degrading BromoCatch, suggesting the newly introduced Cys mutation does not disrupt the PROTAC mode of action. This means the AGB1 degrader could be readily deployed to rapidly deplete the population of BromoCatch-tagged protein might remain unbound/unreacted in a self-labelling experiment.

One of the most widely used applications of SLP’s is the fluorescent labelling of proteins of interest. To demonstrate the utility of our new system in this context, we built a BromoCatch-specific tetramethylrhodamine (TMR) probe based on the MR116 ligand (**MR202, Figure 6D**). Firstly, the fluorescent probe was tested *in vitro* using purified recombinant protein. When the probe was titrated at increasing concentrations against the protein, maximal fluorescence intensity was achieved at stoichiometric levels of protein to probe (10 µM). The exquisite selectivity for BromoCatch was demonstrated over Brd4-BD2 and Brd4-BD2^L387A^, for which no binding was observed (**Figure 6E**). To validate the utility of the MR202 probe for fluorescent labelling of BromoCatch-tagged proteins in a cellular context, H2B-BromoCatch transiently expressing HEK293-FT cells were treated for 2 h with MR202 (100 nM) and the cell lysates analyzed by western blotting. The membrane was co-stained with anti-H2B antibody (IR800 channel) and tubulin. The transfection was confirmed by the appearance of two H2B bands (H2B^WT^ at 15KDa and H2B-BromoCatch at 30 KDa) in the H2B-BromoCatch transfected cell line lysates, with only the lower molecular weight band observable in the un-transfected U2OS cells. The TAMRA fluorescence was detected in the Alexa 546 channel, the specific bands corresponding to the transfected H2B-BromoCatch protein at all concentrations tested (0.25 µM, 1 µM, 2.5 µM) were observed and the identity of the bands were confirmed by the overlapping bands H2B-BromoCatch.

To assess the compatibility of BromoCatch with live-cell fluorescence-based applications, a fluorogenic BromoCatch probe using Janelia Fluor® 635 (JF635) was prepared (**Figure 7A**).^53^ Janelia Fluor dyes are rhodamine-based dyes with excellent photophysical properties including high brightness and photostability. These dyes are small, cell permeable thanks to their lactone-zwitterion K_L-Z_ equilibrium, and have excellent environmental fluorogenic properties, enabling no-wash, live-cell imaging applications. For example, JF635 has been widely used in conjunction with other SLP systems such as Halo- and SNAP-tag ^5,54,55^. It is particularly suited to these applications due to its red-shifted emission (away from cellular autofluorescence and enabling less destructive, higher wavelength activation) and preferential K_L-Z_ providing a high degree of fluorogenicity. To develop a JF635 based fluorescent probe, we exchanged the 5-TMR fluorophore in **MR202**, while retaining the original linker (**Figure 7B**).

**Figure 7.**
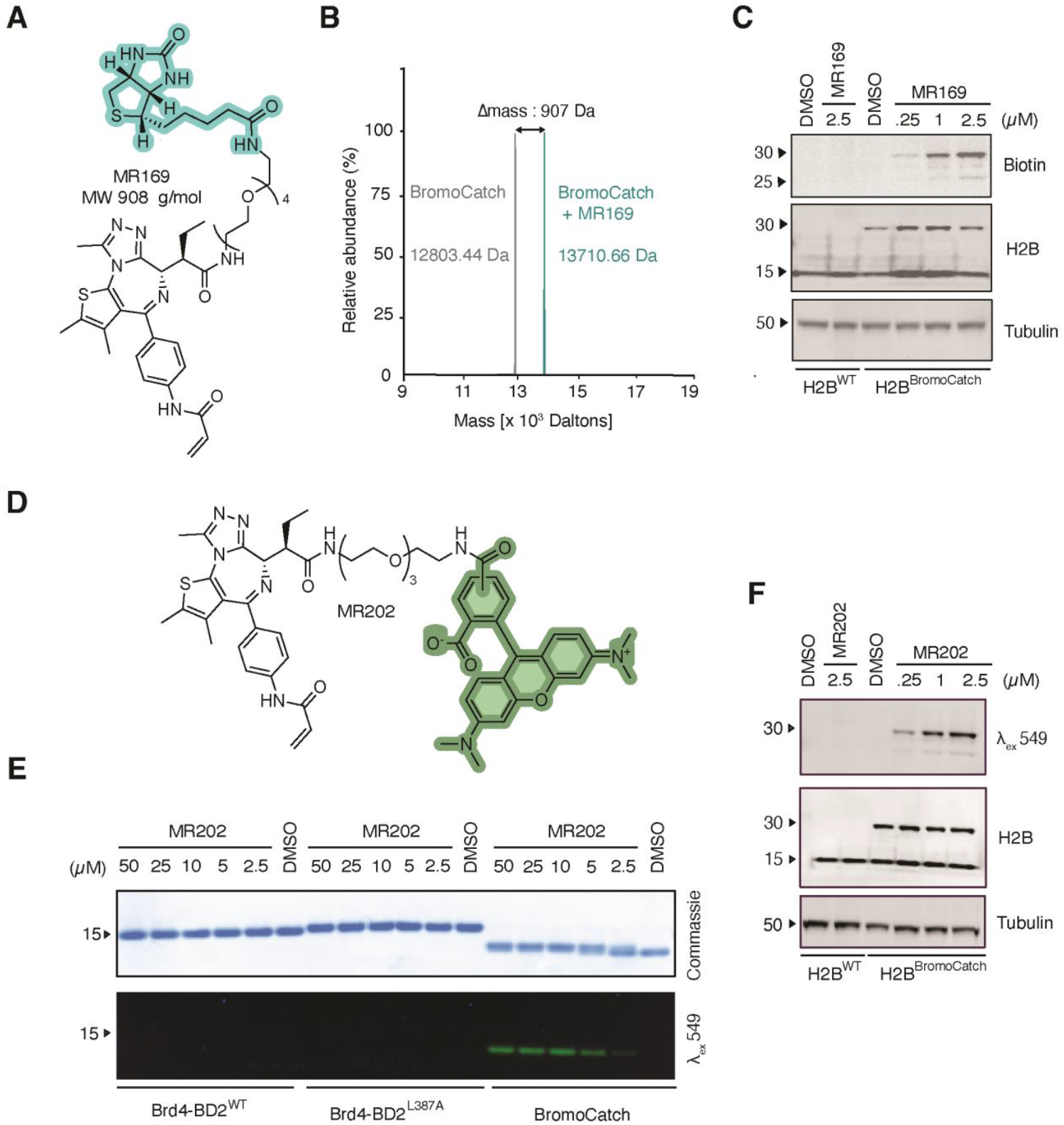
Biotin and TAMRA probes for cell lysate experiments. A) Chemical structure of biotinylated probe MR169. B) Formation of covalent adduct BromoCatch:MR169 evidenced by intact protein MS. C) Cell lysate experiments show that MR169 specifically detects H2B-BromoCatch at 0.25 µM concentration of probe. The probe showed no unspecific binding in HEK293FT WT cells when incubated at up to 2.5 µM. D) Chemical structure of TAMRA fluorescent probe MR202. E) In-gel assay titration of the probe with purified proteins BromoCatch, Brd4-BD2^WT^, or Brd4-BD2^L387A^ and HEK293FT H2B-BromoCatch cell lysate experiment.F) MR202 specifically detects H2B-BromoCatch at 0.25 µM concentration of probe. The probe showed no unspecific binding in HEK293FT WT cells when incubated at up to 2.5 µM.

The JF635 probe C10852S was titrated against the recombinant BromoCatch protein (10 µM). Conveniently, the JF635 fluorophore can be environmentally “switched on” in presence of SDS micelles, an extremely useful feature when it comes to in-gel protein analysis, where the probe’s fluorescence is fully switched on, behaving as an ‘always on’ fluorophore. This feature allowed us to evaluate the concentration dependent increase ‘in-gel’. The fluorescence when titrating the probe showed the same concentration dependent response to that of MR202. These results confirm the high affinity and specificity if the probe is maintained despite the change of fluorophore (**Figure 7C**).

The fluorogenic character of C10852S was evaluated by measuring the fluorescence signal in the presence or absence of BromoCatch, Brd4-BD2^L387A^, or Brd4-BD2^WT^ (10 µM). The probe showed a strong ‘switch on’ with BromoCatch and low background fluorescence in buffer. Fluorogenicity was specific to BromoCatch, with no unspecific signal observed with Brd4-BD2^WT^. Interestingly, C10852S also produced a switch-on effect with Brd4-BD2^L387A^, indicating efficient reversible binding.^56^ However, fluorescence levels were lower compared to the covalent system, suggesting covalency enhances fluorogenicity. SDS gel analysis showed maximal fluorescence at stoichiometric protein-to-probe levels (10 µM), after which the signal plateaued despite increasing C10852S concentration. Notably, the fluorescence signal at 10 µM in the positive control was slightly lower than with the cysteine mutant, possibly due to poor solubility or protein binding (**Figure 7D,E, Supplementary Figure 10**).

Recognizing the similarity in chemical structure between our BromoCatch covalent binder MR116 and JQ1-based molecular glues that induce DCAF16-dependent degradation of endogenous Brd4 protein via a template-assisted covalent cross-reactivity towards DCAF16 ^41,42^, we assessed any undesired on-target DCAF16-mediated degradation of BromoCatch by our C10852S probe compound. Using HiBiT-tagged endogenous Brd4, Brd3, and Brd2 HEK293 cells, along with a BromoCatch-Brd4 HEK293 model, we performed HiBiT-based degradation assays. While MZ1 and AGB1 induced BromoCatch degradation (DC_50_[4H] = 159 and 21.3 nM) C10852S showed no degradation at 4 h in both the BromoCatch and the wild-type BET cell lines, confirming its incompatibility with the molecular glue mechanism (**Supplementary Figure 11, Supplementary Table 6**). We hypothesize that a potential DCAF16-mediated induced degradation is prevented due to a combination of at least two of our design features (i) the ‘bump’, substantially weakening engagement to wild-type bromodomains and (ii) the functionalization of the ligand via the carboxylic acid vector, which would disrupt the molecular glue ternary complex (ref. ^35,41,42^).

Finally, to validate the applicability of BromoCatch for fluorescent labeling of tagged proteins in live cells, we observed the fluorescence ‘switch-on’ of the Janelia Fluor 635-conjugated C10852S probe using live-cell confocal microscopy. For this, we employed the previously described H2B-BromoCatch mammalian expression construct, transiently transfecting both U2-O S and HEK293FT cells. Post-transfection, cells were incubated with C10852S (100 nM), and fluorescence was monitored via confocal microscopy. In U2-O S cells, fluorescence activation was detected within 1 hour and peaking at 8 hours, whereas in HEK293FT cells, ‘switch-on’ was observed from 1 hour and peaking at 6 hours. Fluorescent localization was strictly confined to the nucleus, as confirmed by Hoechst 33342 nuclear counterstaining. Furthermore, the fluorogenic switch on of the probe was effectively prevented upon treatment with a saturating concentration (25 µM) of the bromodomain inhibitor ET-JQ1-OMe (**Figure 9, Supplementary Figures 12-14**). Notably, fluorescence activation of Janalia Fluor 635® was not observed in non-transfected HEK293FT or U2-O S cells, indicating that JF635 fluorescence occurs exclusively upon engagement of C10852S with the H2B-BromoCatch fusion protein. This result strongly qualifies the probes selectivity for the BromoCatch-tagged target and highlights its utility in live-cell imaging applications.

**Figure 8.**
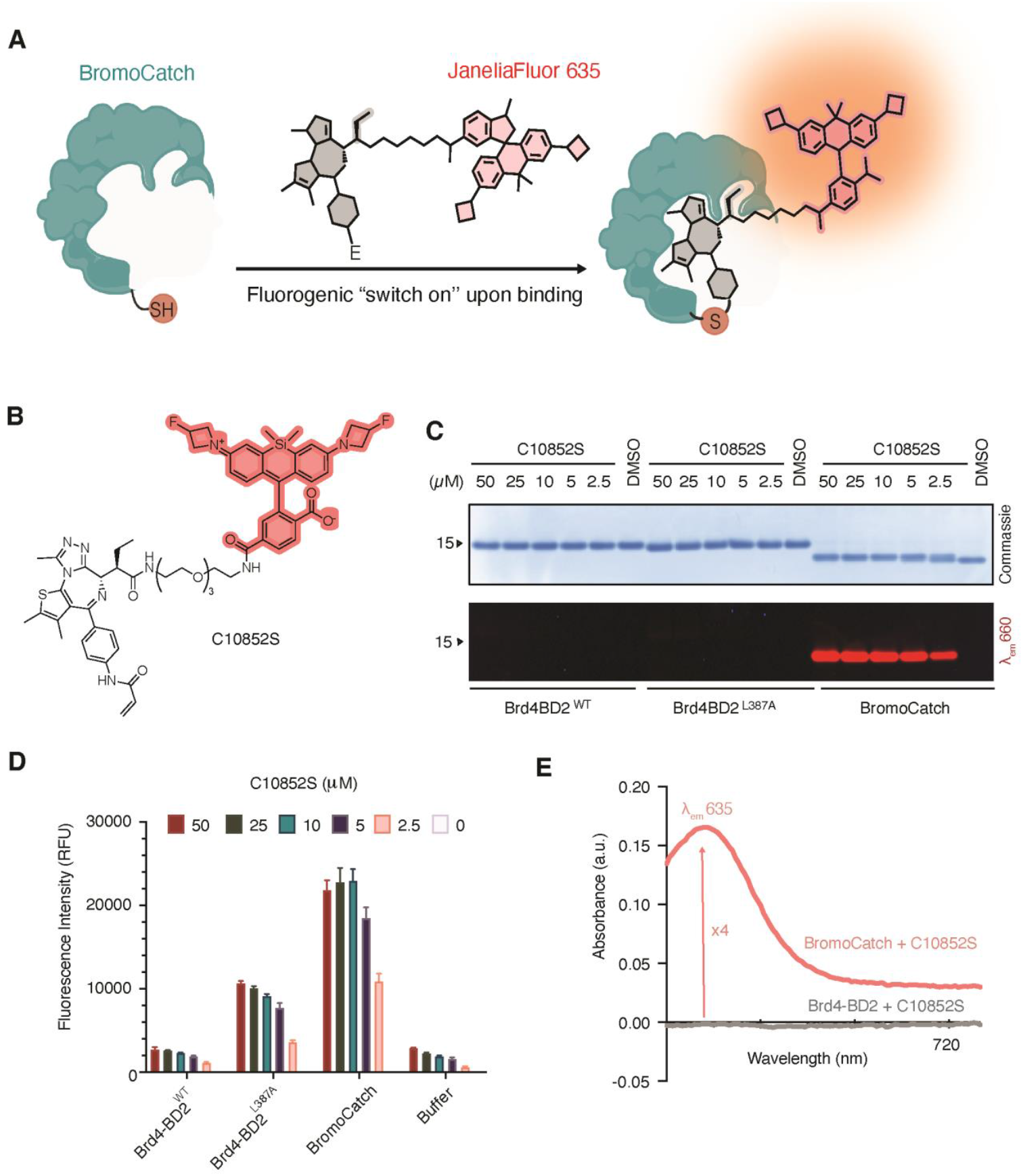
In vitro validation of BromoCatch covalent ligand MR116 functionalized with Janelia Fluor® 635. A) Switch on principle of Janelia Fluor® 635 BromoCatch probe. B) Structure of C10852S probe. C) In gel assay titrating the different proteins with increasing concentrations of the C10852S probe. D) Fluorogenic switch on in presence of the different proteins with increasing concentrations of the C10852S probe. E) Specificity of fluorogenic ‘switch on’ measured by absorbance in presence of BromoCatch or Brd4-BD2^WT^.

**Figure 9.**
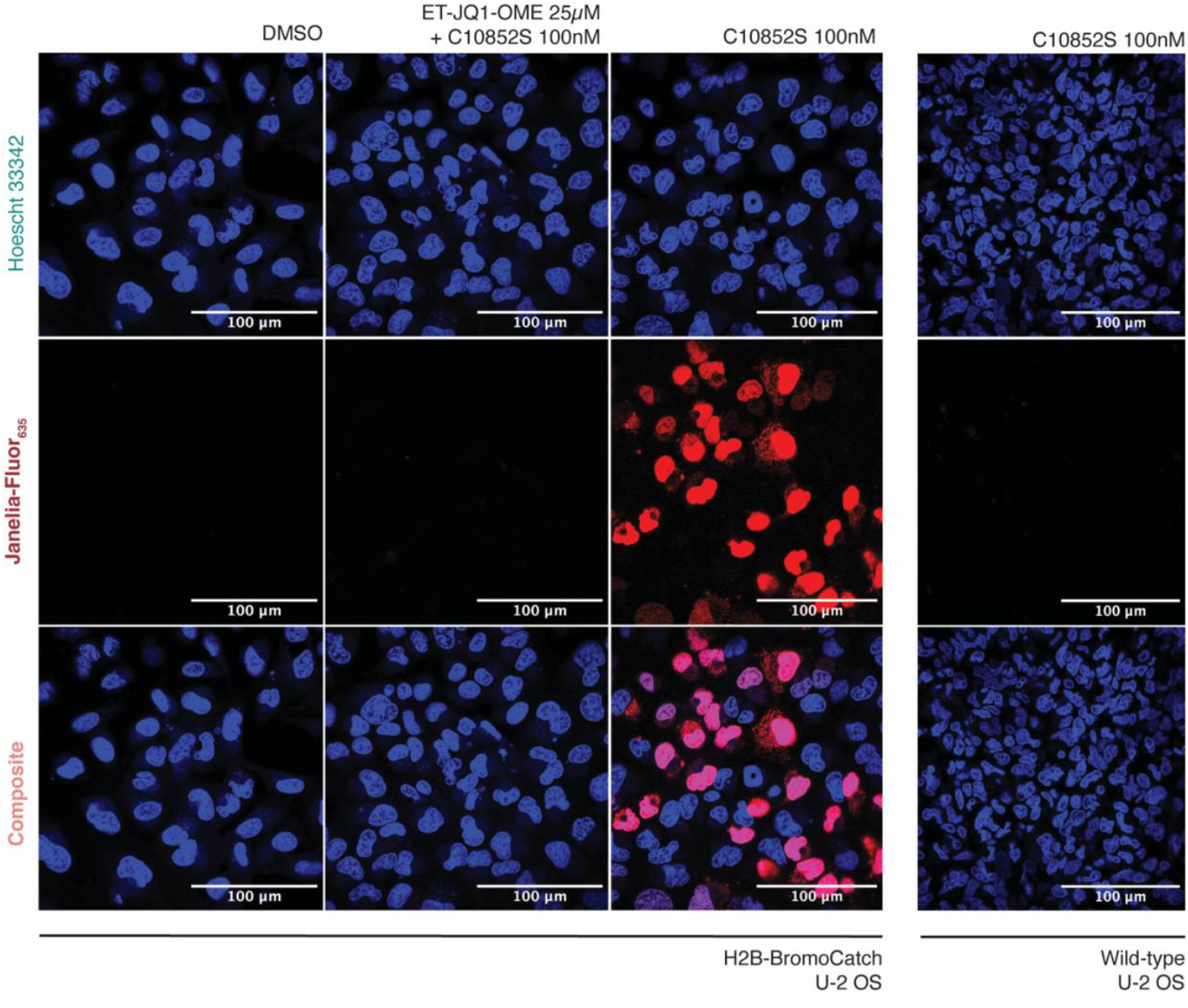
Cellular validation of the functionalized MR116 Janelia Fluor 635 covalent probe (C10852S) using live-cell confocal microscopy. Live-cell confocal imaging of H2B-BromoCatch transiently transfected U2-O S cells treated with C10852S. Cells were incubated in DMSO, 100nm C10852S or a 100nM C10852S cotreatment with 25µM ET-JQ1-OMe for 8 hours prior to imaging. Hoechst 33342 nuclear counterstain and JF 635 fluorescence detected. N=2 independent repeats.

## Conclusion and future perspective

We present BromoCatch, a self-labeling protein tag (SLP) platform for protein analysis and live cell imaging applications. Leveraging our ‘bump and hole’ BromoTag system, we engineered a cysteine into the bromodomain ligand binding pocket, creating the BromoCatch double-mutant protein BRD4-BD2 ^L387A,E438C^. The BromoCatch covalent ligand MR116 was designed by introducing an acrylamide electrophilic group at the *para* position of the pendant aryl ring and proved to be the most effective covalent binder. Functionalisation of MR116 allowed the synthesis of a panel of bivalent compounds for diverse applications including biotin-based probe MR169 for biotin-based detection, and “always on” TMR probe MR202 for fluorescence-based detection of BromoCatch in cell lysates. We additionally synthesised an environmental “switch-on” probe, C10852S, which featured selective binding of a BromoCatch tagged protein *in cells*, and excellent switch-on activation with minimal background fluorescence, demonstrating utility and suitability for live-cell imaging. Together, our data establishes BromoCatch as a versatile tool for protein labeling and live-cell imaging, with the potential for broader applications in protein manipulation and for multiplexing using orthogonal SLT platforms. The small size of the tag minimizes its impact on cellular proteins, enhancing its utility for various experimental needs. Future work will explore further applications of probes for this system utilizing different fluorophores, as well as further protein engineering of the BromoCatch tag for multiplexing purposes – like the CLIP-Tag system. It will be warranted to explore the suitability of BromoCatch as a tag to be fused in protein loops^57^ other than at the N- or C-terminus as these often deactivate the endogenous protein function. Future probe conjugates could be designed to bear other functions and activities beyond those exemplified here. We envisage these applications to include assay development and enablement of induced-proximity modalities beyond targeted protein degradation, for example in combination with existing orthogonal SLPs and non-covalent tags;^58,59^ novel labeling approaches to aid protein detection in cellular structural biology approaches using Cryo-Electron Tomography (cryo-ET) and Cryo correlative light and electron microscopy (CLEM); ^60,61^and new labels for in vivo applications.^62^ We anticipate that our Bromocatch system’s favorable properties demonstrated herein will underpin establishment as a broadly used tag for protein labeling and chemical manipulation of biological systems.

## ASSOCIATED CONTENT

Supporting Information is available free of charge and includes: Illustration of Brd2-BD2 and Brd4-BD2 homology; full dataset for docking scores and RMSD; protein mutant sequences, illustration, and characterization; complete differential scanning fluorimetry data and INTACT MS covalent modification percentages as represented in heatmaps in Fig. 3 and 4 in the main manuscript; additional crystallography data of the MR116/Brd3-BD2 L383A D434C co-crystal; full data on NanoBRET assays; BromoCatch click experiment data; MR169 covalent modification validation by INTACT MS and DSF; additional plots on the fluorogenicity of C18052S; extended confocal live-cell imaging including time-dependent reads; covalent PROTAC MR170 and AGB1 degradation experiments; HiBiT lytic degradation assays. Additional, the supporting information includes -general experimental details and compound characterization.

## Accession Codes

Atomic coordinates have been deposited in the Protein Data Bank and will be released upon article publication. Accession code of Brd2-BD2^L383A,D434C^ in complex with compound MR116 is PDB 9QRK.

## Author Contributions

M.R.R. and C.G.C. designed and conducted experiments. M.R.R., G.P.M., and A.C. designed compounds. A.G.B. established synthetic routes to bumped ligands and contributed to initial compound design. M.R.R., A.K.E., M.C.N., R.E.A., P.M.W., S.J.R. synthesized compounds. M.R.R. designed and performed computational docking calculations, biophysical assays, intact LC-MS, in vitro fluorescent-based assay, and cell lysate assays. C.G.C. performed biological assays in live cells and designed and developed the genetically engineered cell lines used in paper. M.A.N., M.R.R. and M.D. expressed and purified recombinant proteins for biophysical and structural studies. M.A.N. crystallized the protein-ligand adduct, collected and processed the X-ray data and M.R.R. performed molecular replacement, structure refinement and data deposition. J.O.C.-B., G.P.M., and H.J.M. provided input to project design. A.C. conceived the idea, acquired research funds, and supervised the project. M.R.R., C.G.C., G.P.M., H.J.M. and A.C. wrote the manuscript with input from all co-authors. All authors have reviewed and approved the final version of the manuscript.

## Notes

The authors declare the following competing financial interest(s): A.C. is a scientific founder and shareholder of Amphista Therapeutics, a company that is developing targeted protein degradation therapeutic platforms. A.C. is on the Scientific Advisory Board of ProtOS Bio. The Ciulli laboratory receives or has received sponsored research support from Almirall, Amgen, Amphista Therapeutics, Boehringer Ingelheim, Eisai, Merck KGaA, Nurix Therapeutics, Ono Pharmaceutical and Tocris(Bio-Techne). A.K.E., M.C.N., R.E.A., P.M.W., S.J.R., J.O.C.-B., G.P.M., and H.J.M. are employees of Tocris(Bio-Techne).

## ACKNOWLEDGEMENTS

The authors thank Dundee University’s imaging facility for provision of the Zeiss 880 Airyscan Confocal Microscope as well as the flow cytometry and cell sorting facility for the use of their SH800 cell sorter and expertise; Satpal Virdee and Mathieu Soetens (MRC-PPU) for providing access and assistance with the Agilent system for intact ESI-MS; and Alena Kroupova and Ryan Casement (Ciulli Lab, CeTPD) for technical support with solving the co-crystal structure. We acknowledge Diamond Light Source, United Kingdom for provision of synchrotron radiation facilities (proposal mx35324-32), and we would like to thank the staff of MX beamline I24 for assistance and support with data collection.

