## Supplementary Materials for "BromoCatch: a self-labelling tag platform for protein analysis and live cell imaging"

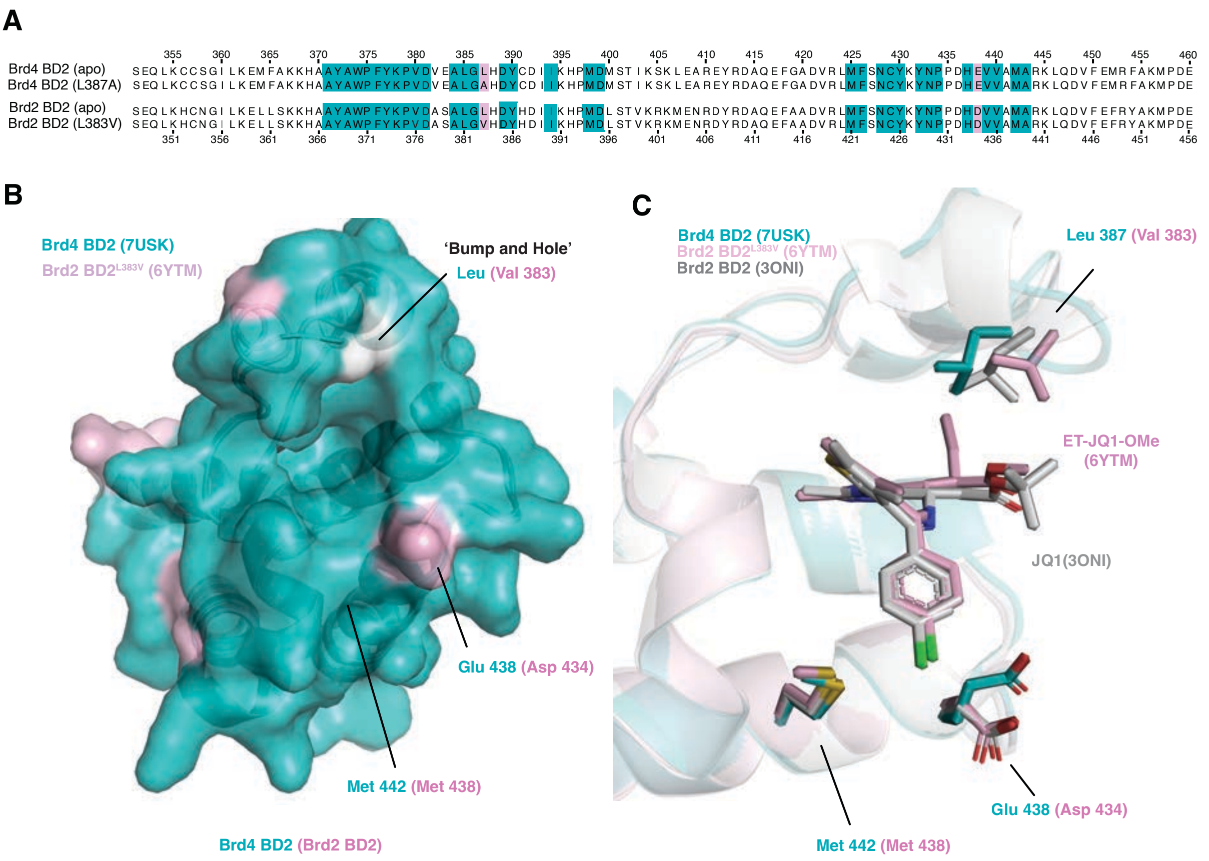


**Supplementary Figure 1. Brd2-BD2 / Brd4-BD2 sequence homology.** *In silico* studies were done using the Brd2-BD2$^{L383V}$ /ET-JQ1-OMe (6YTM) crystal as reference. A) Sequence alignment shows the homology of the JQ1 binding pocket in both the apo Brd4-BD2 and Brd2-BD2 and the corresponding 'bump and hole' mutants Brd2-BD2$^{L383V}$ and3Brd4-BD2$^{L387A}$. Highlighted in blue or pink are residues that are within the bromodomain binding pocket (within 15 Å from the ligand). B) Surface of Brd4-BD2 / Brd4-BD2 crystals (7USK/6YTM) alignment, highlighting where residues are different in pink. Leucine 387 in apo Brd4-BD2 (Leu 383 in apo Brd2-BD2) has the functional 'hole' in the 6YTM crystal with the mutation L383V (functionally silent L387A mutation in BromoCatch). C) Brd4-BD2 / Brd4-BD2 alignment to show the side chains of amino acids that have been chosen for engineering BromoCatch.

**Supplementary Table 1**. **Reversible and Covalent docking results for both cysteine-containing mutants.**

| | | | Brd4BD2$^{L387A,M442C}$ | | | | Brd4BD2$^{L387A,E438C}$ | | | |
|---|---|---|---|---|---|---|---|---|---|---|
| "B&H" residue | | | L387A (L383V) | | | | L387A (L383V) | | | |
| "Trap" residue | | | M442C (M438C) | | | | E438C (D434C) | | | |
| Docking type | | | Reversible | | Covalent | | Reversible | | Covalent | |
| | | | Glide score (kcal/mol) | RMSD (Å) | Cov score (kcal/mol) | RMSD (Å) | Glide score (kcal/mol) | RMSD (Å) | Cov score (kcal/mol) | RMSD (Å) |
| 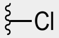 | **ET-JQ1** | *para* | -7.38 | 0 | N/A | N/A | -8.12 | 0 | N/A | N/A |
| 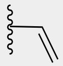 | **MR100** | *para* | -7.38 | 0.03 | N/R | N/A | -7.85 | 0.15 | N/R | N/A |
| | **MR108** | *meta* | -7.76 | 0.14 | N/R | N/A | -7.8 | 0.04 | N/R | N/A |
| 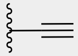 | **1** | *para* | -7.41 | 0.04 | -5.54 | 1.592 | -8.14 | 0.04 | -7.546 | 0.09 |
| | **2** | *meta* | -7.94 | 0.1 | -7.4 | 0.299 | -7.55 | 0.74 | -7.006 | 0.39 |
| 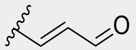 | **3** | *para* | -7.40 | 0.7 | -4.79 | 3.428 | -7.78 | 0.17 | -7.552 | 0.13 |
| | **4** | *meta* | -6.03 | 2.87 | -5.91 | 3.316 | -6.16 | 2.97 | -7.19 | 0.21 |
| 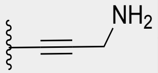 | **MR101** | *para* | -8.29 | 0.73 | -7.97 | 0.421 | -8 | 0.04 | -7.36 | 0.09 |
| | **MR109** | *meta* | -6.72 | 2.93 | -7.01 | 1.687 | -6.59 | 3.02 | -7.18 | 0.15 |
| 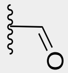 | **MR70** | *para* | -7.52 | 0.04 | -4.31 | 3.139 | -7.86 | 0.75 | -7.45 | 0.15 |
| | **MR115** | *meta* | -7.77 | 0.2 | -7.28 | 0.473 | -7.68 | 0.32 | -7.32 | 0.2 |
| 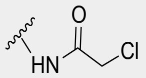 | **MR112** | *para* | -7.50 | 0.08 | -5.82 | 0.791 | -7.77 | 0.04 | -7.17 | 0.09 |
| | **MR118** | *meta* | -7.78 | 0.09 | -7.68 | 0.119 | -7.62 | 0.06 | -7.23 | 0.16 |
| 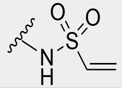 | **MR121** | *para* | -7.31 | 0.1 | -7.53 | 0.148 | -7.35 | 0.08 | -7.14 | 0.08 |
| | **MR117** | *meta* | -7.05 | 0.06 | -7.39 | 0.147 | -7.48 | 0.07 | -7.18 | 0.12 |
| 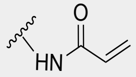 | **MR116** | *para* | -5.00 | 3.09 | -6.68 | 0.468 | -6.1 | 3.03 | -7.29 | 0.09 |
| | **MR119** | *meta* | -8.27 | 0.34 | -8.2 | 0.355 | -7.8 | 0.34 | -6.67 | 1.2 |
| 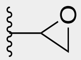 | **MR104** | *para* | -6.31 | 2.92 | -4.14 | 1.60 | -7.37 | 3.41 | -7.4 | 1.52 |
| | **MR111** | *meta* | -7.99 | 0.09 | -7.65 | 0.414 | -7.59 | 0.08 | -7.31 | 0.21 |

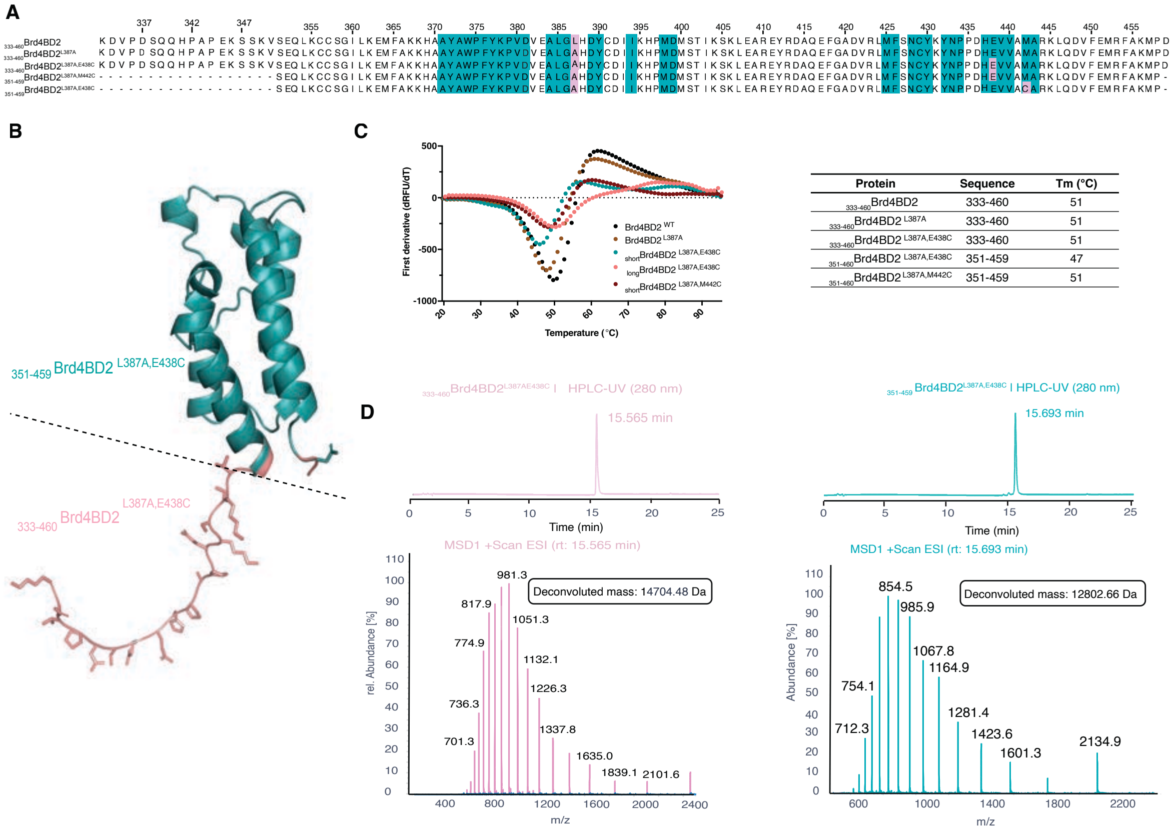


**Supplementary Figure 2. Protein sequences / summary stability / gel.** A) Sequence alignment of the relevant region of Brd4-BD2 where the corresponding mutations are highlighted. B) aligment of Brd4-BD2 for the long and the short version illustrating the cleavage site. C) Melting temperature (Tm) for the different cysteine containing mutants of Brd4-BD2 was determined by nanoDSF. The data demonstrates that the protein mutants developed have similar stability to wild-type Brd4-BD2 (Brd4-BD2 WT). Interestingly, there was a decrease of 4 °C for the short mutant Brd4-BD2$^{L387A,E438C}$ suggesting that the E438C mutation had a slight detrimental effect on stability. The longer version of Brd4-BD2$^{L387A,E438C}$ was expressed to understand the effect of the elongation and it seemed to compensate that difference and 'recover' the stability. D) purity of proteins and corresponding observed mass by INTACT MS for the Brd4-BD2$^{L387A,E438C}$ mutants.

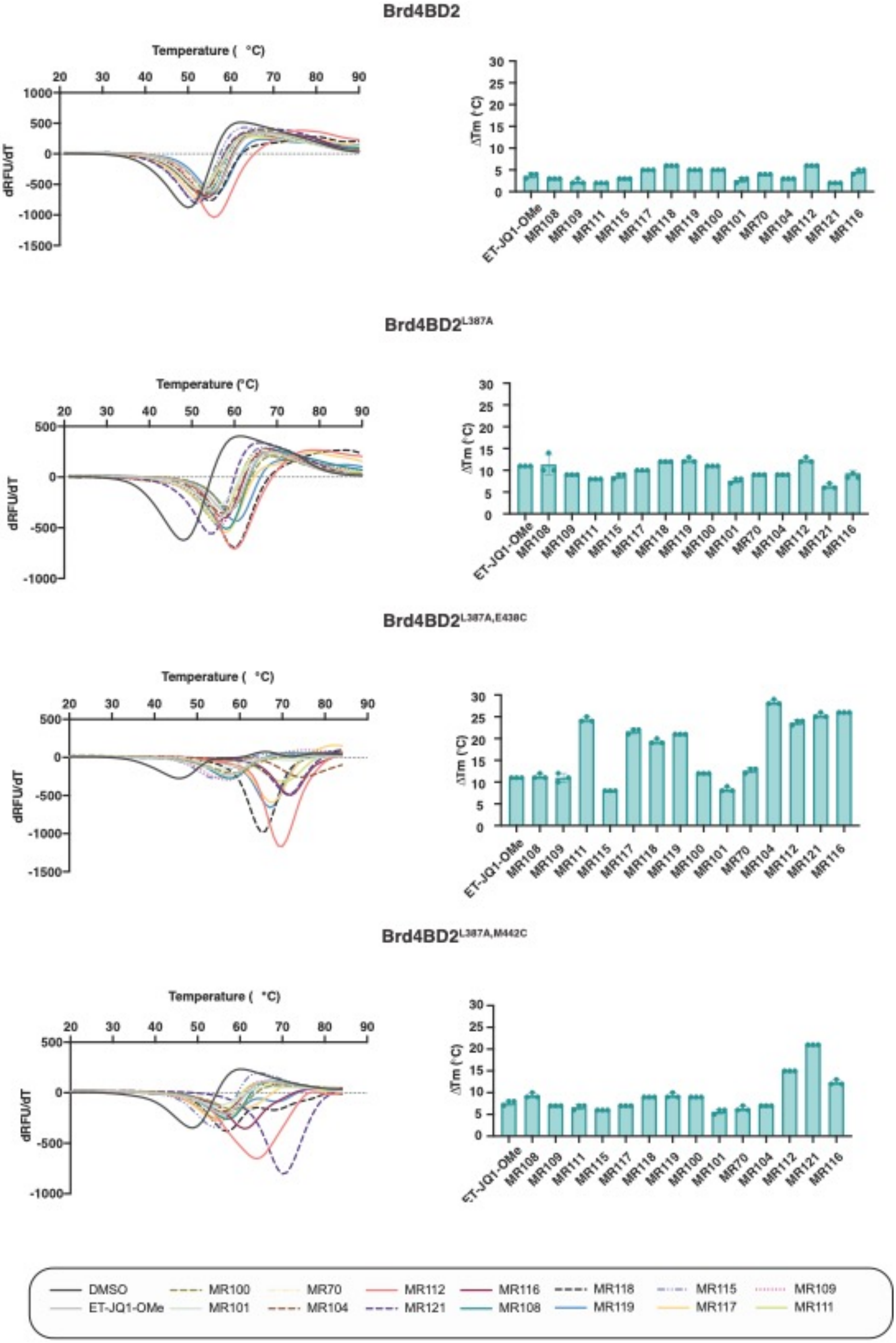


**Supplementary Figure 3. Differential scanning fluorimetry studies.** Left: First derivative melting curves for each protein mutant co-incubated with the compounds at room temperature for 2 hours at a 5:1 ratio (L:P). Right: Corresponding increment in Tm for each of the ligands tested (°C)

**Supplementary Table 2. Systematic studies to evaluate the covalency of the ligands for the different proteins.** Values are % modification of the protein in presence of the ligand based on the UV absorbance and the mass increase.

| Brd4-BD2 protein | ET-JQ1 | MR100 | MR108 | MR101 | MR109 | MR70 | MR115 | MR104 | MR111 | MR112 | MR118 | MR121 | MR117 | MR116 | MR119 |
|---|---|---|---|---|---|---|---|---|---|---|---|---|---|---|---|
| WT | 0 | 0 | 0 | 0 | 0 | 0 | 0 | 5 | 0 | 0 | 0 | 0 | 5 | 0 | 0 |
| L387A | 0 | 0 | 0 | 0 | 0 | 0 | 0 | 0 | 0 | 0 | 0 | 0 | 10 | 0 | 0 |
| L387A, M442C | 0 | 0 | 0 | 0 | 0 | 0 | 0 | 0 | 0 | 10 | 0 | 100 | 67 | 50 | 35 |
| L387A, E438C | 0 | 0 | 0 | 0 | 0 | 0 | 0 | 100 | 100 | 100 | 100 | 100 | 100 | 100 | 100 |

**Supplementary Table 3. Systematic studies to evaluate the stabilization of the different proteins by the different ligands.** The $\Delta T_m$ (°C) were calculated in reference to the corresponding apo proteins. The melting temperatures for the apo proteins were: Brd4-BD2 (apo) Tm = 53 °C; Brd4-BD2$^{L387A}$ (apo) Tm = 51°C; Brd4-BD2$^{L387A,E438C}$ (apo) Tm = 49.3 °C; Brd4-BD2$^{L387A,M442C}$ (apo) Tm = 51°C.

| Brd4-BD2 protein | ET-JQ1 | MR100 | MR108 | MR101 | MR109 | MR70 | MR115 | MR104 | MR111 | MR112 | MR118 | MR121 | MR117 | MR116 | MR119 |
|---|---|---|---|---|---|---|---|---|---|---|---|---|---|---|---|
| WT | 5.0 ± 0 | 5.0 ± 0 | 3.7 ± 0.6 | 2.7 ± 0.6 | 3.0 ±0 | 4.0 ± 0 | 2.0 ± 2 | 3.0 ± 0 | 2.3 ± 0.6 | 6.0 ± 0 | 5.0 ± 0 | 2.0 ± 0 | 3.0 ± 0 | 4.7 ± 0.6 | 6.0 ± 0 |
| L387A | 10 ± 0 | 11 ± 0 | 10 ± 0 | 7.7 ± 0.6 | 9 ± 0 | 9 ± 0 | 8.7 ± 0.6 | 9 ± 0 | 8 ± 0 | 12.3 ± 0.6 | 12 ± 0 | 6.3 ± 0.6 | 10 ± 0 | 9.3 ± 0.6 | 12.3 ± 0.6 |
| L387A, M442C | 7.3 ± 0.6 | 9 ± 0 | 9.3 ± 0.6 | 5.7 ± 0.6 | 7 ± 0 | 6.3 ± 0.6 | 6.0 ± 0 | 7 ± 0 | 6.7 ± 0.6 | 15 ± 0 | 9 ± 0 | 21 ± 0 | 7 ± 0 | 12.3 ± 0.6 | 9.3 ± 0 |
| L387A, E438C | 11.6 ± 0.6 | 12 ± 0 | 11.3 ± 0.6 | 8.3 ± 0.6 | 11 ± 1.0 | 12.7± 0.6 | 8 ± 0 | 28.3 ± 0.6 | 24.3 ± 0.6 | 23.7 ± 0.6 | 19.3 ± 0.6 | 25.3 ± 0.6 | 21.7 ± 0.6 | 26 ± 0 | 21 ± 0 |

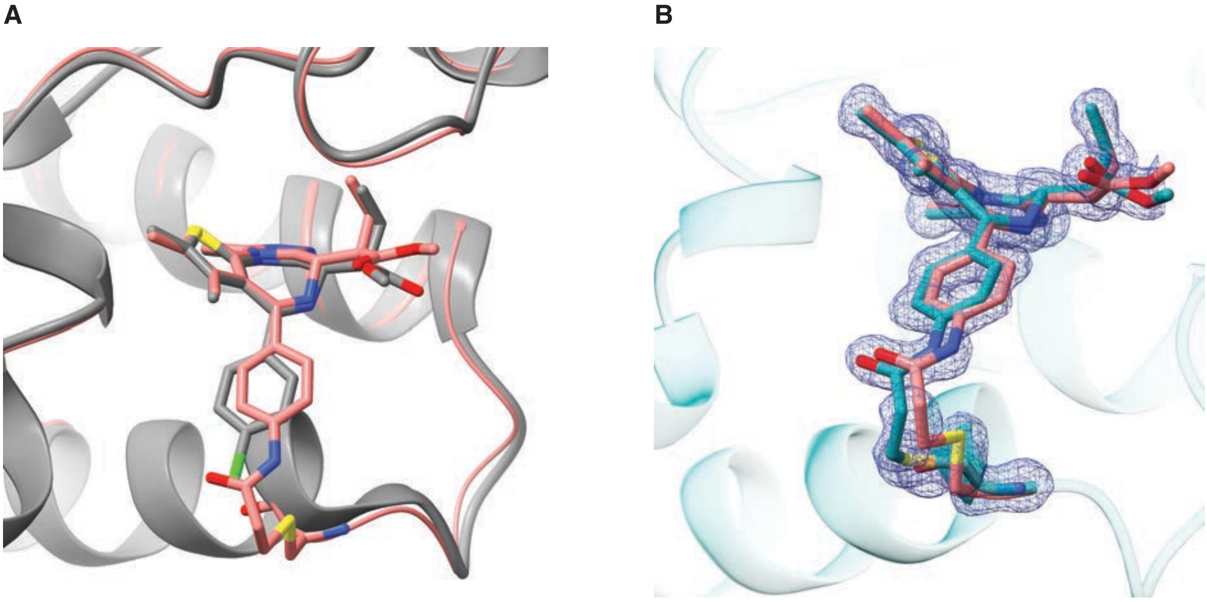


**Supplementary Figure 4. Binding mode and electron density of covalent adduct.** A) Overlay of the Brd2-BD2$^{L383V}$/ ET-JQ1-OMe (6YTM) and rotamer 2 of the Brd2-BD2$^{L383A,D434C}$/MR116 co-crystal, where the aromatic ring is slightly more shifted than rotamer 1 (RMSD$_{CS}$ 1.13 Å) B) 1 sigma 2Fo-Fc electron density around the two conformations of the bonded MR116 ligand.

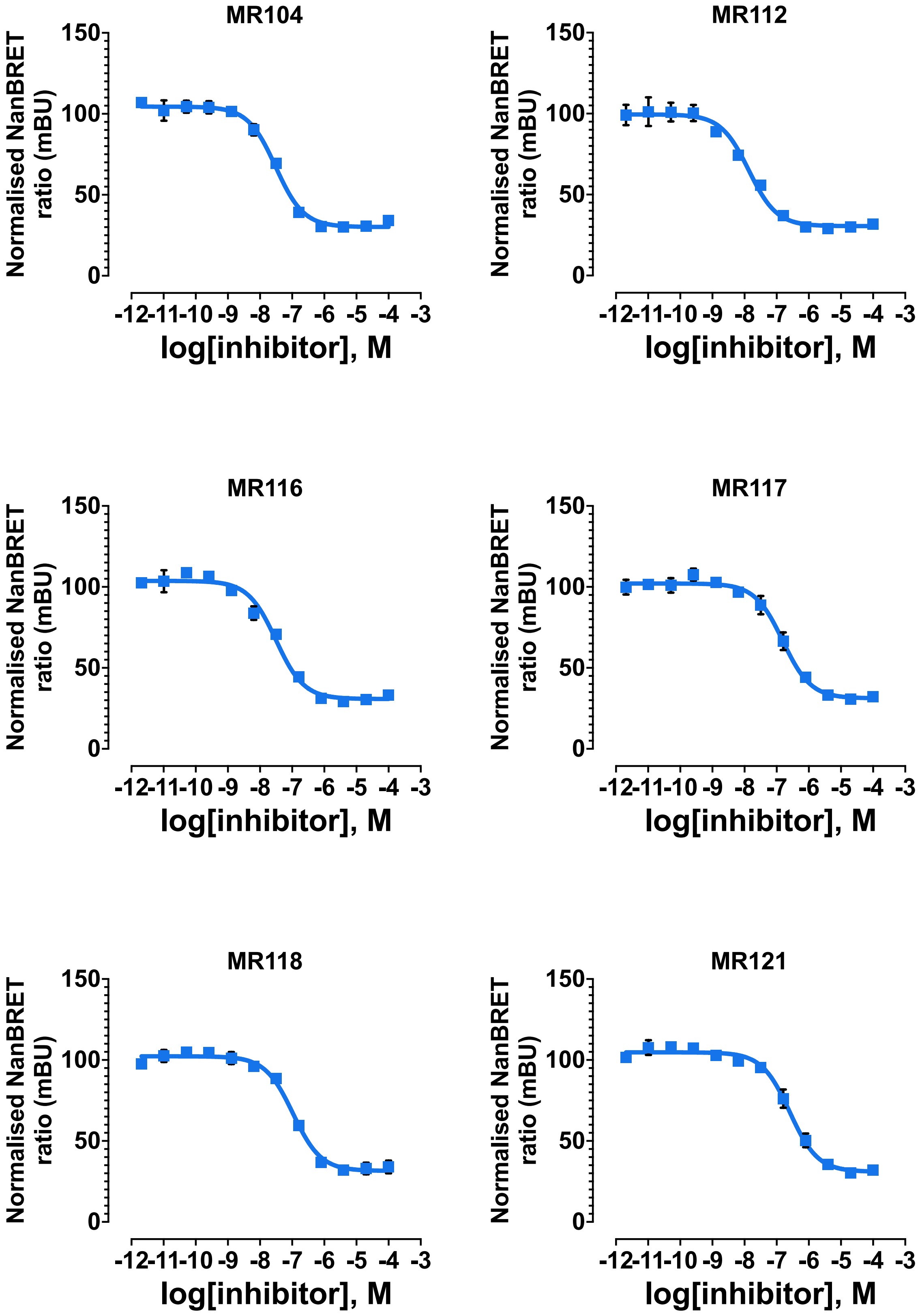

MR104
Normalised NanBRET ratio (mBU)
150
100
50
0
-12 -11 -10 -9 -8 -7 -6 -5 -4 -3
log[inhibitor], M
MR112
Normalised NanBRET ratio (mBU)
150
100
50
0
-12 -11 -10 -9 -8 -7 -6 -5 -4 -3
log[inhibitor], M
MR116
Normalised NanBRET ratio (mBU)
150
100
50
0
-12 -11 -10 -9 -8 -7 -6 -5 -4 -3
log[inhibitor], M
MR117
Normalised NanBRET ratio (mBU)
150
100
50
0
-12 -11 -10 -9 -8 -7 -6 -5 -4 -3
log[inhibitor], M
MR118
Normalised NanBRET ratio (mBU)
150
100
50
0
-12 -11 -10 -9 -8 -7 -6 -5 -4 -3
log[inhibitor], M
MR121
Normalised NanBRET ratio (mBU)
150
100
50
0
-12 -11 -10 -9 -8 -7 -6 -5 -4 -3
log[inhibitor], M

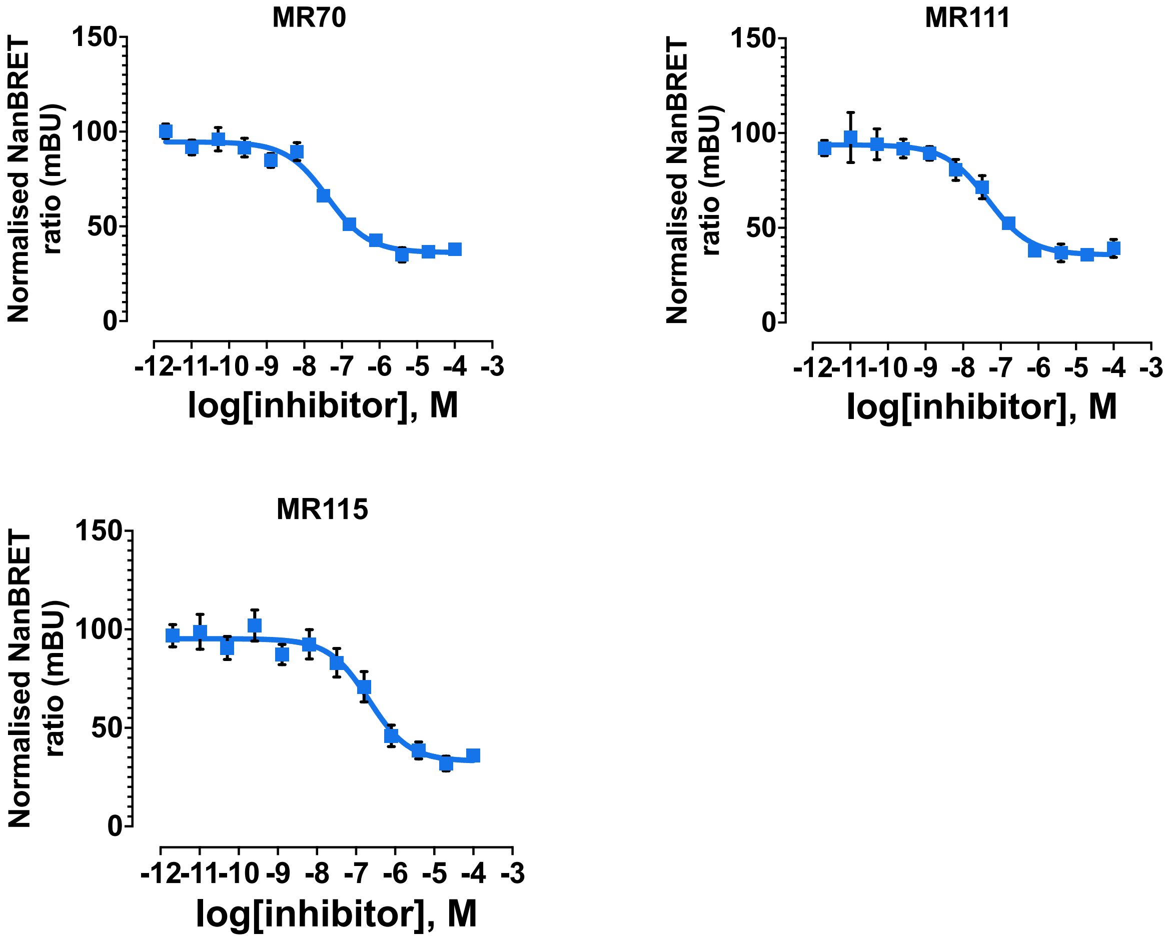


**Supplementary Figure 5. NanoBRET whole plots.** Inhibition of the specific binding of 1 µM ET-JQ1-PEG3-Bodipy to NanoLuc-tagged Brd4-BD2$^{L387A, E438C}$ expressed in HEK293 cells by increasing the concentrations of electrophilic warheads and ET-JQ1-OMe. Data are mean ± SE from N4 independent repeats.

| Compound | $IC_{50}$ (nM) | S.E.M (nM) |
|---|---|---|
| **ET-JQ1-Ome** | 88.2 | 12.4 |
| **MR70** | 40.2 | 11.7 |
| **MR104** | 30.7 | 5.3 |
| **MR111** | 47.9 | 30.8 |
| **MR112** | 13.5 | 4.3 |
| **MR115** | 211.3 | 94.0 |
| **MR116** | 31.2 | 5.5 |
| **MR117** | 155.0 | 52.0 |
| **MR118** | 7.0 | 3.0 |
| **MR119** | 106.3 | 21.0 |
| **MR121** | 255.3 | 108.7 |

**Supplementary Table 4. NanoBRET target engagement binding affinities.** Extracted $IC_{50}$ data of electrophilic warhead binding to NanoLuc-tagged Brd4-BD2$^{L387A,E438C}$ as a result of Inhibition of the specific binding of 1 µM ET-JQ1-Bodipy in HEK293 cells. Data are mean ± SE from N4 independent repeats. IC50 values calculated as mean (±S.E.M.) from four independent biological experiments using log(inhibitor) vs. response (three parameters) using Graphpad Prism Version 10.2.3.

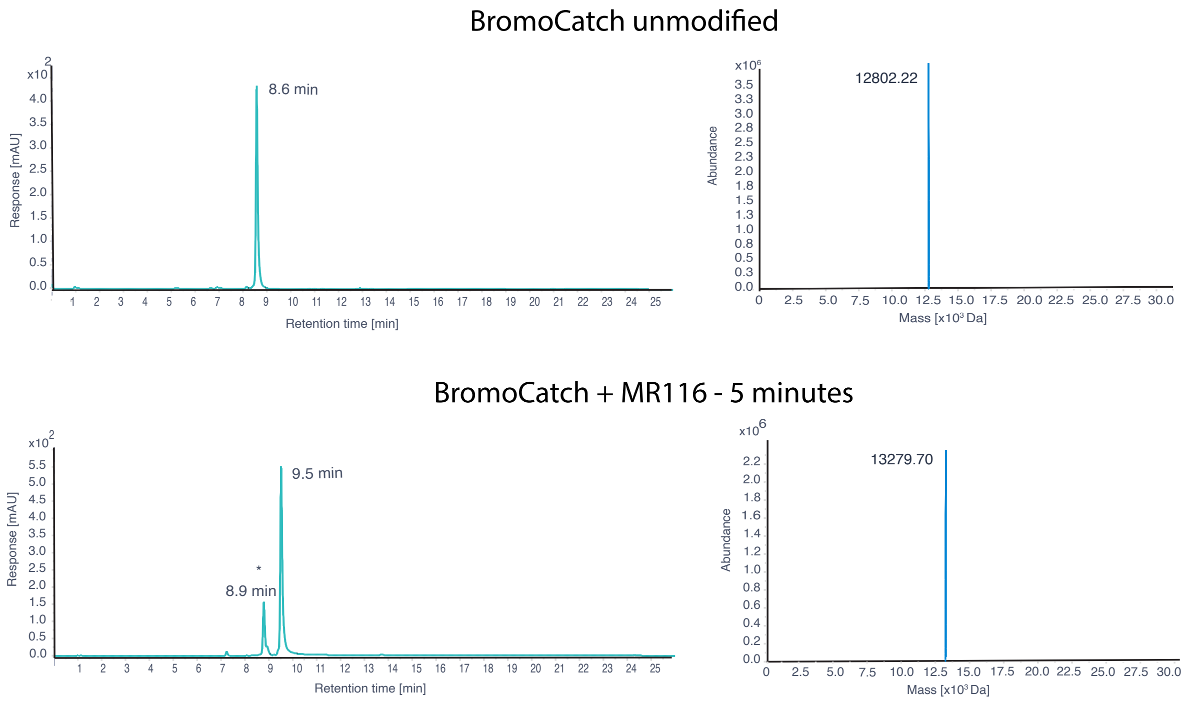


**Supplementary Figure 6. Evidence of rapid kinetic formation of covalent adduct.** BromoCatch is modified within 5 min of incubation with the acrylamide ligand MR116. INTACT-MS analysis after 5 min incubation at a 2:1 (ligand:protein) shows full modification within 5 min of incubation. * no protein mass envelope detected (excess ligand absorbtion).

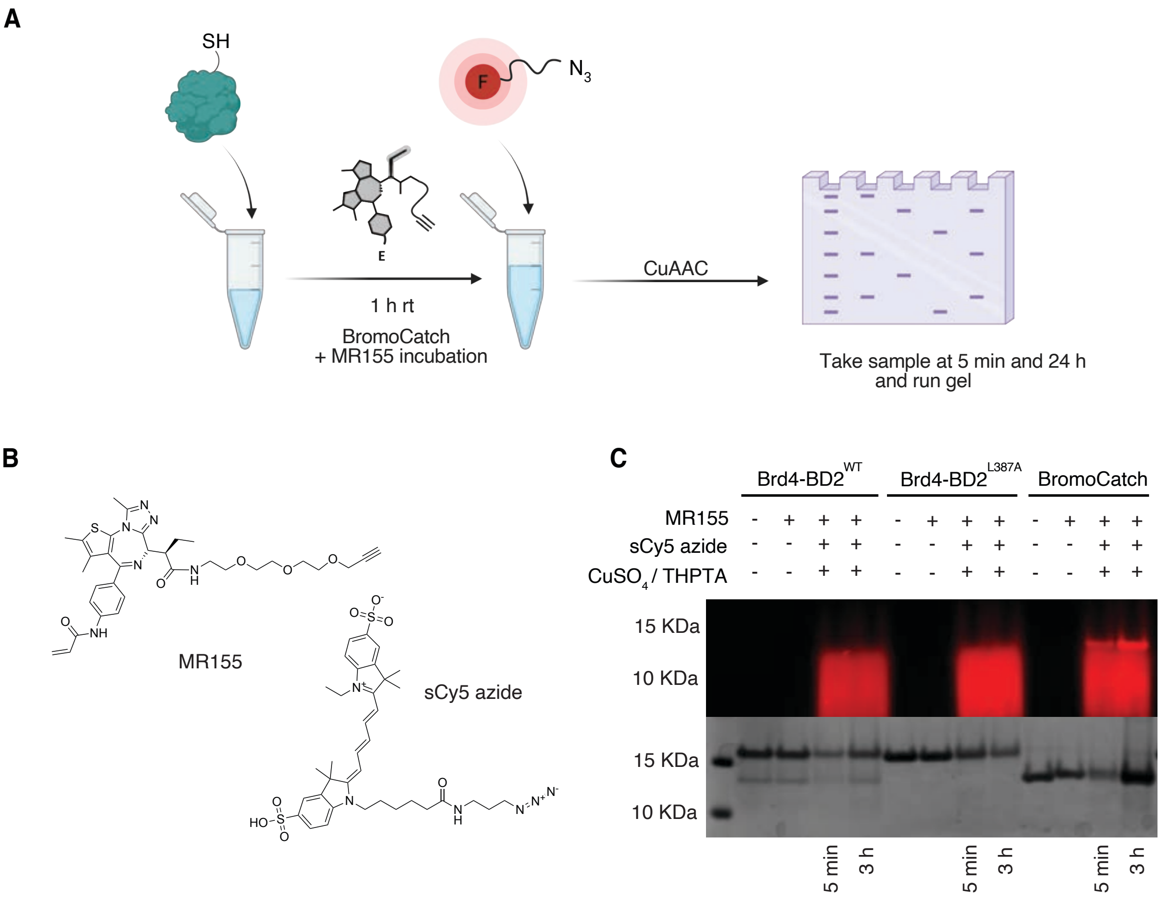


**Supplementary Figure 7. BromoCatch can be used for clicking fluorophores in a to-step CuAAC mediated reaction.** A) Workflow for the CuAAC experiment. B) The alkyne bearing probe MR155 (1.5 eq) was pre-incubated with BromoCatch, Brd4-BD2 WT or Brd4-BD2$^{L387A}$ (purified proteins) for 1 h before adding azide sulfonated Cy5 (excess) and the CuAAC reagents (CuSO4/THTPA/NaAsc) and samples were taken at 5 min and 3 h and analysed by SDS-PAGE. The conjugation selectively happens for BromoCatch, with no labelling observed for the Brd4-BD2 WT or Brd4-BD2$^{L387A}$ proteins.

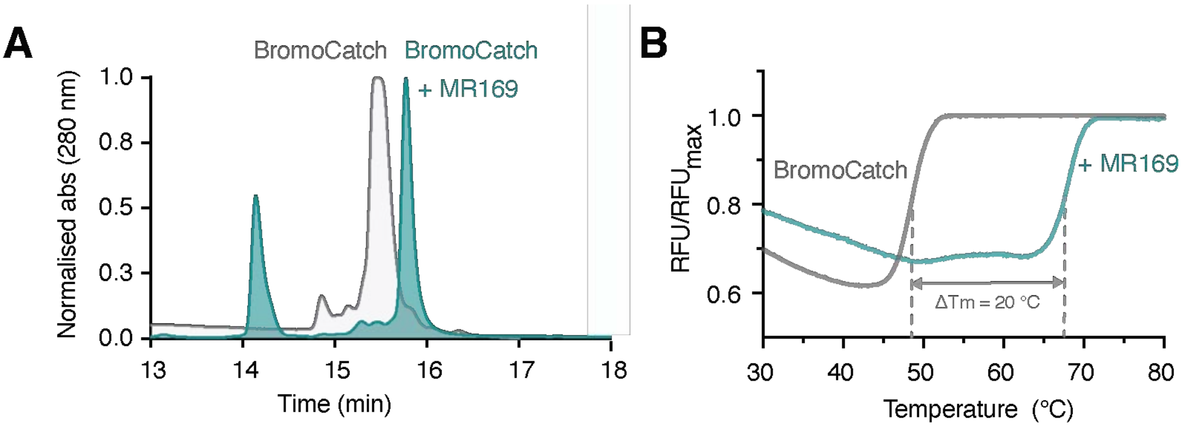


**Supplementary Figure 8.** MR169 biotin probe in vitro validation. HPLC-UV (280 nm) showed full covalent modification of BromoCatch when incubated in a 2:1 ratio (L:P) for 2 hours at room temperature. B) The significant protein stabilisation effect was measured by nanoDSF and confirmed to be maintained for the biotin bivalent ligand MR169.

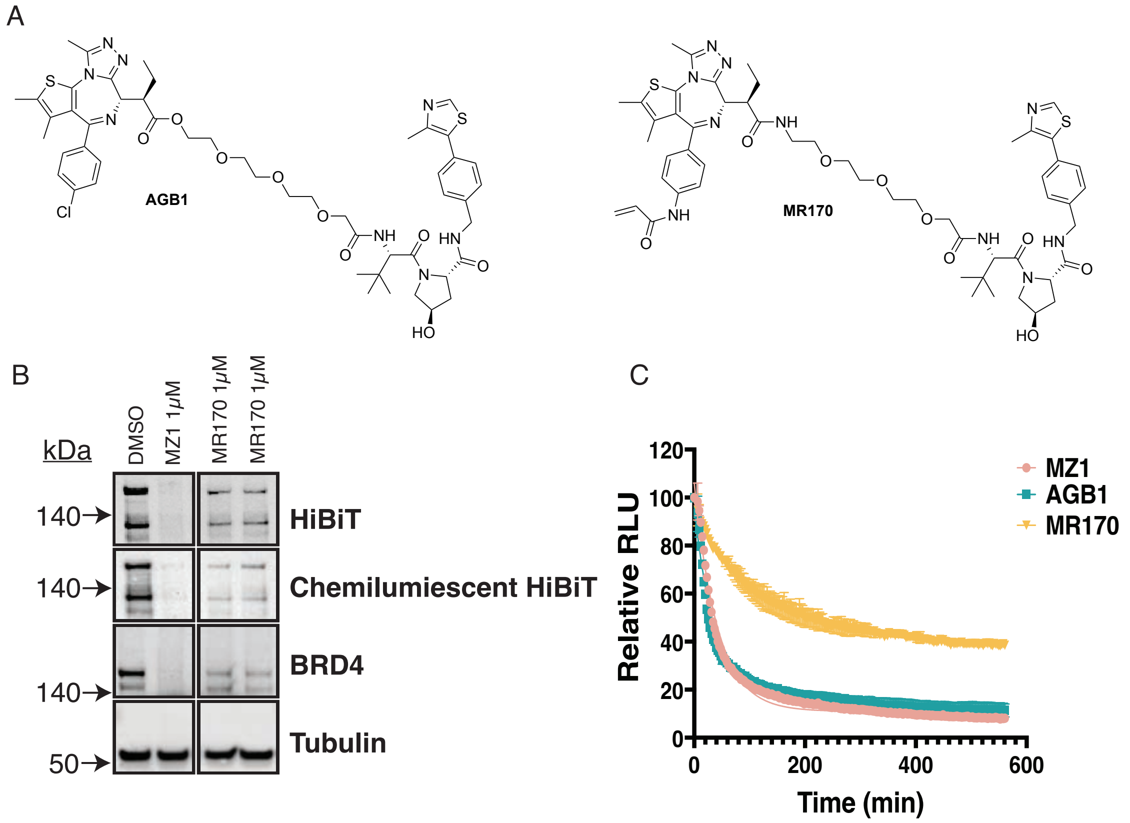


**Supplementary Figure 9. MR170 degrader activity.** A) Structures of AGB1 and MR170 used to test PROTAC and covalent degrader activity against BromoCatch in an endogenous Bromocatch-Brd4 HEK293 cell line. B) Immunoblot validation of the BromoCatch Knock-in in Brd4 using a 1µM treatment of MZ1 (Positive control) DMSO (vehicle) and MR170 and left to incubate for 6 hours prior to lysis. C) Live cell kinetic study of the activity of MZ1, AGB1 and MR170 in an endogenous HiBiT-Bromocatch-Brd4 Hek293 cell line. Cells were transiently transfected with the pCMV-LgBit plasmid prior to initiation of experiment and then treated with 1 µM MZ1 and AGB1 or 2.5 µM MR170 and continuously imaged for 9 hours.

| | **MZ1 (1µM)** | **AGB1 (1µM)** | **MR170 (2.5µM)** |
|---|---|---|---|
| **$T^{1/2}$ (Minutes)** | 29.2 | 26.7 | 83.8 |
| **$D_{MAX}$ [9H]** | 89.1 | 85.8 | 60.9 |

**Supplementary Table 5. $T^{1/2}$ and Dmax values for MZ1, AGB1 and MR170 derived from live cell kinetic assay.** $T^{1/2}$ and $D_{Max}$ values extracted from live cell kinetic degradtion assay in an endogenous HiBiT-Bromocatch-Brd4 HEK293 cell line. Data are mean from N3 independent repeats. $T^{1/2}$ and $D_{Max}$ values calculated as mean from three independent biological experiments using “one phase decay’ model from Graphpad Prism Version 10.2.3.

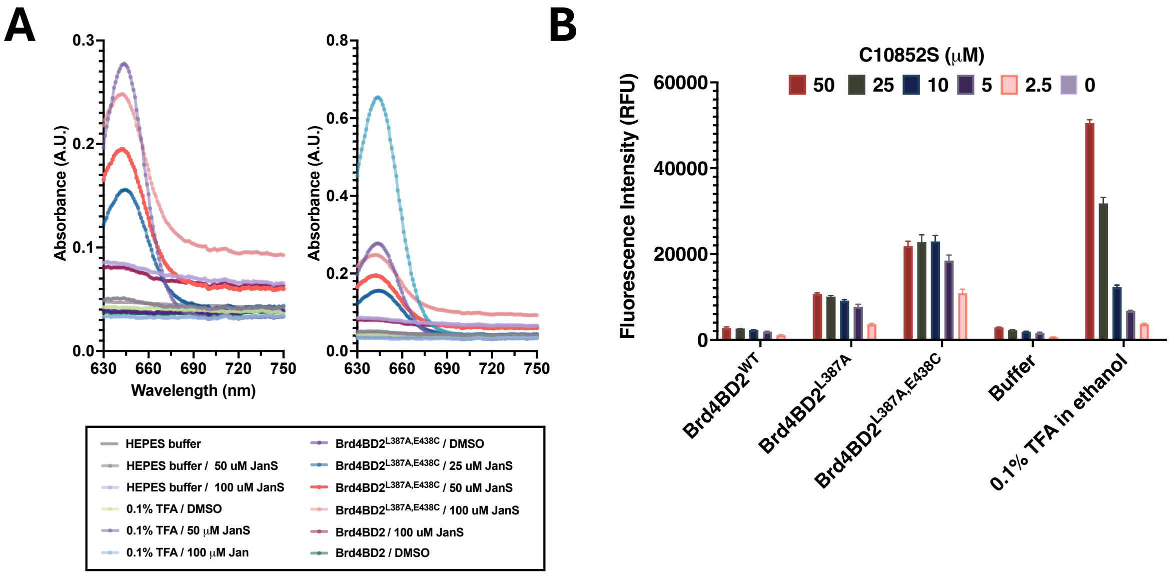


**Supplementary Figure 10. Fluorogenic switch on with the C18052S probe.** A) Extended data from Figure 7 where absorbance in presence of different concentrations of the probe and additional negative and positive controls are included. B) Extended data from figure 7 to show the fluorogenic response in presence of the different proteins and including the positive control of 0.1% TFA in ethanol. Importantly, the maximal fluorescence emission (RFU) at 10 µM of C10852S is slightly lower than the maximal achieved upon binding BromoCatch.

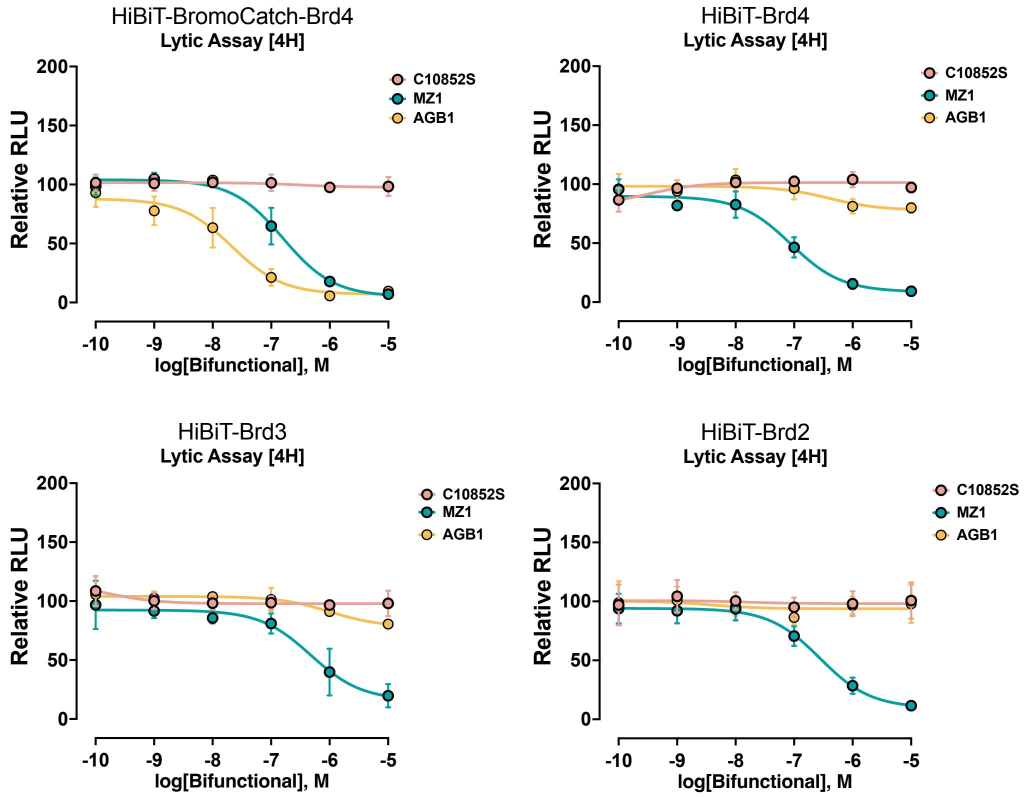


**Supplementary Figure 11. PROTAC-induced degradation of BromoCatch protein.** HiBiT lytic degradation of BromoCatch-Brd4 and wild-type Brd4, Brd3 and Brd2 using C10852S, MZ1 and AGB1 at 10µM-100pM. Cells were incubated with compounds for 4 hours prior to addition of HibIt lytic reagent and subsequent imaging on a BMG Labtech PHERAstar luminescence plate reader. Plots created in Graphpad Prism Version 10.2.3 using log(inhibitor) vs. response (three parameters). Data are mean ± SE from N3 independent repeats.

| $DC_{50}$ (nM) | MZ1 | AGB1 | C10852S |
|---|---|---|---|
| **BromoCatch** | 158.7 | 21.3 | N.A |
| **Brd4** | 88.0 | N.A | N.A |
| **Brd3** | 477.7 | N.A | N.A |
| **Brd2** | 275.2 | N.A | N.A |

**Supplementary Table 6. $DC_{50}$ values for MZ1, AGB1 and C10852S measured in lytic cell degradation assay.** $DC_{50}$ values extracted from HiBiT lytic degradtion assay in endogenous HiBiT-Bromocatch-Brd4, HiBiT-Brd4, HiBiT-Brd3 and HiBiT-Brd2 HEK293 cell lines. Data are mean from N3 independent repeats. $DC_{50}$ values calculated as mean from three independent biological experiments using log(inhibitor) vs. response (three parameters) using Graphpad Prism Version 10.2.3. N.A. - not applicable.

**H2B-(333-460)BRD4BD2$^{L387A + E438C}$**
**N1 U2OS**

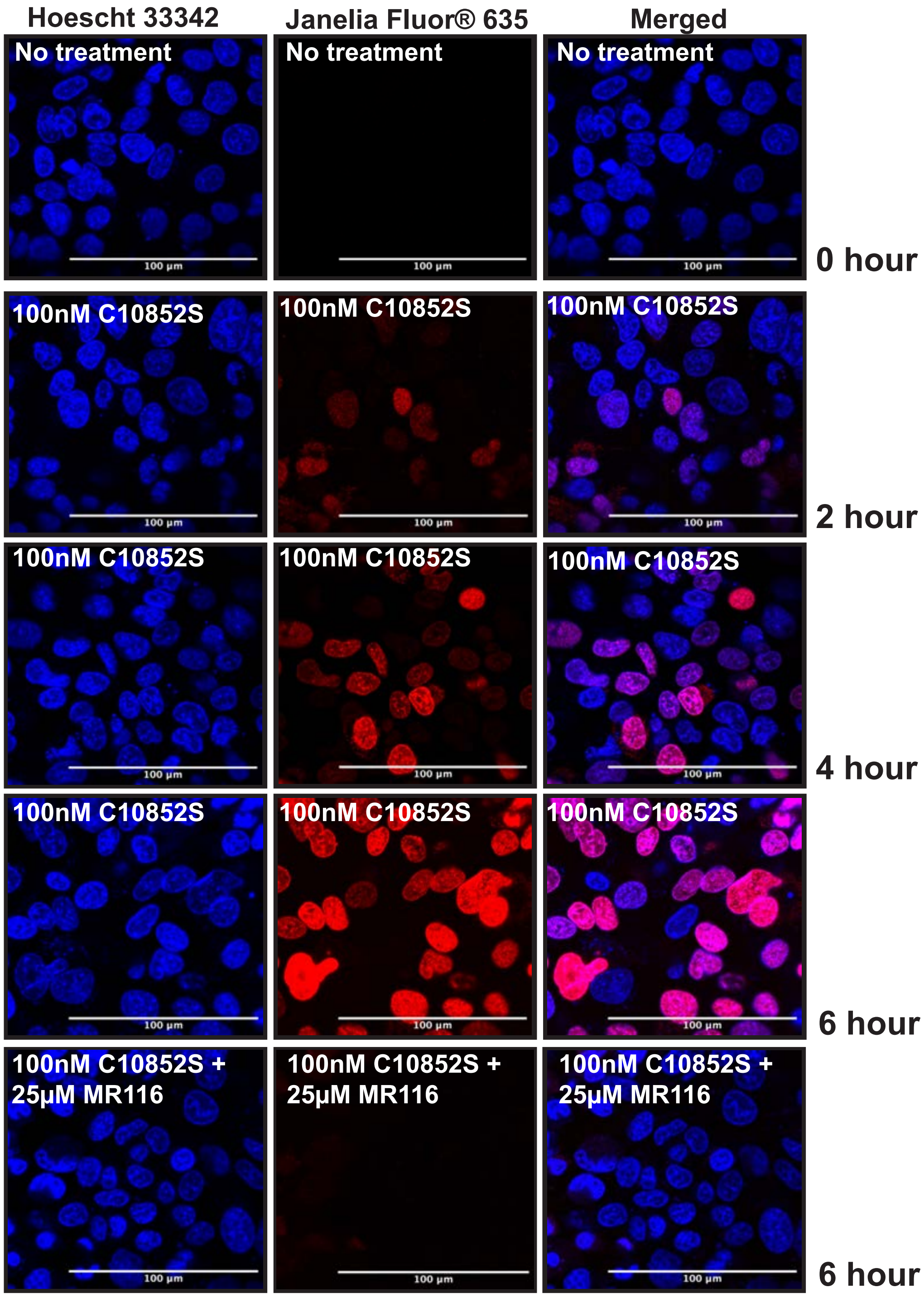

**H2B-(333-460)BRD4BD2$^{L387A + E438C}$**
**N2 U2OS**

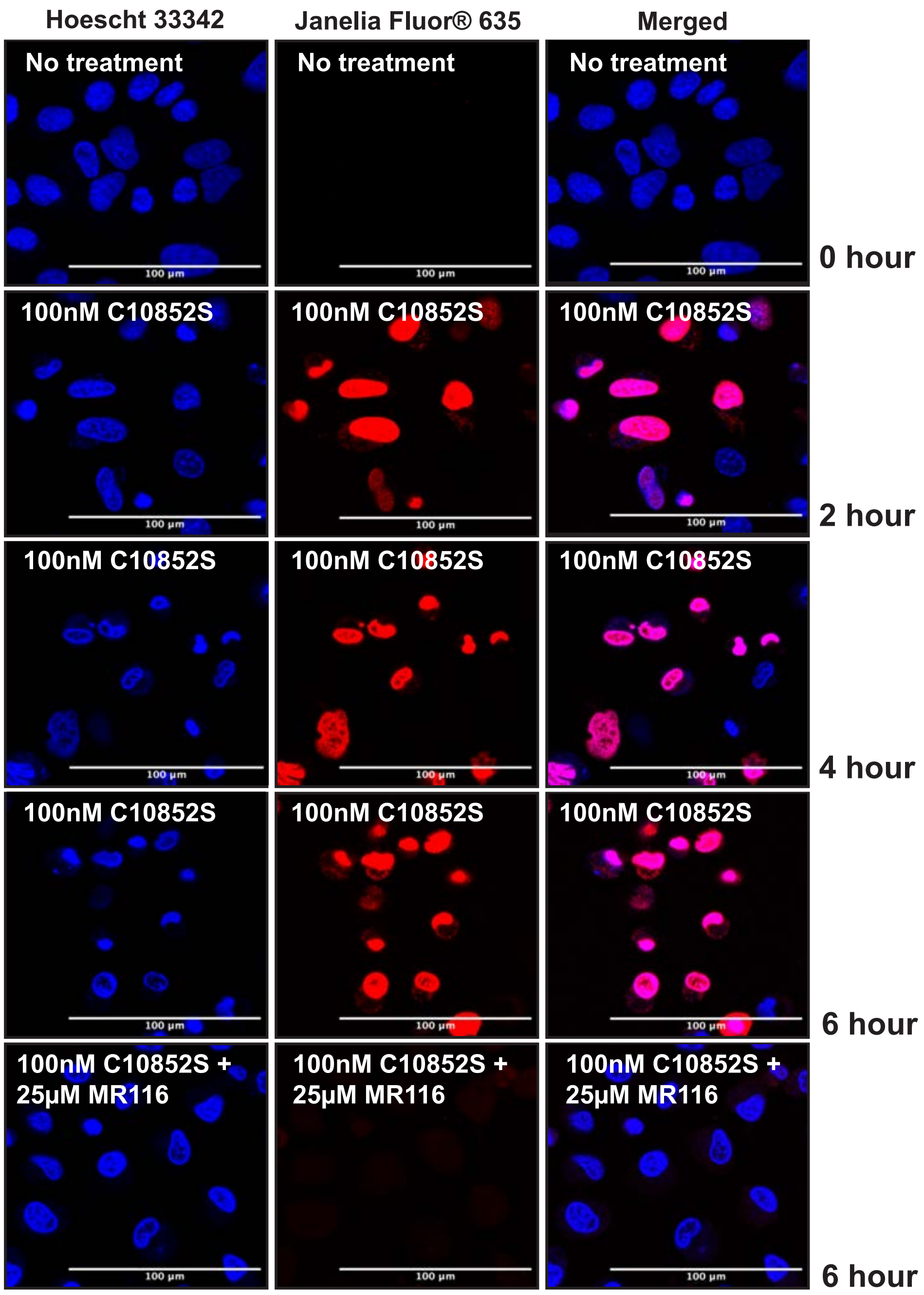


**Supplementary Figure 12. C10852S 200nM treatment of H2B-(333-460) BromoCatch in U2-O S.** Live-cell confocal imaging of H2B-BromoCatch(333-460) transiently transfected in U2-O S cells treated with C10852S. Cells were incubated in DMSO, 200nm C10852S and imaged up to 6 hours. Hoechst 33342 nuclear counterstain and JF 635 fluorescence detected. N2 independent repeats.

### H2B-(351-460)BRD4BD2$^{L387A + E438C}$
## N1 U2OS

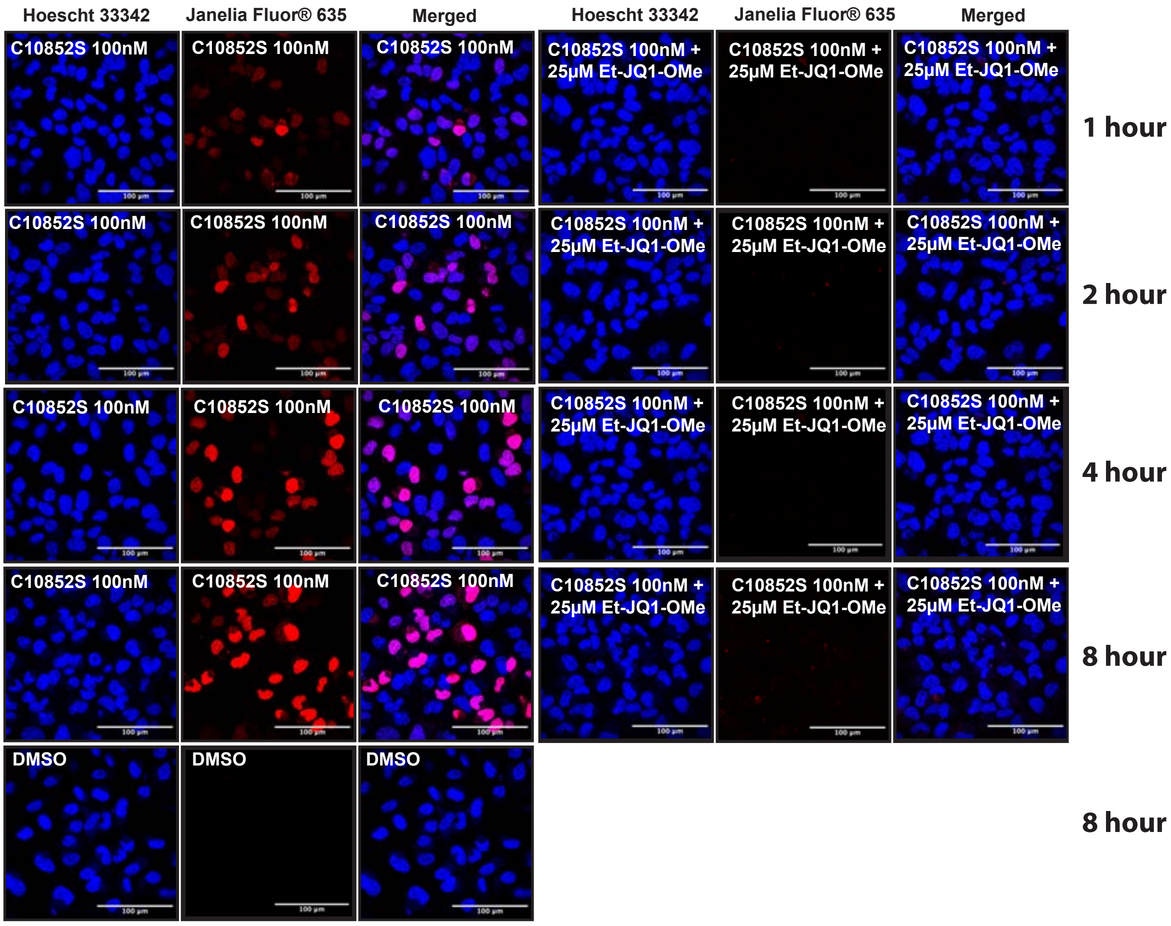


### Wild-type U2OS
## N1

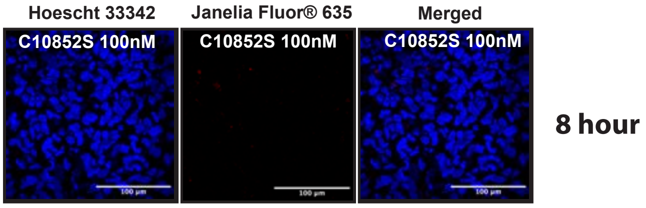

**H2B-(351-460)BRD4BD2$^{L387A + E438C}$**
**N2 U2OS**

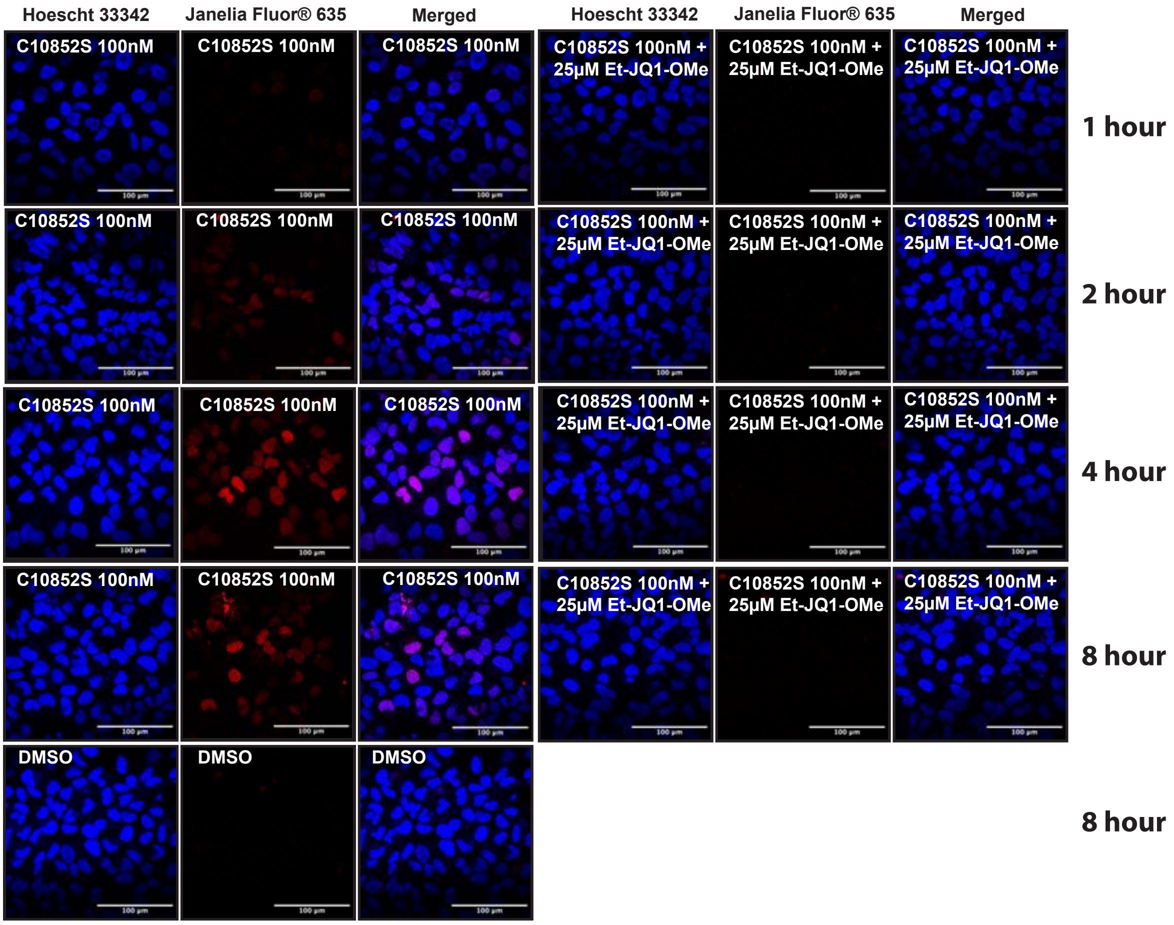


**Wild-type U2OS**
**N2**

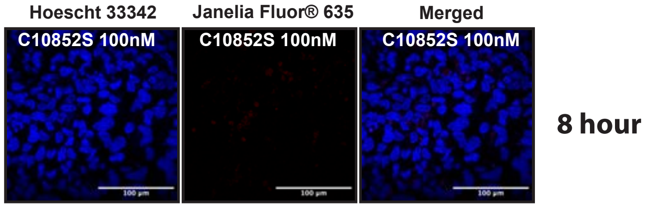


**Supplementary Figure 13. C10852S 100nM treatment of H2B-(351-460)BromoCatch in U2-O S.** Live-cell confocal imaging of H2B-BromoCatch(351-460) transiently transfected in U2-O S cells treated with C10852S. Cells were incubated in DMSO, 200nm C10852S or a 100nM C10852S cotreatment with 25µM ET-JQ1-OMe for 8 hours prior to imaging. Hoechst 33342 nuclear counterstain and JF 635 fluorescence detected. N2 independent repeats.

### H2B-(351-460)BRD4BD2$^{L387A + E438C}$
### N1 HEK293FT

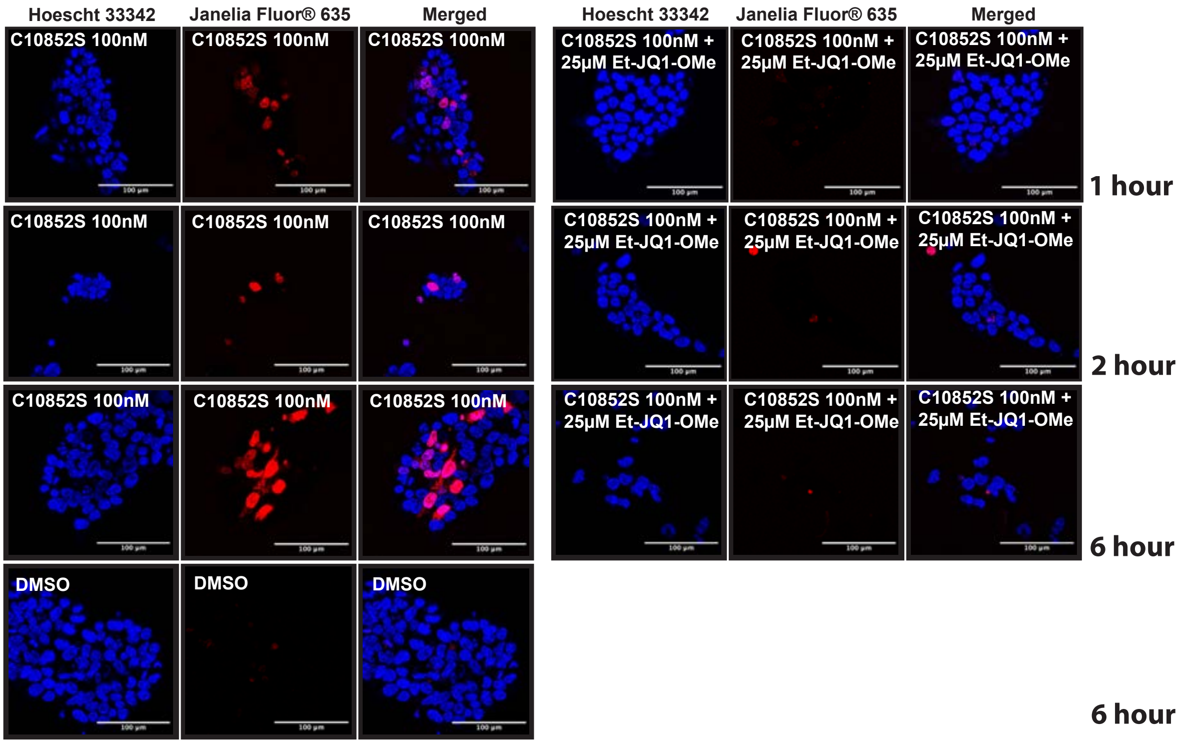


### Wild-type HEK293FT
## N1

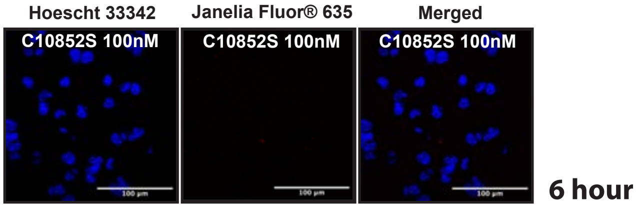

### H2B-(351-460)BRD4BD2$^{L387A + E438C}$
### N2 HEK293FT

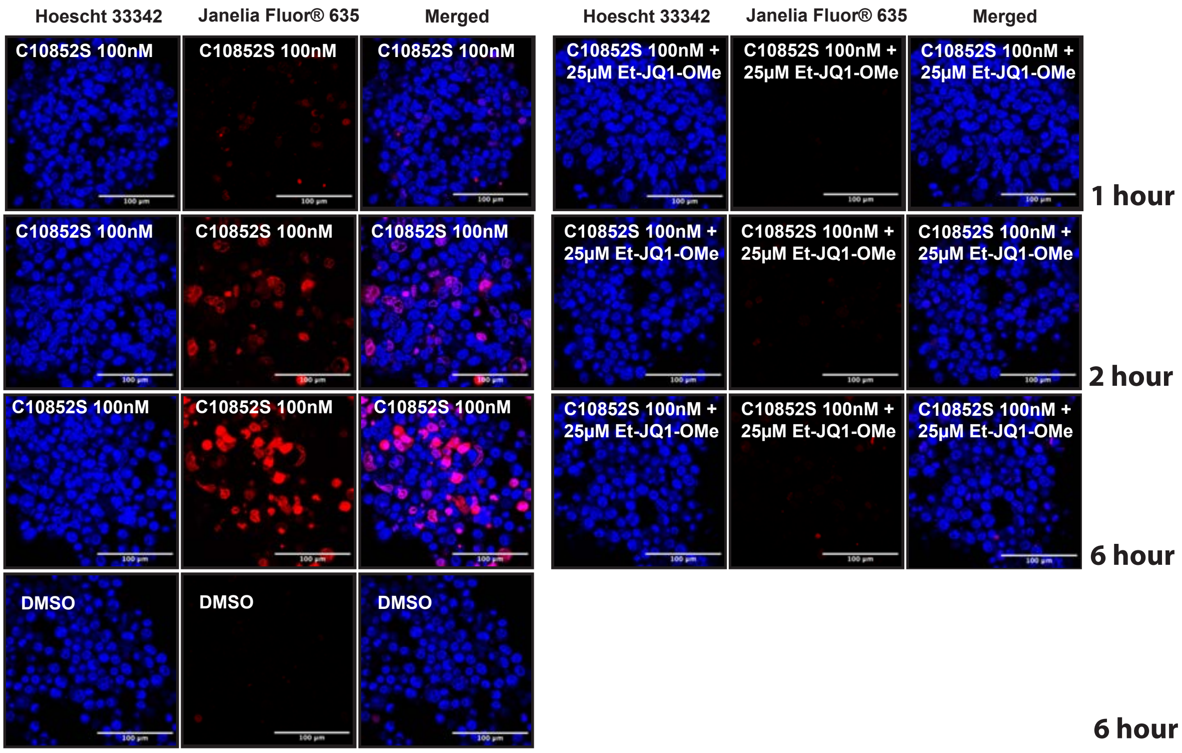


### Wild-type HEK293FT
## N2

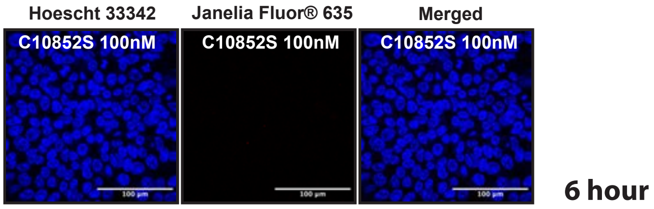

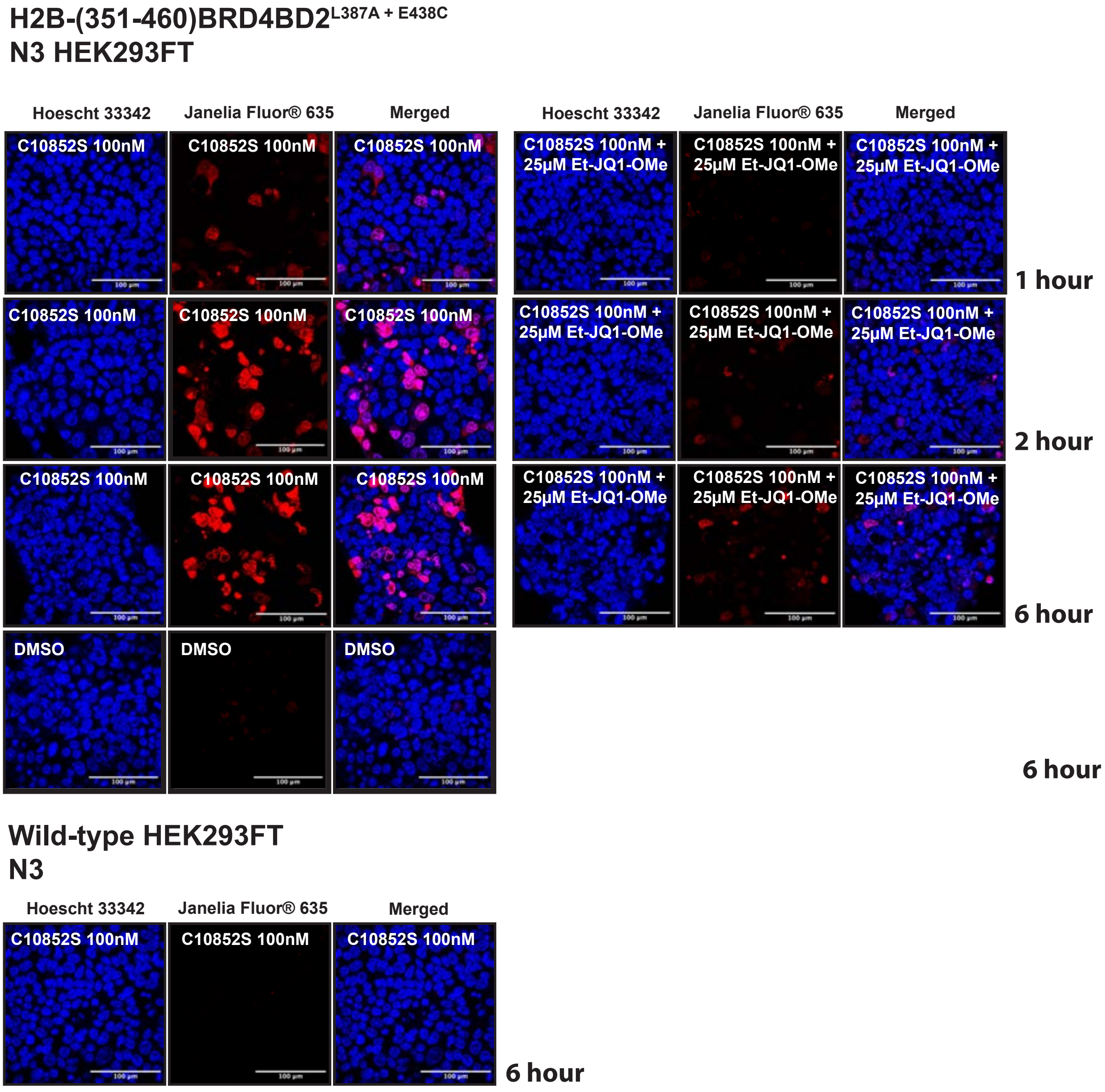


**Supplementary Figure 14. C10852S 100nM treatment of H2B-(351-460)BromoCatch in HEK293FT.** Live-cell confocal imaging of H2B-BromoCatch(351-460) transiently transfected in HEK293FT cells treated with C10852S. Cells were incubated in DMSO, 100nm C10852S or a 100nM C10852S cotreatment with 25µM ET-JQ1-OMe for 6 hours prior to imaging. Hoechst 33342 nuclear counterstain and JF 635 fluorescence detected. N3 independent repeats.

**Biology and biophysical assays:**

**Supplementary Table 7. List of buffers used for *in vitro* and *in cellulo* experiments.**

| Buffer | Composition |
|---|---|
| Activity buffer | 25 mM HEPES, 100 mM NaCl, 1 mM TECP |
| Loading buffer | 4×LDS sample buffer, 2% DTT |
| MES Running Buffer | 50 mM MES, 50 mM Tris Base, 0.1% SDS, 1 mM EDTA, pH 7.3 |
| MOPS Running Buffer | 50 mM MOPS, 50 mM Tris Base, 0.1% SDS, 1 mM EDTA, pH 7.7, 0.01-0.09% N,N-dimethylformamide |
| Transfer buffer | 10% v/v 10x Transfer Buffer, 20% v/v Ethanol, 80% v/v $ddH_2O$ |
| TBS-T buffer | TBS-T, 0.1% Tween |
| RIPA Buffer | 1% Triton X-100, 0.1% SDS, and 1:200 Protease inhibitor cocktail |

**Plasmids for recombinant protein expression in *E. coli***

To generate plasmids for recombinant expression in *E. coli* DNA encoding the respective Brd2-BD2 or Brd4-BD2 mutant, plasmids were PCR amplified using primers encoding 5' BamHI and 3' EcoRI restriction sites. The PCR fragments were double digested with BamHI and EcoRI, gel purified and extracted. Mutant fragments were ligated into pRSF-DUET1 vectors with N-terminal 6xHis-TEV or His-SUMO. All sequences were confirmed by conventional sequencing. Additional mutations were introduced using site directed mutagenesis with the relevant primer pairs for the desired point mutation (Tables 5 and 6).

**Supplementary Table 8. Protein expression plasmids.**

| Protein | Vector | Tag / Cleavage Site | Cloning | Antibiotic | Restriction Primers | Mutation primers |
|---|---|---|---|---|---|---|
| Brd4-BD2 L387A+E338C (351-460) | pRSF-DUET | His-TEV | BamHI-EcoRI | KAN | Olg13/Olg14 | Olg-80/Olg-81/Olg-84/Olg-85 |
| Brd4-BD2 L387A+M442C (351-460) | | | | | Olg13/Olg14 | Olg-80/Olg-81/Olg-86/Olg-87 |
| Brd4-BD2 (333-460) | | | | | Olg132/Olg14 | Olg132/Olg14 |
| Brd4-BD2 L387A (333-460) | | | | | Olg132/Olg14 | Olg80/Olg81 |
| Brd4-BD2 L387A+E338C (333-460) | | His-SUMO | | | Olg132/Olg14 | Olg132/Olg14 |
| Brd2-BD2 L383A + D434C (351-460) | DUET | His-MBP-TEV | | | Olg229/Olg230 | Olg232/Olg233/Olg234/Olg235 |

**Supplementary Table 9. Primers for DNA constructs.**

| Name | Construct | DNA sequence (5' to 3') | Direction |
|---|---|---|---|
| Olg-13 | Brd4-BD2 (351-C) | taagcaGGATCCTCGGAGCAGCTCAAGTGCTGCAGC | Forward |
| Olg-132 | Brd4-BD2 (333-C) | taagcaGGATCCAAGGACGTGCCCGACTCTCAGCAG | Forward |
| Olg-14 | Brd4-BD2 (459-N) | tgcttaGAATTCTTAGTCCGGCATCTTGGCAAAGCGCATTTCG | Reverse |
| Olg-80 | Brd4D2 (L387A) | GACGTGGAGGCACTGGGCGCACACGACTACTGTGACATCAT | MUT-forward |
| Olg-81 | Brd4-BD2 (L387A) | ATGATGTCACAGTAGTCGTGTGCGCCCAGTGCCTCCACGTC | MUT-reverse |

| | | | |
|---|---|---|---|
| Olg-84 | Brd4D2 (E438C) | GTACAACCCTCCTGACCATTGCGTGGTGGCCATGGCC CGCAAG | MUT-forward |
| Olg-85 | Brd4D2 (E438C) | CTTGCGGGCCATGGCCACCACGCAATGGTCAGGAGG GTTGTAC | MUT-reverse |
| Olg-86 | Brd4D2 (M442C) | GACCATGAGGTGGTGGCCTGCGCCCGCAAGCTCCAGG ATG | MUT-forward |
| Olg-87 | Brd4D2 (M442C) | CATCCTGGAGCTTGCGGGCGCAGGCCACCACCTCATG GTC | MUT-reverse |
| Olg-234 | Brd2-BD2 (D434C) | GTATAACCCGCCGGACCATTGCGTCGTAGCAATGGCC CGCAAG | MUT-forward |
| Olg-235 | Brd2-BD2 (D434C) | CTTGCGGGCCATTGCTACGACGCAATGGTCCGGCGGG TTATAC | MUT-reverse |
| Olg-229 | Brd2-BD2 (348-C) | taagcaGGATCCGCAAACCCTGAACAACTTAAACATTGC AATGGTATTTAAAAGAG | Forward |
| Olg-230 | Brd2-BD2 (455-N) | tgcttaGAATTCTTAATCGGGCATCTTTGCATAGCGGAAC TCG | Reverse |
| Olg-232 | Brd2-BD2 (L383A) | GTGGACGCCAGTGCCTTGGGAGCACACGACTATCATG ATATTATCAAG | MUT-forward |
| Old-233 | Brd2-BD2 (L383A) | CTTGATAATATCATGATAGTCGTGTGCTCCCAAGGCA CTGGCGTCCAC | MUT-reverse |

**Protein expression and purification**

Human Brd2-BD2 or Brd4-BD2 mutants were expressed with an N-terminal $His_6$-TEV tag in *E. coli* BL21(DE3) at 37 °C with LB supplemented with 50 µg/ml kanamycin once $OD_{600}$ reached 0.8. Protein expression was induced with 0.5 mM IPTG with 50 µM $MgCl_2$ added at induction and grown overnight at 18°C. After harvesting, the cells were resuspended in buffer containing 120 mM sodium phosphate buffer, 500 mM NaCl and 40 mM imidazole and $MgCl_2$ (1mM) and DNase I (10 µg/mL) added, and cells were lysed using Continuous Flow Cell Disruptor (Constant Systems) at 30,000 psi. Cell lysates were clarified by centrifugation at 18,000 g for 30 minutes at 4°C. The lysate was filtered and loaded to HisTrap FF affinity column (GE Healthcare) and eluted with 120 mM sodium phosphate buffer, 500 mM NaCl and 500 mM imidazole. The proteins were then dialyzed against 25 mM HEPES, pH 7.5, 150 mM NaCl and 1 mM TCEP with TEV protease (1:100 ratio) overnight. This mixture was passed through a HisTrap FF column, concentrated in a 3,500 MWCO centrifugal unit (Amicon) and loaded on a Superdex 16/600 size exclusion column pre-equilibrated in 25 mM HEPES, 150 mM NaCl, 1 mM TCEP, pH 7.5. Pure variants of Brd4 eluted ~0.74 cv and were confirmed using SDS-PAGE. The final proteins were all >95% pure, concentrated, and stored at -80°C until further use. All chromatography purification steps were performed using a BioRad NGC system at 4°C. Purity and= mass of proteins was confirmed by LC-MS.

**Supplementary Table 10. Protein sequences.**

| Protein | Residues | Sequence | MW (g/mol) |
|---|---|---|---|
| Brd4-BD2 | 333-460 | KDVPDSQQHPAPEKSSKVSEQLKCCSGILKE MFAKKHAAYAWPFYKPVDVEALGLHDYCDI IKHPMDMSTIKSKLEAREYRDAQEFGADVRL MFSNCYKYNPPDHEVVAMARKLQDVFEMRF AKMPDE | 14688.90 |
| Brd4-BD2$^{L387A}$ | 333-460 | VKDVPDSQQHPAPEKSSKVSEQLKCCSGILK EMFAKKHAAYAWPFYKPVDVEALGAHDYC DIIKHPMDMSTIKSKLEAREYRDAQEFGADV RLMFSNCYKYNPPDHEVVAMARKLQDVFE MRFAKMPDE | 14875.07 |
| Brd4-BD2$^{L387A,E438C}$ | 333-460 | SKDVPDSQQHPAPEKSSKVSEQLKCCSGILKE MFAKKHAAYAWPFYKPVDVEALGAHDYCDI IKHPMDMSTIKSKLEAREYRDAQEFGADVRL | 14707.92 |

| | | | |
|---|---|---|---|
| | | MFSNCYKYNPPDHCVVAMARKLQDVFEMRF AKMPDE | |
| Brd4-BD2$^{L387A,E438C}$ | 351-460 | SEQLKCCSGILKEMFAKKHAAYAWPFYKPV DVEALGAHDYCDIIKHPMDMSTIKSKLEARE YRDAQEFGADVRLMFSNCYKYNPPDHCVVA MARKLQDVFEMRFAKMPDE | 12661.70 |
| Brd4-BD2$^{L387A,M442C}$ | 351-459 | SEQLKCCSGILKEMFAKKHAAYAWPFYKPV DVEALGAHDYCDIIKHPMDMSTIKSKLEARE YRDAQEFGADVRLMFSNCYKYNPPDHEVVA CARKLQDVFEMRFAKMPD | 12659.62 |
| Brd2-BD2$^{L383A,D434C}$ | 348-455 | EQLKHCNGILKELLSKKHAAYAWPFYKPVD ASALGAHDYHDIIKHPMDLSTVKRKMENRD YRDAQEFAADVRLMFSNCYKYNPPDHCVVA MARKLQDVFEFRYAKMPD | 12963.82 |

**Differential Scanning Fluorimetry**

Differential Scanning Fluorimetry (DSF) experiments were performed on a Biorad CFX96 RT-PCR machine or in a Nanotemper Phanta (NanoTemper Technologies GmbH). For assays in the Biorad CFX96 RT-PCR machine, ligand (50 µM) or DMSO stock were incubated with Brd4-BD2$^{WT}$, Brd4-BD2$^{L387A}$ Brd4-BD2$^{L387A,E438C}$, Brd4-BD2$^{L387C}$, Brd4-BD2$^{L387A,M442C}$ (10 µM, 50 µl) in 25 mM HEPES, 100 mM NaCl, 1 mM TECP for 2 hours at room temperature followed by addition of SYPRO Orange (1 uL) to give a final dilution of 5 × SYPRO Orange, the final DMSO concentration was 4 %. The temperature was ramped up in 1 °C steps between 25 and 95 °C with 30 s incubation at each step. Melting curves were analysed by determining the minimum of the first derivative using the Biorad CFX Manager. For analysis in the Nanotemper Phanta (NanoTemper Technologies GmbH), a similar procedure was used, but no addition of SYPRO Orange was required, samples containing a mixture of the protein (10 µM) and ligand (50 µM) were taken into capillaries for subsequent reading by Nanotemper, the temperature was similarly ramped up in 1 °C steps between 25 and 95 °C with 30 s incubation at each step. Values of $T_m$ and were obtained from the melting curve at 350 nm emission (inherent fluorescence emission of aromatic amino acids).

**Covalent modification of protein LC-MS analysis**

Brd4-BD2, Brd4-BD2$^{L387A}$, Brd4-BD2$^{L387A,E438C}$ and Brd4-BD2$^{L387A,M442C}$ (50 µM), were incubated with a 1:2 P:L concentration of ligands (100 µM) in activity buffer at room temperature or at 37 °C. At 2 hours, the sample was precipitated by the addition of 4 volumes of cold methanol. The precipitated protein was pelleted by centrifugation and resuspended in an aqueous solution of 15% acetonitrile and 0.1% TFA. Samples were separated by HPLC over 20 minutes on a C3 column using a 10–95% gradient of acetonitrile and analysed using an Agilent 6130 quadrupole MS. Spectra were deconvoluted and integrated using Agilent LC/MSD OpenLab. The quantification of modified protein was done by integration of peaks on the 280 nm UV channel trace for the unmodified and modified protein.

**Sodium dodecyl sulfate polyacrylamide gel electrophoresis of recombinant (modified) proteins**

Protein samples were separated by gel electrophoresis on NuPAGE™ 4–12%, Bis-Tris precast polyacrylamide gels (Thermofisher) unless otherwise stated. Protein or cell lysate proteins were separated by SDS-PAGE gel electrophoresis. Samples (10 µL) were prepared addition of loading buffer (10 uL, 4×LDS sample buffer, 2% DTT) and then heated at 95 °C for 5 min. Protein samples (15–28 µL) were loaded into NuPAGE 4-12% Bis-Tris Gels and electrophoresed for 45 min at 180V in MES buffer. The Protein Standards ladder (Bio-rad) (5 µL) was used as a molecular weight reference. Following electrophoresis, gels were imaged with ChemiDoc MP Imaging System for fluorescence read-out at the appropriate wavelength (for fluorescent labelling experiments) or stained for 15 min with Instant Blue Coomasie® stain (Abcam), destained, and imaged using Bio-rad molecular imager.

Bio-rad Image Lab 6.1 was used to process the data. Protein concentrations were measured prior to use in assays using a NanoDrop Microvolume Spectrophotometer.

***In vitro* protein labelling with biotin covalent ligand and streptavidin capture assay**

Brd4-BD2, Brd4-BD2$^{L387A}$ or Brd4-BD2$^{L387A,E438C}$ (50 µM), were incubated with acrylamide-based biotin containing covalent ligand **MR169** (75 µM) in activity buffer for 1 hour. At 1 hour, to this mixture (5 µL), a solution of streptavidin in activity buffer (45 µL, 10 eq., 270 µM) was added and incubated for 2 hours. After 2 hours of incubation with streptavidin, samples were taken and prepared for gel electrophoresis by mixing the sample (10 µL) with loading buffer (10 µL). Prepared samples (15 µL) were separated by NuPAGE™ 4–12%, Bis-Tris precast polyacrylamide gels as described above.

***In vitro* protein labelling with fluorescent covalent ligands**

Brd4-BD2, Brd4-BD2$^{L387A}$, Brd4-BD2$^{L387A,E438C}$, Brd4-BD2$^{L387A,M442C}$ (10 µM) were co-incubated with either DMSO or increasing concentrations of **MR202** or **C10852S** in activity buffer. Concentrations covered a range of concentrations from sub-stoichiometric up to excess (2.5 to 50 µM). Samples (10 µL) were mixed with a solution of LDS/ DTT (10 µL), loaded (15 µL) and separated by NuPAGE™ 4–12%, Bis-Tris precast polyacrylamide gels as described above and imaged for fluorescence before staining with Coomassie. The in-gel fluorescence ChemiDock channels: (TAMRA channel $\lambda_{ex}$: 500-XXX nm $\lambda_{em}$: 550–600 nm; JF635 channel $\lambda_{ex}$: 625–650 nm $\lambda_{em}$: 675–725 nm).

**Docking studies**

A model of the mutants Brd2-BD2$^{L383V,D434C}$ and Brd2-BD2$^{L383V,M438C}$ were generated by introducing the mutation with the maestro editing tools, using the x-ray structure of Br2BD2$^{L383V}$ co-crystallised with ET-JQ1-OMe (6YTM) as a template. The Brd2-BD2$^{L383V}$ co-crystal and the corresponding mutations were prepared using single point mutations and the Protein Preparation Wizard from Schrodinger, and the corresponding grids were generated with Glide. Ligands were prepared (Ligprep) and docked reversibly (Glide) or covalently (CovDock) in either mutant or reference crystal. The best 5 scored poses from each docked ligand were filtered and analysed visually with Maestro. Root mean square deviation (RMSD) was obtained with respect to the reference crystal for the common structures.

**Click test**

Click probe Copper catalysed Alkyne Azide Click (CuAAC) assays were performed in 1.5 mL Eppendorf tubes at 37 °C or at room temperature, in the dark, 15 and gently shaken, unless otherwise stated. The mutant protein Brd4-BD2$^{L387A,E438}$ (50 µM, 200 µL) was incubated with the alkyne containing ligand MR155 (75 µM) for 1 hour in 25 mM HEPES 7.5 pH, 100 mM NaCl 1 mM TECP. To this mixture, sCy5-azide (300 µM) was added followed by a premixed solution of THPTA/CuSO4 (5:1, 0.5 mM/0.1 mM) in 25 mM HEPES 7.5 pH, 100 mM NaCl 1 mM TCEP and NaAsc 20 (5 mM) as added. The reaction was mixed at 37 °C, after which point a ssamples were taken at 5 min or 3 hours. Samples were diluted by 5-fold, and prepared for gel electrophoresis by mixing the sample (10 µL) with a mixture of LDS/ DTT (10 µL). Prepared samples (10 µL) were separated by NuPAGETM 4–12%, Bis-Tris precast polyacrylamide gels. The 25 gels were the imaged with ChemiDock (sulfoCyanine 5 fluorescence in the Cyanine 5 channel (excitation range of 625–650 nm; emission range of 675–725 nm). The gels were subsequently stained with Instant Blue Coomasie® stain, destained, and images in the GelDock.

**Cell culture**

HEK293, HEK293FT and U2-O S were all purchased from the American Type Culture Collection (ATCC). All cell lines were cultured in 10 cm cell culture plates with Dulbecco's Modified Eagle Medium high glucose, GlutaMAX™ Supplement (HEK293 & HEK293FT)/ Dulbecco's Modified

Eagle Medium Nutrient Mixture F-12 (U-2 OS) (DMEM high glucose, GlutaMAX™ Gibco™ Catalog number: 10566016 /F12, Gibco™ Catalog number:11320033) in a humidified incubator with 5% CO2 at 37 °C. DMEM/F12 media was supplemented with 10% fetal bovine serum (FBS, Gibco™ Catalog number: A5256701). HEK293 FT containing the SV40 large T antigen (Invitrogen) was cultured in DMEM supplemented with 50 µg/ml gentamycin.

**Construct design**

DNA constructs for this project were all designed in Snapgene (Version 8.0.2). Construct used for NanoBRET validation of electrophilic warheads consisted of pcDNA3.1(+) vector harbouring a Brd4-BD2$^{L387A,\ E438C}$-Nanoluciferase 3' to the CMV promoter Construct used to probe functionality of the bifunctional probes using the elongated version of BromoCatch consisted of a pcDNA3.1(+) vector harbouring a H2B-Brd4-BD2$^{L387A,\ E438C}$ (containing residues 333-460 of Brd4-BD2) 3' to the CMV promoter.  Construct used to probe functionality of the bifunctional probes using the shortened version of BromoCatch) consisted of a pcDNA3.1(+) vector harbouring a H2B-Brd4-BD2$^{L387A,\ E438C}$ (containing residues 351-460 of Brd4-BD2) 3' to the CMV promoter. Donor construct used to produce the endogenously tagged BromoCatch Brd4 cell line consisted of a pMK-RQ vector with an donor insert consisting of an eGFP-IRES-HiBiT-Bromocatch-Brd4 insert in-between a 500 bp left and right homology arms for the N-terminus of Brd4.  Two single pBABED vectors harbouring U6-driven Brd4 gRNA sequences and a pCMV driven cas9 cassette were also developed with the help of the MRC-PPU CRISPR services (**Supplementary Table 11, 12**).

**Supplementary Table 11. Plasmids for live cell work**

| **Plasmid** | **Residues** | **Sequence** | **Vector Backbone** |
|---|---|---|---|
| NanoLuciferase-linker- Brd4-BD2$^{L387A,E438C}$ | 333-460 | MVFTLEDFVGDWRQTAGYNLDQVLEQG GVSSLFQNLGVSVTPIQRIVLSGENGLKID IHVIIPYEGLSGDQMGQIEKIFKVVYPVDD HHFKVILHYGTLVIDGVTPNMIDYFGRPY EGIAVFDGKKITVTGTLWNGNKIIDERLIN PDGSLLFRVTINGVTGWRLCERILAGSSG AIAKDVPDSQQHPAPEKSSKVSEQLKCCS GILKEMFAKKHAAYAWPFYKPVDVEAL GAHDYCDIIKHPMDMSTIKSKLEAREYR DAQEFGADVRLMFSNCYKYNPPDHCVV AMARKLQDVFEMRFAKMPDE* | pcDNA™3.1 (+) Mammalian Expression Vector |
| **H2B-linker-** Brd4-BD2$^{L387A,E438C}$ | 333-460 | MPEPSKSAPAPKKGSKKAITKAQKKDGK KRKRSRKESYSIYVYKVLKQVHPDTGISS KAMGIMNSFVNDIFERIAGEASRLAHYN KRSTITSREIQTAVRLLLPGELAKHAVSEG TKAVTKYTSSGAGAGAGAGAKDVPDSQ QHPAPEKSSKVSEQLKCCSGILKEMFAKK HAAYAWPFYKPVDVEALGAHDYCDIIKH PMDMSTIKSKLEAREYRDAQEFGADVRL MFSNCYKYNPPDHCVVAMARKLQDVFE MRFAKMPDE | pcDNA™3.1 (+) Mammalian Expression Vector |
| **H2B-linker-** Brd4-BD2$^{L387A,E438C}$ | 351-460 | MPEPSKSAPAPKKGSKKAITKAQKKDGK KRKRSRKESYSIYVYKVLKQVHPDTGISS KAMGIMNSFVNDIFERIAGEASRLAHYNK RSTITSREIQTAVRLLLPGELAKHAVSEGT KAVTKYTSSGAGAGAGAGASEQLKCCSG ILKEMFAKKHAAYAWPFYKPVDVEALGA HDYCDIIKHPMDMSTIKSKLEAREYRDAQ | pcDNA™3.1 (+) Mammalian Expression Vector |

| | | | |
|---|---|---|---|
| | | EFGADVRLMFSNCYKYNPPDHCVVAMAR KLQDVFEMRFAKMPDE* | |
| **eGFP-IRES-HiBiT**- Brd4-BD2$^{L387A,E438C}$-BRD4 | 333-460 | MSKGEELFTGVVPILVELDGDVNGHKFSV SGEGEGDATYGKLTLKFICTTGKLPVPWP TLVTTLTYGVQCFSRYPDHMKQHDFFKS AMPEGYVQERTIFFKDDGNYKTRAEVKFE GDTLVNRIELKGIDFKEDGNILGHKLEYN YNSHNVYIMADKQKNGIKVNFKIRHNIED GSVQLADHYQQNTPIGDGPVLLPDNHYLS TQSALSKDPNEKRDHMVLLEFVTAAGITL GMDELYK*RSAPLPPPPLTLLAEAAWNKA GVRLSICYFPPYCRLLAM*GPGNLALSS*R AFLGVFPLSPKECKVC*MS*RKQFLWKLL EDKQRL*RPFAGSGTPHLATGASAAKSHV YKIHLQRRHNPSATL*VG*LWKESNGSPQ AYSTRG*RMPRRYPIVWDLIWGLGAHAL HVFSRG*KNV*APRTTGTWFSFEKHDDN MATTMVSGWRLFKKISGGGGGGKDVPDS QQHPAPEKSSKVSEQLKCCSGILKEMFAK KHAAYAWPFYKPVDVEALGAHDYCDIIK HPMDMSTIKSKLEAREYRDAQEFGADVR LMFSNCYKYNPPDHCVVAMARKLQDVFE MRFAKMPDEGAGAGAGAGA | pMK-RQ |

**CRISPR knock-in of BromoCatch-Brd4 HEK293 cells.**

To perform the knock-in of BromoCatch into the N-terminus of BRD4, 1 mL of HEK293 cells in DMEM:10% FBS were plated at a density of 1x10$^6$ cells per mL into a single well of a six-well plate in the 24 hours leading up to the initiation of the experiment. The cells were subsequently transfected the following day using Fugene HD using the donor pMK-RQ vector containing 500bp homology arms of N-terminal Brd4 on either side of eGFP-IRES-HiBiT-BromoCatch(351-460) sequence (table 11). Simultaneously, the cells were transfected with two single gRNA/cas9 containing pBABED vectors (Table 12). The HEK293 cells were transfected with 1 µg of the pMK-RQ donor vector and woth 0.5 µg of each pBABED vector. The DNA was mixed together in 100µL of Optimem media and Fugene HD was added at a ratio of 1:3 DNA:Fugene HD or 2 µg:6 µL total and this mixture was left to form complexes for 20 minutes at room temperature. This mixture was then added dropwise to the adherent HEK293 cells in 2mL of fresh DMEM:10% FBS and the cells were left to incubate in a humidified incubator with 5% CO2 at 37 °C overnight. The following day cells were washed in PBS before fresh DMEM: 10% FBS was applied. The HEK293 cells were left to incubate in a humidified incubator with 5% CO2 at 37 °C for 72 hours to allow for recovery post-transfection. The surviving HEK293 cells were subsequently re-transfected with the CRISPR reagents for BromoCatch insertion into Brd4 using the same conditions stated above, allowing for a further 72 hours to recover post-transfection in a humidified incubator with 5% CO2 at 37 °C. Following this, the surviving HEK293 cells were subsequently prepared for fluorescence-activated cell sorting (FACS).

**Supplementary Table 12. gRNA sequences used for BromoCatch knock-in of Brd4.**

| **Target** | **Sequence** |
|---|---|
| ***BRD4*** | GTGGGATCACTAGCATGTCTG |
| ***BRD4*** | GACTAGCATGTCTGCGGAGAG |

**Cell sorting**

The surviving HEK293 cells from the previous stage were subsequently trypsinised using trypsin–EDTA (0.25%). Once in suspension, the trypsin–cell mixture was neutralized with DMEM:10%FBS. The cell suspension was pelleted at 1000 rpm for 5 min. The cell pellet produced was subsequently resuspended in optimem supplemented with 1% FBS at a concentration of $1\times10^6$ cells per mL. Wild-type HEK293 cells were used as a baseline control for GFP expression. Single-cell clones were generated by FACS using an SH800 cell sorter from Sony Biotechnology of the Dundee University Flow Cytometry and Cell Sorting Facility. A 488-nm laser was used for the excitation of fluorescence and generation of light scattering. Forward angle light scatter (FSC) and backscatter were detected using 488 ± 17 nm band-pass filters. Cells were distinguished from debris based on FSC-area (A) and SSC-A measurements. Single cells were distinguished from doublets and clumps based on FSC-A and FSC-width measurements. GFP fluorescence was detected using a 525 ± 50 nm band-pass filter, and autofluorescence was detected using a 600 ± 60 nm band-pass filter. GFP-positive cells were identified by first assessing the background GFP and autofluorescence of a control sample of cells which did not express GFP. Using the measurements for GFP and autofluorescence of this sample, a collection gate was set, which identified GFP-positive cells. The samples to be sorted were then analysed, and GFP-positive cells were sorted for collection. A single GFP +ve cell was sorted into each well of 6 × 96 well plate in 200 µL of 50% filtered pre-conditioned media from growing healthy cells and 50% fresh DMEM:10% FBS. Sorted plates were subsequently spun down at 1000 rpm for 5 minutes and the left to grow in a humidified incubator with 5% CO2 at 37 °C for 2 weeks. After 2 weeks, all visible colonies were expanded to 24, 12, and then subsequently 6-well plates and cryogenically persevered prior to validation.

**CRISPR validation**

Validation of BromoCatch knock-in at the N-terminus of BRD4 knock-in was accomplished initially by performing a HiBIT lytic assay on the FACS single-cell sorted CRISPR clones. The clones were initially trypsinised and the cell suspension was plated out onto individual wells of a white-walled 96 well plate at a density of $2\times10^4$ cells per well in 100 µL of DMEM:10% FBS. The HiBiT lytic reagent consisting of a lytic buffer, fumarizine substrate and the complementary LgBIT protein was then prepared and added to the plate containing the CRISPR cell suspension following manufacturer's instructions. Upon addition of the lytic substrate, the plate was spun on an orbital shaker for 5 min to encourage lysis and left for a further 5 min to reach peak luminescence. Luminescence was then recorded on a BMG Labtech PHERAstar luminescence plate reader. Data extracted from this analysis was analysed with GraphPad Prism (v. 10.4.1, GraphPad) and clones with detectable levels of luminescence was identified. A FACS sorted clone that displayed a high luminescent signal was chosen and subsequently validated by western blotting. This clone was plated at a density of $1\times10^6$ cells per well into two wells of a 6-well plate using 2mL of DMEM:10%FBS and left to incubate in a humidified incubator with 5% CO2 at 37 °C for16 hours prior to initiation of experimentation. This adherent FACS sorted HEK293 clone cell line was subsequently treated with DMSO (vehicle), MZ1 at 1µM or MR170 for a total treatment time of 6 hours. The clone was subsequently washed with PBS and 100 µL of RIPA lysis buffer containing cOmplete proteinase inhibitor cocktail and benzonase at manufacture specified concentrations was added to each condition. Total protein quantity was determined using the BCA protein assay (#23225, Pierce, Rockford, Illinois). The samples were then prepared and loaded thrice onto a NuPAGE 4–12% bis–tris midi gel (Thermo Fisher Scientific,), followed by the transfer of proteins onto nitrocellulose membranes (EMD Millipore). The nitrocellulose membrane was split and each were blocked for 1 h prior to incubation with the primary antibodies using 5% Milk TBST. The membranes were probed for Brd4 (Abcam, Ab128874, 1:1000), HiBiT monoclonal antibody (N7200) and left overnight to incubate at 4°C overnight on an orbital shaker. The third blot was probed using the one-step chemiluminescent Nano-Glo® HiBiT blotting system - following manufactures instructions - and was imaged a Bio-Rad imager (LI-COR Biosciences). Following overnight incubation with the primary antibodies at 4 °C, the membranes were incubated with secondary antibodies (anti-rabbit, Abcam AB216773, 1:5000 or anti-mouse, Abcam AB216774, 1:5000) and hFABTM rhodamine anti-tubulin antibody (Biorad, 12004165, 1:10,000) for 1 h and then imaged with a Bio-Rad imager (LI-

COR Biosciences). All western blots were analysed for band intensities using Image Lab from Bio-Rad (LI-COR, Biosciences). Using this validation approach it was determined that this clone contained a heterozygous insert of HiBiT-BromoCatch-Brd4 and will subsequently be referred to as an endogenous BromoCatch-Brd4 HEK293 for all subsequent experiments using this cell line.

**HiBiT live-cell kinetic degradation assay of MR170**

The validated endogenous BromoCathc-Brd4 cell line was subsequently used to perform a live-cell kinetic degradation assay to determine the activity of the functionalised degrader probes for BromoCatch MR170. BromoCatch-Brd4 HEK293 cells were plated at a density of $1x10^6$ cells into a single well of a 6-well plate in 2 mL of DMEM:10%FBS and left to adhere for 16 hours in a humidified incubator with 5% CO2 at 37 °C. The BromoCatch-Brd4 HEK293 cells were then transfected with 2 µg of pCMV LgBIT expression vector (Promega, N2681) using Fugene HD at a ratio of 1:3 DNA: transfection reagent. The DNA: transfection reagent mixture was incubated for 20 minutes at room temperature and then subsequently added dropwise to the BromoCatch-Brd4 HEK293 cells and left to incubate in a humidified incubator with 5% CO2 at 37 °C overnight.

The following day, the transfected BromoCatch-Brd4 HEK293 cells were trypsinised and plated at a density of $2x10^4$ cells per well into a non-adherent white walled 96 well plate in 100 µL of DMEM containing 20 µM Endurazine (Promega) and plates were incubated at 37 °C, 5% CO2, for 2 h. The plate was then imaged for luminescence using a GloMax Discover (Promega) imager as a baseline, then subsequently addition of a range of concentrations of between 25-2.5 µM of-MR170 or 10µM-1µM MZ1 or AGB1 was added. A breathable film was placed over the plate and continuously imaged every ~5 min for a period of 9 h on a GloMax Discover (Promega) set to 37 °C. The data produced from this experiment was subsequently analysed in graphpad Prism 10.2.3 using a one phase decay model to extract the DMAX and $T^{1/2}$ degradation of each bifunctional compound.

**HiBiT lytic degradation assay of MZ1, AGB1 and C10852S**

Our endogenous BromoCatch Brd4 HEK293 cell line along with endogenous Brd4, Brd3 and Brd2-HiBiT lines were plated at a density of $2x10^4$ cells per well of a white-walled 96 well plate 16 hours prior to initiation of experiment to enable adherence. The endogenous Brd4, Brd3 and Brd2 were designed as previously published from our group. At initiation of the experiment cells were incubated in DMEM:10% FBS containing DMSO (vehicle), MZ1, AGB1 or C10852S at a concentration range of 10µM to 0.1nM. Cells were left incubate for 4 hours at 37 °C, 5% CO2, and 95% humidity. At the assay endpoint, cells were treated with HiBiT lytic reagent following manufacture instructions. The plates were then imaged for luminescence on a BMG Labtech PHERAstar plate reader. The data produced from this experiment was subsequently analysed in graphpadPrism 10.2.3, with degradation curves being produced using a "log(inhibitor) vs. response (three parameters)" model to extract the $DC_{50}$ nM for each bifunctional normalised to the DMSO (vehicle) control.

**NanoBRET competition ligand binding assay of electrophilic warheads**

In order to perform the NanoBRET competition assay, HEK293 cells were first transfected with the pCMV_Brd4-BD2$^{L387A, E438C}$-Nanoluciferase vector, described above. Two days prior to initiation of the experiment 1 mL of HEK293 cells were plated at a density of $2x10^6$ cells per mL into one well of a 6-well plate and the cells were incubated at 37 °C, 5% CO2 overnight. To perform the transfection, firstly 2 µg of the Brd4-BD2$^{L387A, E438C}$-Nanoluciferase vector was added to 100 µL of Optimem media and 6 µL of Fugene HD was added for a 1:3 DNA:Fugene HD ratio and gently mixed and left at room temperature for 20 minutes. After this incubation period, the mixture was added dropwise onto the HEK293 cells plated the day prior and then left to incubate in a humidified incubator with 5% CO2 at 37 °C for at least 16 hours prior to initiation of experimentation.

For the live cell mode of NanoBRET, the HEK293 cells transfected with the Brd4-BD2$^{L387A, E438C}$-Nanoluciferase vector, and were subsequently trypsinised following standard procedure, counted and plated in Optimem media - containing 10% FBS – at a density of $2.36x10^5$ cells per mL in 10 mL of media. 85 µL of this cell containing media was then subsequently plated into every well of a non-adherent white walled 96 well plate to have a final density of ~$2x10^4$ cells per well. 5µL of optimem media containing 20µM ET-JQ1-Bodipy probe was added to each well and the plate was incubated for 5 minutes at room temperature on a plate shaker. Following this 10µL of a 10x solution of electrophilic warheads in optimem was added to the cells with concentration range from 100 µM – 2pM with ET-JQ1-OMe and DMSO (Vehicle) used as positive controls. The plate was subsequently incubated at 5% CO2 at 37 °C for 10 minutes. A 50µL 3x solution of NanoBRET Nano-Glo® Substrate + extracellular Nanoluciferase inhibitor was added to each experimental condition and subsequently imaged on a PHERAstar plate reader for both luminescent and fluorescence using emission (460nm) and acceptor emission (618nm). The relative BRET ratio was calculated by normalizing the data. The data produced from this experiment was subsequently analysed in graphpadPrism 10.2.3, with competitive binding curves being produced using a "log(inhibitor) vs. response (three parameters)" model to extract the $IC_{50}$ nM for each electrophilic warhead normalised to the DMSO (vehicle) control.

**NanoBRET Residence time of electrophilic warheads**

In order to perform the NanoBRET residence time assay, HEK293 cells were first transfected with the pCMV_Brd4-BD2$^{L387A, E438C}$-Nanoluciferase vector, described above. Two days prior to initiation of the experiment 1 mL of HEK293 cells were plated at a density of $2x10^6$ cells per mL into one well of a 6-well plate and the cells were incubated at 37 °C, 5% CO2 overnight. To perform the transfection, firstly 2 µg of the Brd4-BD2$^{L387A, E438C}$-Nanoluciferase vector was added to 100 µL of Optimem media and 6 µL of Fugene HD was added for a 1:3 DNA:Fugene HD ratio and gently mixed and left at room temperature for 20 minutes. After this incubation period, the mixture was added dropwise onto the HEK293 cells plated the day prior and then left to incubate in a humidified incubator with 5% CO2 at 37 °C for at least 16 hours prior to initiation of experimentation.

The HEK293 cells transfected with the Brd4-BD2$^{L387A, E438C}$-Nanoluciferase vector, were subsequently trypsinised following standard procedure, counted and plated in Optimem media - containing 10% FBS – at a density of 2 $x10^5$ cells per mL in 1 mL of media into 5 separate 15 mL conical tubes. These tubes of cells were then treated with a saturating concentration of 250 nM of MR112, MR116, 250 nM ET-JQ1-OMe, equivalent DMSO and a non-treatment control. These flacons were then left to equilibrate for 2 hours in a humidified incubator with 5% CO2 at 37 °C. After the 2 hours, cells were spun down and washed with warm optimem: 10% FBS containing media, this was performed 3 times. Cells were then plated at a density of $2x10^4$ cells in 90 µL per well of a white-walled 96 well plate. 100µl of 2X NanoBRET™ Nano-Glo® Substrate plus Extracellular NanoLuc® Inhibitor Solution was added to each well of the 96-well plate. The plate was then imaged on a GloMax Discover (Promega) plate reader for both luminescent and fluorescence using emission (460nm) and acceptor emission (618nm) as a baseline measurement. 10µL of a 20x ET-JQ1-BODIPY tracer final concentration of 25µM was added to each well and quickly shaken for 10 seconds on a plate shaker. A breathable film was then placed on the plate and the cells were continuously imaged for 120 minutes at 37 °C. The data produced from this experiment was subsequently analysed in graphpadPrism 10.2.3, with BRET curves being produced using a "One-phase decay" model normalised to the DMSO (vehicle) and non-treatment controls.

**Confocal live cell imaging of U2-O S, HEK293 FT using C10852S**

To perform confocal live cell imaging of C10852S, U2-O S were first transfected with the pCMV_H2B-BromoCatch vector, based on the smaller 15 kDa BromoCatch tag. Two days prior to initiation of the experiment 1 mL of cells were plated at a density of $2x10^6$ cells per mL into one well of a 6-well plate and the cells were incubated at 37 °C, 5% CO2 overnight. To perform the transfection, firstly 2 µg of the H2B-BromoCatch vector was added to 100 µL of Optimem media and 6 µL of Fugene HD was added for a 1:3 DNA:Fugene HD ratio and gently mixed and left at room temperature for 20 minutes. After this incubation period, the mixture was added dropwise onto the cells plated the day prior and then left to incubate in a humidified incubator with 5% CO2 at 37 °C for at least 16 hours prior to initiation of experimentation. In parallel $2x10^6$ cells of HEK293FT and U2-O S were plated but not transfected with the H2B-BromoCatch vector to act as a control.

These transfected cells were then subsequently trypsinised and plated at a density of $4x10^4$ cells per well of an ibidi μ-slide 18 Well chamber microscopy slide and left to adhere for 16 hours. Following adherence, the DMEM media was replaced with Optimem:10% FBS containing either DMSO, 200nM C10852S left to incubate for upto 8 hours in the transfected cells in a humidified incubator with 5% CO2 at 37 °C . The cells were then washed gently in warm Optimem:10% FBS media three times and Hoescht 33342 was added to each well 30 minutes prior to imaging at manufactures specified concentration. Confocal microscopy images were acquired on a Zeiss 880 Airyscan Confocal Microscope using Plan-Apochromat 63x/1.4 objective lens and equipped with 488/561/633-nm excitation laser lines. Post-acquisition analysis was conducted with Fiji (version 1.54j).

To perform confocal live cell imaging of C10852S using the pCMV_H2B-BromoCatch vector, based on the smaller 12 kDa Bromocatch tag. U2-O S and HEK293FT cells were first transfected with the pCMV_H2B-BromoCatch vector. Two days prior to initiation of the experiment 1 mL of cells were plated at a density of $2x10^6$ cells per mL into one well of a 6-well plate and the cells were incubated at 37 °C, 5% CO2 overnight. To perform the transfection, firstly 2 μg of the H2B-BromoCatch vector was added to 100 μL of Optimem media and 6 μL of Fugene HD was added for a 1:3 DNA:Fugene HD ratio and gently mixed and left at room temperature for 20 minutes. After this incubation period, the mixture was added dropwise onto the cells plated the day prior and then left to incubate in a humidified incubator with 5% CO2 at 37 °C for at least 16 hours prior to initiation of experimentation. In parallel $2x10^6$ cells of HEK293FT and U2-O S were plated but not transfected with the H2B-BromoCatch vector to act as a control.

Both these transfected and un-transfected cells were then subsequently trypsinised and plated at a density of $4x10^4$ cells per well of an ibidi μ-slide 18 Well chamber microscopy slide and left to adhere for 16 hours. Following adherence, the DMEM media was replaced with Optimem:10% FBS containing either DMSO, 100nM C10852S or 100nM C10852S + 25μM ET-JQ1-OME and left to incubate for 6 hours in HEK293FT or 8 hours in U2OS in the transfected cells in a humidified incubator with 5% CO2 at 37 °C . In the un-transfected control 100nM C100852S in optimem: 10% FBS was added and left in a humidified incubator with 5% CO2 at 37 °C for the same length as the transfected cells. The cells were then washed gently in warm Optimem:10% FBS media three times and Hoescht 33342 was added to each well 30 minutes prior to imaging at manufactures specified concentration. Confocal microscopy images were acquired on a Zeiss 880 Airyscan Confocal Microscope using Plan-Apochromat 63x/1.4 objective lens and equipped with 488/561/633-nm excitation laser lines. Post-acquisition analysis was conducted with Fiji (version 1.54j).

**Cell harvesting and lysis**

To harvest adherent HEK293FT cells, the medium was removed, and the cells washed with rt PBS. Cells were detached by incubating with RIPA buffer (1% Triton X-100, 0.1% SDS, and 1:200 Protease inhibitor cocktail) and incubating at 4 °C for 15 min. The lysate was centrifuged (13,000 ×g, 4 °C, 10 min) to pellet cell debris, and the resulting supernatant was transferred to fresh Eppendorf tubes.

**Western blotting**

Proteins in the electrophoresed gels were transferred onto a nitrocellulose membrane for 90 min at 90 V, at 4 °C. The primary antibodies were diluted in 5% milk in TBS-Tween (TBS-T, 0.1% Tween). The membrane was incubated with the primary antibodies overnight at 4 °C. The membrane was washed with TBS-T (3 × 5 min), and then treated with the appropriate secondary antibody diluted in 5% milk in TBS-Tween (TBS-T, 0.1% Tween), and then incubated for 1 h at rt. The membrane was washed with TBS-T (3 × 5 min) before being imaged on a Bio-rad molecular imager.

**Sodium dodecyl sulfate polyacrylamide gel electrophoresis of lysates**

Protein samples were separated by gel electrophoresis on NuPAGE™ 4–12%, Bis-Tris precast polyacrylamide gels (Thermofisher) unless otherwise stated. Protein or cell lysate proteins were separated by SDS-PAGE gel electrophoresis. Samples (10 uL) were prepared addition of LDS/DTT loading buffer (10 uL, 4×LDS sample buffer, 2% DTT) and then heated at 95 °C for 5 min. Protein samples (15–28 µL) were loaded into NuPAGE 4-12% Bis-Tris Gels and electrophoresed for 45 min at 180V in MES buffer. The Protein Standards ladder (Bio-rad) (5 µL) was used as a molecular weight reference. Following electrophoresis, gels were imaged with ChemiDoc MP Imaging System for fluorescence read-out at the appropriate wavelength (for fluorescent labelling experiments) or stained for 15 min with Instant Blue Coomasie® stain (Abcam), destained, and imaged using Bio-rad molecular imager. Bio-rad Image Lab 6.1 was used to process the data. Protein concentrations were measured prior to use in assays using a NanoDrop Microvolume Spectrophotometer.

**Determination of total protein concentration**

Total protein concentration of the lysate was determined by a BCA protein assay (Thermo Scientific) and the absorbance at 562 nm measured and plotted using a PHERAStar plate reader. Cell lysates were diluted to 2 mg/mL with miliQ water.

**Crystallization of Brd2-BD2$^{L383A,D434C}$**

For crystallization, the 1.2 mM stock of Brd2-BD2$^{L383A,D434C}$ was diluted into crystallization buffer (25 mM HEPES, 120 mM NaCl, 0.5 mM TCEP, pH 7.5) at a final concentration of 100 µM in 1 ml. From a 10 mM stock, 12 µl of MR116 was added in 2 µl increments with gentle mixing in-between reach a final concentration of 120 µM (1.2-fold molar excess). Brd2-BD2$^{L383A,D434C}$ and MR116 were allowed to react at ambient temperature for 120 min. The reaction was spun at top speed in a microcentrifuge for 10 minutes and the 1ml of supernatant was injected to a 5 ml sample look and loaded on a 16/600 Superdex s75 pg size exclusion column (SEC) pre-equilibrated in crystallization buffer. The run was carried out on a BioRad NGC equipped with a multi-wavelength detector and monitored at 215 nm, 255 nm, and 280 nm. Fractions with UV absorbance at 280 nm were analyzed using NuPAGE Bis-Tris mini protein 4-12% gels (Invitrogen) with MES running buffer ran at 200 V for 34 min. The gel was stained using colloidal Coomassie Instant Blue Stain (ISB1L, abcam), revealing pure BRD2-BD2$^{L383A,D434C}$. The additional UV absorbance at 215 nm suggested the presence of MR116, which was confirmed from the $\Delta T_m$ over 25 °C ($T_m$ Brd2-BD2$^{L383A,D434C}$ = 45.37 °C and $T_m$ MR116/Brd2-BD2$^{L383A,D434C}$ = 71.72 °C) using nanoDSF (20 to 95 °C, at 1°C/min) on the Prometheus Panta (Nanotemper). With this verification, MR116/Brd2-BD2$^{L383A,D434C}$ was concentrated to 610 µM (8.2 mg/ml) using a 3,000 MWCO centrifuge concentrator (Amicon). Because MR116 did not exhibit UV-absorbance at 280 nm on the SEC run, the concentration was determined using the molar extinction coefficient at 280 nm($\varepsilon_{280}$) = 15,930 cm$^{-1}$*M$^{-1}$. Aliquots of MR116/Brd2-BD2$^{L383A,D434C}$ were stored at -80 °C until use.

Initial crystal screens of MR-116/Brd2-BD2$^{L383A,D434C}$ (8.2 mg/ml) were set against commercially available screens (JSCG+, PACT, ProPLEX, Morpheus-I, PGA, Index-HT, PEGion, MIDAS, Classics, and BCS) in 96-well MRC 2-drop sitting well plates with 50 µl reservoirs, using a Mosquito (SPT LabTech) dispensing 200 nl of reservoir and 200 nl of MR-116/Brd2-BD2$^{L383A,D434C}$ for each drop. Plates were sealed and crystals were monitored at 19 °C in a Rock Imager (Formulatrix). Numerous conditions yielded suitable crystals between 1-3 days. Crystals were harvested with Dual Thickness MicroLoops (MiTeGen) ranging from 50-200 µm. Our reported structure was obtained from a single ~100 µm crystal after 2 days in the Morpheus-I condition E9 (PEG 500 MME 40% (v/v), PEG 20,000 20 % (w/v), 0.12 M ethylene glycols, 0.1 M Tris-BICINE pH 8.5) and was harvested without additional cryoprotection.

**Macromolecular X-ray Data collection**

Data sets for all crystals were collected at Diamond Light Source (Didcot, UK) on MX beamline I24 under bag proposal MX-35324-32. A total of 3,600 images were recorded on the EIGER2 X CdTe 9M detector positioned at a distance corresponding to a maximum resolution of 1.1 Å, total exposure time of 0.005 sec/image, 0.1° oscillation over 360°, wavelength of 0.6199 Å, and beam size of 20x20 µm. Data sets were automatically processed SynchWeb/ISPyB for indexing (XDS), scaling, and merging.

The crystal was processed in space group P 21 21 2 with unit cell dimensions of 32.05 Å, 52.12 Å, 71.64 Å in length and angles of 90° 90° 90°. We estimate that this crystal achieved diffraction at the limit of the beamline and detector, but the highest resolution shells were incomplete. Therefore, we reprocessed the images with a resolution cutoff of 1.3 Å to obtain a complete dataset for structural determination.

**Structure determination of MR116/Brd2-BD2$^{L383A,D434C}$**

To facilitate molecular replacement, we generated an atomic model of Brd2-BD2$^{L383A,D434C}$ without MR116 using Boltz-1[1] and automatic MSA generation[2]. Crystallographic analysis was carried out in CCP4 v8[3]. Initial inspection of the crystal had a Mathews coefficient of 2.28 and high probability of 1 copy of Brd2-BD2$^{L383A,D434C}$ per unit cell with a solvent content of 46 %. Running CCP4 pipeline (Data reduction to complete structure with ligand fitting) with Aimless for data reduction, Phaser for molecular replacement using the Blotz-I model, Buccaneer autobuild, and REFMAC5. This initial attempt resulted with structure factors of $R_{work}$=0.20 and $R_{free}$=0.23 and clear density for MR116. The SMILES string with the saturated $CH_2$-$CH_3$ analogue of the MR116 acrylamide was used to generate restrains for the ligand and for subsequent linkage with the right carbon valency. The covalent bond was formed using coot, between the beta $CH_3$ carbon in the ligand and the sulfur in C434 of Brd2-BD2$^{L383A,D434C}$ in CPP4 and the hydrogen removed manually. This was used as a covalent restraint for further refinement. After several rounds of manual model building in COOT and REFMAC, the final structure factors were $R_{free}$= 0.16 and $R_{work}$= 0.15 for the predominant bond conformation (see Table SX). An alternate conformation of the covalent bond was built and further refined with REFMAC5 to give a final $R_{free}$= 0.16 and $R_{work}$= 0.15 (See Supplementary Figure S). The polder map (omit map) was prepare in PHENIX omitting the ligand density MR116 and the cysteine and then visualized in UCSF ChimeraX (Supplementary Figure 9).

**Supplementary Table 9. Data collection and refinement statistics**

| | **PDB 9QRK** |
|---|---|
| **Data collection** | |
| Space group | P 21 21 2 |
| Cell dimensions | |
| *a*, *b*, *c* (Å) | 32.05, 52.13, 71.64 |
| α, β, γ (°) | 90, 90, 90 |
| Resolution (Å) | 35.85 – 1.30 (1.32-1.30)* |
| $R_{merge}$ | 0.035 (0.096) |
| $(I / I_{SD})$ | 43.6 (20.7) |
| Completeness (%) | 99.9 (99.2) |
| Redundancy | 13.2 (13.2) |
| **Refinement** | |
| Resolution (Å) | 35.85 – 1.30 |
| Number of reflections/free | 30269 (1503) |
| $R_{work}$ / $R_{free}$ | 0.147 / 0.163 |
| No. atoms | |
| Protein | 1987 |
| Ligand/ion | 134 |
| Water | 161 |
| *B*-factors | |
| Protein | 11.37 |
| Ligand/ion | 11.51 |
| Water | 21.72 |
| R.m.s. deviations | |
| Bond lengths (Å) | 0.0144 |
| Bond angles (°) | 2.08 |

*Values in parentheses are for highest-resolution shell.

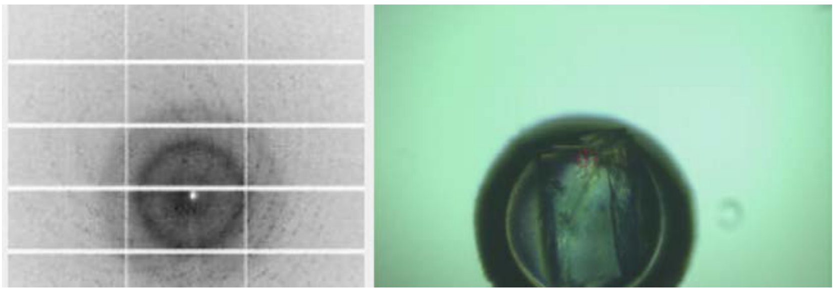

**Chemistry**

Unless otherwise stated, all reagents and solvents were purchased from commercial sources and used without further purification. Nuclear magnetic resonance spectra were recorded on a Bruker Ascend 500 MHz spectrometer or a Bruker Avance III HD spectrometer, operating at 500 MHz and 400 MHz for $^{1}$H NMR, respectively, 100 MHz for $^{13}$C NMR, and 376 MHz for $^{19}$F NMR. $^{1}$H NMR and $^{13}$C NMR chemical shifts (δ) are reported in parts per million (ppm) and are referenced to residual protium in solvent and to the carbon resonances of the residual solvent peak respectively. DEPT and correlation spectra were run in conjunction to aid assignment. $^{19}$F NMR chemical shifts are reported in ppm and are uncorrected. Coupling constants (*J*) are quoted in Hertz (Hz), and the following abbreviations were used to report multiplicity: s= singlet, d= doublet, dd= doublet of doublets, ddd= double doublet of doublets, t= triplet, dtt= double triplet of triplets, q= quartet, m= multiplet, br s= broad singlet. Liquid chromatography-mass spectrometry (LC-MS) was carried out on a Shimadzu HPLC/MS 2020 equipped with a Hypersil Gold column (1.9 µm particle size, 50 × 2.1 mm), photodiode array detector and ESI detector, or by using Agilent InfinityLab LC/MSD systems. Purification by flash column chromatography was carried out using Fisher Scientific silica gel 60Å (35-70 µm), or by using Biotage Selekt, Biotage Isolera, Grace Reveleris, Buchi Pure, or Teledyne Isco Combiflash systems. Thin layer chromatography was performed on glass plates pre-coated with silica gel (Analtech, UNIPLATE™ 250 µm / UV254), with visualization being achieved using UV light (254 nm) and/or by staining with alkaline potassium permanganate dip. High-resolution mass spectral (HRMS) data were collected in the laboratories of the University of Bath Chemistry Department using an Agilent 6545 LC/Q-TOF system or at the University of Dundee on a Bruker MicroTOF II focus ESI mass spectrometer connected in parallel to a Dionex Ultimate 3000 RSLC system with a diode array detector and a Waters XBridge C18 column (50 mm × 2.1 mm, 3.5 µm particle size).

**Methyl (*R*)-2-((*S*)-2,3,9-trimethyl-4-(4-vinylphenyl)-6*H*-thieno[3,2-*f*][1,2,4]triazolo[4,3-*a*][1,4]diazepin-6-yl)butanoate (MR100)**

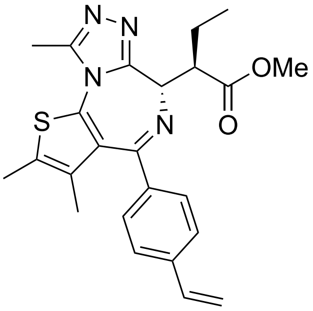

A solution of methyl (*R*)-2-((*S*)-4-(4-chlorophenyl)-2,3,9-trimethyl-6*H*-thieno[3,2-*f*][1,2,4]triazolo[4,3-*a*][1,4]diazepin-6-yl)butanoate ((+)-ET-JQ1-OMe bump, OMe) 40 mg, 0.09 mmol) , potassium vinyltrifluoroborate (36 mg, 0.27 mmol) , XPhos Pd G2 (21 mg, 0.03 mmol) and *N,N*-diisopropylethylamine (0.08 mL, 0.45 mmol) in DMF (0.6 mL) and water (0.06 mL) was stirred at 100 °C for 24 hours. Upon cooling to ambient temperature, the mixture was filtered through diatomaceous earth, the filtrate was concentrated under reduced pressure and the residue was and redissolved in dichloromethane. The solution was washed with water (2 x 50 mL) and brine (20 x 50 mL), dried over anhydrous magnesium sulfate and concentrated under reduced pressure. Purification by flash column chromatography, eluting with 0-5% methanol/dichloromethane, afforded the title compound as a yellow solid (35 mg, quant.). $^1$H NMR (500 MHz, $CDCl_3$) δ 7.32 – 7.24 (m, 4H), 6.64 (dd, *J* = 17.6, 10.9 Hz, 1H), 5.72 (d, *J* = 17.6, 1H), 5.24 (d, *J* = 10.9 Hz, 1H), 4.17 (d, *J* = 10.9 Hz, 1H), 3.93 (td, *J* = 10.8, 3.6 Hz, 1H), 3.79 (s, 3H), 2.60 (s, 3H), 2.34 (s, 3H), 2.16-2.07 (m, 1H), 1.64-1.55 (m, 4H), 0.96 (t, *J* = 7.4 Hz, 3H). $^{13}$C NMR (126 MHz, $CDCl_3$) δ 175.41, 163.14, 154.42, 149.73, 143.51, 140.90, 136.75, 136.53, 132.12, 130.90, 130.80, 130.42, 129.81, 128.73, 128.68, 59.38, 51.57, 49.65, 23.24, 14.48, 13.14, 11.87, 11.65. m/z (ES+): 435.0 $[M+H^+]^+$. HRMS (ES+) calculated for $[(C24H27O2N4S)+H^+]^+$ 435.18492 found 435.18550

**Methyl (*R*)-2-((*S*)-4-(4-formylphenyl)-2,3,9-trimethyl-6*H*-thieno[3,2-*f*][1,2,4]triazolo[4,3-*a*][1,4] diazepin-6-yl)butanoate (MR70)**

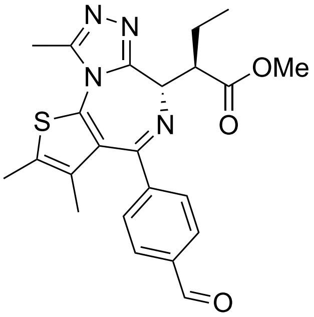

To a stirred solution of methyl (*R*)-2-((*S*)-2,3,9-trimethyl-4-(4-vinylphenyl)-6*H*-thieno[3,2-*f*][1,2,4]triazolo[4,3-*a*][1,4]diazepin-6-yl)butanoate (MR100, 20 mg, 0.05 mmol) and sodium periodate (30 mg, 0.14 mmol) in acetone (0.5 mL) and water (0.1 mL) was added osmium tetroxide (15 µL, 4%w/v solution in water). After stirring for 2 hours, the reaction mixture was diluted with EtOAc. The solution was washed with water, and the organics were dried over anhydrous magnesium sulfate and concentrated under reduced pressure. Purification by preparative reversed phase column chromatography, eluting with 5/95 to 95/5 MeCN/$H_2O$ (0.1 % $NH_4OH$) afforded the title compound as a white solid (18 mg, quant). $^1$H NMR (500 MHz, $CDCl_3$) δ 10.07 (s, 1H), 7.88 (d, *J* = 8.5 Hz, 2H), 7.57 (d, *J* = 8.1 Hz, 2H), 4.32 (d, *J* = 11.0 Hz, 1H), 4.05 (td, *J* = 10.7, 3.7 Hz, 1H), 3.90 (s, 3H), 2.71 (s, 3H), 2.45 (s, 3H), 2.26 – 2.18 (m, 1H), 1.77 – 1.66 (m, 4H), 1.06 (t, *J* = 7.4 Hz, 3H); m/z (ES+): 437.0 $[M+H^+]^+$. HRMS (ES+) calculated for $[(C23H25O2N4S)+H^+]^+$ 437.16419 found 435.16515

**Methyl (*R*)-2-((*S*)-4-(4-(3-aminoprop-1-yn-1-yl)phenyl)-2,3,9-trimethyl-6*H*-thieno[3,2-*f*][1,2,4] triazolo [4,3-*a*][1,4]diazepin-6-yl)butanoate (MR101)**

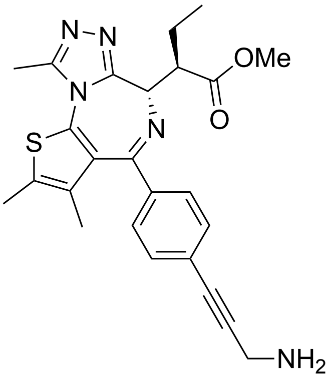


To a stirred solution of methyl (*R*)-2-((*S*)-4-(4-chlorophenyl)-2,3,9-trimethyl-6*H*-thieno[3,2-*f*][1,2,4]triazolo[4,3-*a*][1,4]diazepin-6-yl)butanoate (70 mg, 0.16 mmol) in THF (1.5 mL) was added XPhos Pd G2 (37. 3mg, 0,3 eq. 0.05 mmol), XPhos (23 mg, 0.05 mmol), cesium carbonate (258 mg, 0.79 mmol) and propargyl amine (0.10 mL, 1.58 mmol).The reaction was stirred for 3 h at 90 °C, then cooled to 60 °C and stirred overnight. Upon cooling to ambient temperature, the reaction mixture was suspended in dichloromethane and filtered through diatomaceous earth. The filtrate was washed with water, and the organic phase was dried over anhydrous magnesium sulfate and concentrated under reduced pressure. Purification by preparative reversed phase column chromatography, eluting with 5/95 to 95/5 MeCN/$H_2O$ (0.1 % formic acid), afforded the title compound as a white solid (3 mg, 20%). $^1$H NMR (500 MHz, $CDCl_3$) δ 7.36 (d, J= 8.2 Hz, 2H), 7.31 (d, J= 8.0 Hz, 2H), 4.23 (d, *J* = 10.9 Hz, 1H), 3.98 (td, *J* = 10.7, 3.6 Hz, 1H), 3.84 (s, 3H), 3.65 (s, 2H**)**, 2.65 (s, 3H), 2.40 (s, 3H), 2.20 – 2.13 (m, 1H), 1.71 – 1.60 (m, 4H), 1.01 (t, *J* = 7.4 Hz, 3H); m/z (ES+): 462.1 $[M+H^+]^+$. HRMS (ES+) calculated for $[(C25H28O2N5S)+H^+]^+$ 462.19582 found 462.19632

**Methyl 2-((6*S*)-2,3,9-trimethyl-4-(4-(oxiran-2-yl)phenyl)-6*H*-thieno[3,2-*f*][1,2,4]triazolo[4,3-*a*][1,4]diazepin-6-yl)butanoate (MR104)**

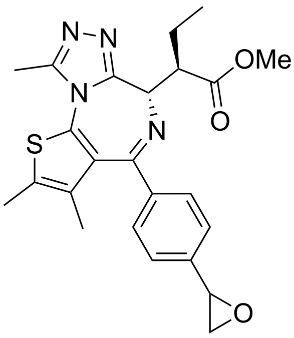


To a stirred solution of methyl 2-[(9S)-4,5,13-trimethyl-7-(4-vinylphenyl)-3-thia-1,8,11,12-tetrazatricyclo[8.3.0.02,6]trideca-2(6),4,7,10,12-pentaen-9-yl]acetate (MR100, 30 mg, 0.05 mmol) in dichloromethane (1 mL) was added m-chloroperoxybenzoic acid (12 mg) and the reaction was stirred overnight. The reaction mixture was diluted with dichloromethane and then washed with water (2 x 50 mL) and brine (2 x 50 mL). The organic phase was dried over anhydrous magnesium sulfate and concentrated under reduced pressure. Purification by preparative reversed phase column chromatography, eluting with 5/95 to 95/5 MeCN/$H_2O$ (0.1 % $NH_4OH$), afforded the title compound as a white solid (3 mg, 10%). $^1$H NMR (500 MHz, $CDCl_3$) δ 7.40 – 7.32 (m, 3H), 7.32 (d, *J* = 8.5 Hz, 1H), 7.23 (d, *J* = 7.6 Hz, 2H), 4.23 (d, *J* = 11.0 Hz, 1H), 3.98 (td, *J* = 10.9, 3.6 Hz, 1H), 3.86 – 3.85 (m, 1H), 3.84 (s, 3H), 3.16 – 3.13 (m, 1H), 2.74 (td, *J* = 5.8, 2.5 Hz, 1H), 2.69 (s, 1H), 2.66 (d, *J* = 2.9 Hz, 3H),

2.40 (s, 3H), 2.21 – 2.13 (m, 1H), 1.71 – 1.59 (m, 5H), 1.01 (t, *J* = 7.4 Hz, 3H); m/z (ES+): 451.0 $[M+H^+]^+$. HRMS (ES+) calculated for $[(C24H26N4O3S)+H^+]^+$ 451.17984 found 451.18087

**Methyl (*R*)-2-((*S*)-4-(4-aminophenyl)-2,3,9-trimethyl-6H-thieno[3,2-f][1,2,4]triazolo[4,3a][1,4] diazepin-6-yl)butanoate (Compound 7)**

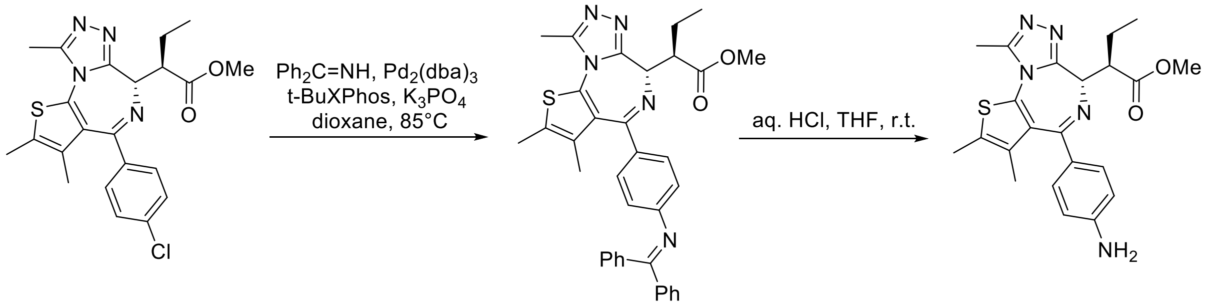


To a stirred solution of methyl (*R*)-2-((*S*)-4-(4-chlorophenyl)-2,3,9-trimethyl-6H-thieno[3,2-f][1,2,4]triazolo[4,3-a][1,4]diazepin-6-yl)butanoate (2.50 g, 5.64 mmol) in 1,4-dioxane (125 mL) was added benzophenone imine (1.22 g, 6.77 mmol) and potassium phosphate tribasic (2.99 g, 14.11 mmol), and argon (g) was bubbled through the stirring mixture for 30 minutes. After this time, *t*-BuXPhos (0.36 g, 0.85mmol) and $Pd_2(dba)_3$ (0.26 g, 0.28 mmol) were added, and after passage of argon (g) for a further 5 minutes, the reaction mixture was heated at 85 °C overnight. The reaction mixture was cooled to 55 °C and partially concentrated under reduced pressure, the subsequent residue being partitioned between water and ethyl acetate. The organic phase was separated, and the aqueous component was extracted with ethyl acetate. The combined organic extracts were washed with brine, dried over anhydrous magnesium sulfate and concentrated under reduced pressure to a brown oil. The crude imine was dissolved in THF (50 mL) and treated with 1M hydrochloric acid (50 mL), and the resulting solution was stirred at ambient temperature for 2 hours before being partially concentrated under reduced pressure. The aqueous residue was further diluted with water and washed with ethyl acetate. The aqueous phase was then adjusted to pH8-9 by slow addition of saturated aqueous $NaHCO_3$ solution and extracted with ethyl acetate (3 portions). These organic extracts were combined, dried over anhydrous magnesium sulfate and concentrated under reduced pressure to a pale-yellow foam. Purification by flash column chromatography, eluting with 1-2% 7M methanolic ammonia/dichloromethane, afforded a yellow foam. Lyophilisation from MeCN/$H_2O$ (1:2) afforded the title compound as a pale-yellow solid (1.20 g, 50%). $^1H$ NMR (400 MHz, $CDCl_3$) δ: 7.19 (d, *J*= 8.3 Hz, 2H), 6.58 (d, *J*= 8.8 Hz, 2H), 4.18 (d, *J*= 11.0 Hz, 1H), 3.97 (td, *J*= 10.7, 3.7 Hz, 1H), 3.83 (s, 3H), 2.64 (s, 3H), 2.40 (s, 3H), 2.22-2.11 (m, 1H), 1.74-1.72 (m, 3H), 1.68-1.59 (m, 1H), 1.01 (t, *J*= 7.4 Hz, 3H); m/z (ES+): 446.2 $[M+Na^+]^+$

**Methyl (*R*)-2-((*S*)-4-(4-(2-chloroacetamido)phenyl)-2,3,9-trimethyl-6*H*-thieno[3,2-*f*][1,2,4]triazolo [4,3-*a*][1,4]diazepin-6-yl)butanoate (MR112)**

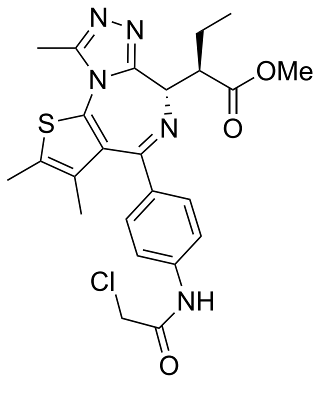


To a stirred solution of compound **7** (75 mg, 0.18 mmol) in dichloromethane (2 mL) at ambient temperature was added *N,N*-diisopropylethylamine (31 µL, 0.18 mmol), followed by the slow dropwise addition of 2-chloroacetyl chloride (043 µL, 0.53 mmol). After 2 hours, the reaction was quenched with a few drops of MeOH and concentrated under reduced pressure. The residue was partitioned between water and dichloromethane, and the organic phase was washed with water (2x50 mL) and brine (2x50 mL), dried over anhydrous magnesium sulfate and concentrated under reduced pressure, affording the title compound as a white solid (60 mg, 67%). $^{1}$H NMR (500 MHz, $CDCl_3$) δ 8.49 (s, 1H), 7.51 (d, *J* = 9.0 Hz, 2H), 7.31 (d, *J* = 8.6 Hz, 2H), 4.17 (d, *J* = 11.0 Hz, 1H), 4.12 (s, 2H), 3.91 (td, *J* = 10.7, 3.7 Hz, 1H), 3.78 (s, 3H), 2.60 (s, 3H), 2.34 (s, 3H), 2.13 – 2.05 (m, 1H), 1.65 – 1.53 (m, 4H), 0.95 (t, *J* = 7.4 Hz, 3H); $^{13}$C NMR (126 MHz, $CDCl_3$) δ 175.43, 164.05, 163.42, 154.55, 149.83, 139.01, 134.52, 131.77, 131.13, 130.82, 130.80, 129.50, 119.43, 59.30, 51.63, 49.73, 42.97, 23.25, 14.49, 13.17, 11.84, 11.69; m/z (ES+): 500.1 $[M+H^+]^+$; HRMS (ES+) calculated for $[(C24H26ClN5O3S)+H^+]^+$ 500.15176 found 500.15278

**Methyl (*R*)-2-((*S*)-2,3,9-trimethyl-4-(4-(vinylsulfonamido)phenyl)-6*H*-thieno[3,2-*f*][1,2,4]triazolo[4,3-*a*][1,4]diazepin-6-yl)butanoate (MR121)**

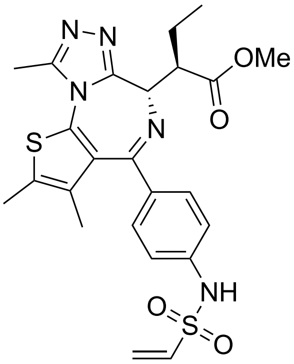


To a stirred solution of compound **7** (75 mg, 0.18 mmol) and pyridine (31 µL, 0.18 mmol) in dichloromethane (1.9 mL) at 0 °C was dropwise added vinyl sulfonyl chloride (17 µL, 0.18 mmol). The reaction mixture was stirred for 5 minutes before being quenched with a few drops of methanol and concentrated under reduced pressure. The residue was partitioned between dichloromethane and water, and the organic phase was washed with water (2x50 mL) and brine (2x50 mL), dried over anhydrous magnesium sulfate and concentrated under reduced pressure, affording the title compound as a white crystalline solid (20 mg, 30%). $^{1}$H NMR (500 MHz, $CDCl_3$) δ 7.45 (s, 1H), 7.35 (d, *J* = 8.2 Hz, 2H), 7.19 (d, *J* = 8.9 Hz, 2H), 6.53 (dd, *J* = 16.5, 9.9 Hz, 1H), 6.27 (d, *J* = 16.6 Hz, 1H), 5.95 (d, *J* = 9.9 Hz, 1H), 4.25 (d, *J* = 11.0 Hz, 1H), 3.99 (td, *J* = 10.7, 3.7 Hz, 1H), 3.86 (s, 3H), 2.69 (s, 3H), 2.43 (s, 3H), 2.24 – 2.12 (m, 1H), 1.74 – 1.61 (m, 4H), 1.04 (t, *J* = 7.4 Hz, 3H) $^{13}$C NMR (126 MHz, $CDCl_3$) δ 175.37, 163.34, 154.50, 149.83, 138.84, 134.99, 134.43, 131.89, 131.02, 130.82, 130.65, 129.80, 128.65, 119.54, 77.29, 77.03, 76.78, 59.30, 51.61, 49.77, 23.23, 14.45, 13.15, 11.81, 11.69; m/z (ES+): 514.0 $[M+H^+]^+$; HRMS (ES+) calculated for $[(C24H27N5O4S)+H^+]^+$ 514.15772 found 500.15910

**(*R*)-2-((*S*)-4-(4-Acrylamidophenyl)-2,3,9-trimethyl-6*H*-thieno[3,2-*f*][1,2,4]triazolo[4,3-*a*][1,4]diazepin-6-yl)butanoate (MR116)**

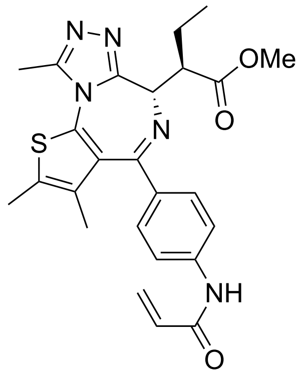

To a stirred solution of compound **7** (1 equiv., 75 mg, 0.18 mmol) and triethylamine (0.03 mL, 0.21 mmol) in dichloromethane (1.9 mL) at ambient temperature was dropwise added acryloyl chloride (0.021 mL, 0.27 mmol), and the reaction mixture was stirred for 30 minutes. The reaction mixture was diluted with dichloromethane and then washed with water (2 x 50 mL) and brine (2 x 50 mL), dried over anhydrous magnesium sulfate and concentrated under reduced pressure. Purification by flash column chromatography, eluting with 0-5% methanol/dichloromethane, afforded the title compound as a yellow solid (60 mg, 71 %). $^{1}$H NMR (500 MHz, $CDCl_3$) δ 8.46 (s, 1H), 7.58 (d, *J* = 8.3 Hz, 2H), 7.26 (d, *J* = 8.8 Hz, 2H), 6.36 (dd, *J* = 16.9, 1.6 Hz, 1H), 6.27 (dd, *J* = 16.8, 9.9 Hz, 1H), 5.65 (dd, *J* = 9.9, 1.7 Hz, 1H), 4.16 (d, *J* = 10.9 Hz, 1H), 3.89 (td, *J* = 10.8, 3.7 Hz, 1H), 3.76 (s, 3H), 2.58 (s, 3H), 2.33 (s, 3H), 2.12 – 2.04 (m, 1H) , 1.58 (m, 4H), 0.93 (t, *J* = 7.4 Hz, 3H); $^{13}$C NMR (126 MHz, $CDCl_3$) δ 175.42, 163.89, 163.61, 154.69, 149.83, 140.47, 133.59, 131.58, 131.27, 131.13, 130.99, 130.78, 129.38, 128.15, 119.41, 59.27, 51.64, 49.83, 23.26, 14.48, 13.16, 11.82, 11.70. ; m/z (ES+): 478.1 $[M+H^+]^+$. HRMS (ES+) calculated for $[(C25H27N5O3S)+H^+]^+$ 478.19074 found 478.19131

**(2-Amino-4,5-dimethylthiophen-3-yl)(3-chlorophenyl)methanone**

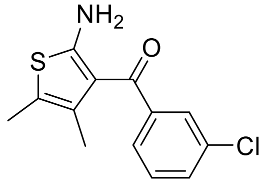

To a stirred suspension of 3-chlorobenzoylacetonitrile (40.00 g, 222.71 mmol) in ethanol (900 mL) was added methyl ethyl ketone (24.94 mL, 278.39 mmol), morpholine (3.90 mL, 44.54 mmol) and then sulfur (62.84 g, 244.98 mmol), and the resulting reaction mixture was stirred at 70 °C overnight. Upon cooling to ambient temperature, the reaction mixture was poured into brine (2500 mL) and extracted with ethyl acetate (3 x 1000 mL). The combined organic extracts were washed with brine (1000 mL), dried over anhydrous magnesium sulfate and concentrated under reduced pressure to an orange oil. Trituration with *tert*-butyl methyl ether overnight at ambient temperature afforded a yellow solid (16.91 g). Purification of the mother liquors by dry flash chromatography, eluting with 0-20% ethyl acetate/dichloromethane followed by trituration with *tert*-butyl methyl ether and petroleum ether (40:60) afforded a second crop of crude product (6.11 g). The combined material was converted to the oxalic acid salt by treatment with oxalic acid dihydrate (10.90 g, 86.6 mmol) in methanol (200 mL) and water (60 mL), and after concentration under reduced pressure, the salt was recrystallised from refluxing acetonitrile (330 mL) to give a crystalline solid (16.35 g). Treatment with 0.5M sodium hydroxide (aq.) (400 mL), dichloromethane (300mL) and *tert*-butyl methyl ether (400 mL) gave a biphasic mixture. The organic component was separated, and the aqueous phase extracted with *tert*-butyl methyl ether (2 x 200mL). The combined organic extracts were dried over anhydrous magnesium sulfate and concentrated under reduced pressure to afford the title compound as a green solid (13.59 g, 23%). $^1$H NMR (400 MHz, $CDCl_3$): δ 7.52-7.32 (m, 4H), 2.13 (d, *J*= 0.7 Hz, 3H), 1.54 (d, *J*= 0.7 Hz, 3H); m/z (ES+): 266.1 $[M+H^+]^+$.

**Methyl (*R*)-2-((*S*)-5-(3-chlorophenyl)-6,7-dimethyl-2-oxo-2,3-dihydro-1H-thieno[2,3-e][1,4]diazepin-3-yl)butanoate**

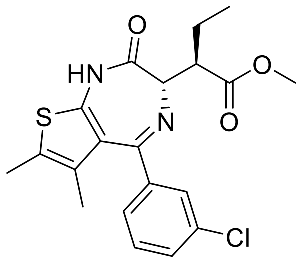

To a stirred suspension of (2-amino-4,5-dimethylthiophen-3-yl)(3-chlorophenyl)methanone (8.59 g, 32.31 mmol) in toluene (40 mL) at ambient temperature was added freshly dried and activated 4Å molecular sieves (30 g) followed by TFA (4.80 mL, 64.52 mmol). The resulting red solution was stirred for 5 minutes, after which time a solution of methyl (*R*)-2-((*S*)-2,5-dioxooxazolidin-4-yl)butanoate (6.50 g, 32.31 mmol) in toluene (20 mL) was added dropwise, and the reaction mixture was heated at 60 °C for 2.5 hours. Triethylamine (13.51 mL, 96.93 mmol) was added, and the reaction mixture was heated at 80 °C overnight. Upon cooling to ambient temperature, the reaction mixture was filtered, the filter cake being washed with dichloromethane, and the filtrate was concentrated under reduced pressure. The residue was partitioned between saturated aqueous sodium hydrogen carbonate solution and dichloromethane. The organic phase was separated, and the aqueous component was extracted with dichloromethane. The combined organic extracts were washed with brine, dried over anhydrous magnesium sulfate and concentrated under reduced pressure. Purification by flash column chromatography, eluting with 0-10% ethyl acetate/dichloromethane, and then a for a second time eluting with 2-10% ethyl acetate/cyclohexane, afforded the title compound as a yellow solid (5.20 g, 38%). $^1$H NMR (400 MHz, $CDCl_3$): δ 8.63 (br s, 1H), 7.42-7.36 (m, 2H), 7.29-7.22 (m, 2H), 3.83 (s, 3H), 3.71-3.61 (m, 1H), 2.30 (d, *J*= 0.5 Hz, 3H), 1.96-1.85 (m, 1H), 1.65-1.53 (m, 5H), 1.02 (t, *J*= 7.4 Hz, 3H); m/z (ES+): 405.0 $[M+H^+]^+$.

**Methyl (*R*)-2-((*S*)-4-(3-chlorophenyl)-2,3,9-trimethyl-6H-thieno[3,2-f][1,2,4]triazolo[4,3-a][1,4] diazepin-6-yl)butanoate (Compound 6)**

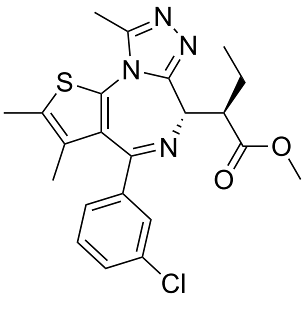

To a stirred solution of methyl (*R*)-2-((*S*)-5-(3-chlorophenyl)-6,7-dimethyl-2-oxo-2,3-dihydro-1H-thieno[2,3-e][1,4]diazepin-3-yl)butanoate (5.00 g, 12.35 mmol) in THF (60 mL) at -78 °C was dropwise added potassium *tert*-butoxide (24.70 mL, 24.70 mmol, 1M solution in THF), and stirring was maintained at this temperature for 30 minutes. Diethyl chlorophosphate (3.55 mL, 24.70 mmol) was added dropwise, and upon completion of the addition, the reaction mixture was allowed to warm to ambient temperature over 90 minutes, with an additional portion of THF (8 mL) being added during this time. Acetohydrazide (2.74 g, 37.05 mmol) was added portion-wise, and stirring was maintained at ambient temperature for 1 hour. *n*-Butanol (90 mL) was added and the reaction mixture was heated at 90 °C overnight, before being re-cooled to ambient temperature and concentrated under reduced pressure. The residue was partitioned between saturated aqueous sodium hydrogen carbonate solution and dichloromethane. The organic phase was separated, and the aqueous component was extracted with dichloromethane. The combined organic extracts were washed with brine, dried over anhydrous magnesium sulfate and concentrated under reduced pressure. Purification by flash column chromatography, eluting with 5-75% ethyl acetate/heptane, followed by re-concentration from cyclohexane/*tert*-butyl methyl ether mixture and drying under high vacuum at 60 °C afforded the title compound as a beige solid (2.80 g, 51%). $^1$H NMR (400 MHz, $CDCl_3$): δ 7.40-7.36 (m, 2H), 7.30-7.23 (m, 2H), 4.25 (d, *J*= 11.0 Hz, 1H), 3.99 (td, *J*= 10.7, 3.7 Hz, 1H), 3.87 (s, 3H), 2.68 (s, 3H), 2.42 (d, *J*= 0.6 Hz, 3H), 2.23-2.11 (m, 1H), 1.74-1.61 (m, 4H), 1.02 (t, *J*= 7.4 Hz, 3H); m/z (ES+): 443.2 $[M+H^+]^+$.

**Methyl (*R*)-2-((*S*)-2,3,9-trimethyl-4-(3-vinylphenyl)-6*H*-thieno[3,2-*f*][1,2,4]triazolo[4,3-*a*][1,4] diazepin-6-yl)butanoate (MR108)**

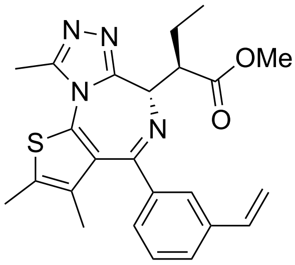

A solution of compound **6** (97 mg, 0.72 mmol) , XPhos Pd G2 (57 mg, 0.07 mmol) and *N,N*-diisopropylethylamine (0.17 mL, 0.96 mmol) in DMF (2 mL) and water (0.5 mL) was heated at 100 °C for 24 hours. Upon cooling to ambient temperature, the mixture was filtered through diatomaceous earth, the filtrate was concentrated under reduced pressure and the residue was and redissolved in dichloromethane. The solution was washed with water (2 x 50 mL) and brine (20 x 50 mL), dried over anhydrous magnesium sulfate and concentrated under reduced pressure. Purification by flash column chromatography, eluting with 0-12% methanol/dichloromethane, afforded the title compound as a yellow solid (90 mg, 91 %). $^1$H NMR (500 MHz, $CDCl_3$) δ 7.45 (d, *J* = 7.8 Hz, 1H), 7.42 (s, 1H), 7.28 (t, *J* = 7.6 Hz, 1H), 7.20 (d, *J* = 7.6 Hz, 1H), 6.68 (dd, *J* = 17.6, 10.9 Hz, 1H), 5.72 (dd, *J* = 17.7, 0.8 Hz, 1H), 5.26 (d, *J* = 10.8 Hz, 1H), 4.24 (d, *J* = 11.0 Hz, 1H), 4.01 (td, *J* = 10.8, 3.6 Hz, 1H), 3.87 (s, 3H), 2.67 (s, 3H), 2.40 (s, 3H), 2.22 – 2.14 (m, 1H), 1.71 – 1.61 (m, 4H), 1.03 (t, *J* = 7.4 Hz, 3H); $^{13}$C NMR (126 MHz, $CDCl_3$) δ 175.56, 164.20, 154.49, 149.71, 138.41, 137.83, 136.30, 132.00, 131.20, 130.76, 130.49, 128.54, 128.11, 128.00, 126.34, 114.69, 59.44, 51.53, 49.73, 23.25, 14.43, 13.14, 11.90, 11.69; m/z (ES+): 435.3 $[M+H^+]^+$; HRMS (ES+) calculated for $[(C23H26N4O2S)+H^+]^+$ 435.18492 found 435.18543

**Methyl (*R*)-2-((*S*)-4-(3-(3-aminoprop-1-yn-1-yl)phenyl)-2,3,9-trimethyl-6*H*-thieno[3,2-*f*][1,2,4]triazolo[4,3-*a*][1,4]diazepin-6-yl)butanoate (MR109)**

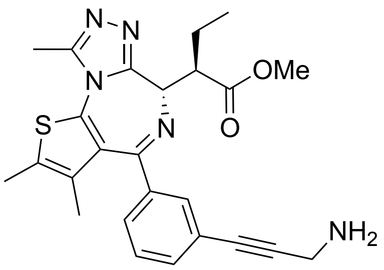

A stirred suspension of compound **6** (50 mg, 0.11 mmol), XPhos (16 mg, 0.03 mmol) , cesium carbonate (184 mg, 0.57 mmol), XPhos Pd G2 (27 mg, 0.03 mmol) and propargylamine (0.07 mL, 1.13 mmol) in THF (1.5mL) was heated at 100 °Cfor 1 hour, 90 °C for 2 hours, and then cooled to 60 °C and stirred overnight. Upon cooling to ambient temperature, the mixture was suspended in dichloromethane and filtered through diatomaceous earth. The filtrate was washed with water, dried over anhydrous magnesium sulfate and concentrated under reduced pressure. Purification by preparative reversed phase column chromatography, eluting with 5/95 to 95/5 MeCN/$H_2O$ (0.1 % formic acid), afforded the title compound as a white solid (7 mg, 13%). $^1$H NMR (500 MHz, $CDCl_3$) δ 7.36 (d, $J$ = 8.2 Hz, 2H), 7.31 (d, $J$ = 8.0 Hz, 2H), 4.23 (d, $J$ = 10.9 Hz, 1H), 3.98 (td, $J$ = 10.7, 3.6 Hz, 1H), 3.84 (s, 3H), 3.65 (s, 2H), 2.65 (s, 3H), 2.40 (s, 3H), 2.17 – 2.05 (m, 1H), 1.62 – 1.55 (m, 4H), 1.24 (s, 1H), 1.01 (t, $J$ = 7.4 Hz, 3H); m/z (ES+): 462.1 $[M+H^+]^+$. HRMS (ES+) calculated for $[(C25H27N5O2S)+H^+]^+$ 462.19582 found 462.19629

**Methyl (*R*)-2-((*S*)-4-(3-formylphenyl)-2,3,9-trimethyl-6*H*-thieno[3,2-*f*][1,2,4]triazolo[4,3-*a*][1,4] diazepin-6-yl)butanoate (MR115)**

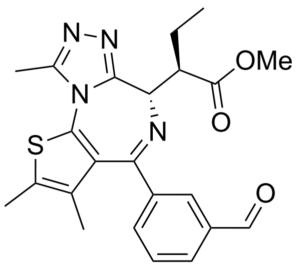

To a stirred solution of methyl **MR108** (29 mg, 0.07 mmol) and sodium periodate (42 mg, 0.19 mmol) in acetone (0.5 mL) and water (0.1 mL) was added osmium tetroxide (21 µL, 4%w/v solution in water). After stirring for 2 hours, the reaction mixture was diluted with EtOAc. The solution was washed with water, and the organics were dried over anhydrous magnesium sulfate and concentrated under reduced pressure. Purification by preparative reversed phase column chromatography, eluting with 5/95 to 95/5 MeCN/$H_2O$ (0.1 % $NH_4OH$) afforded the title compound as a white solid (28 mg, quant). $^1$H NMR (500 MHz, $CDCl_3$) δ 10.00 (d, $J$ = 1.8 Hz, 1H), 7.92 (d, $J$ = 7.6 Hz, 1H), 7.84 (s, 1H), 7.72 (d, $J$ = 7.8 Hz, 1H), 7.54 (t, $J$ = 7.7 Hz, 1H), 4.28 (d, $J$ = 11.1 Hz, 1H), 4.01 (td, $J$ = 10.7, 3.5 Hz, 1H), 3.89 (d, $J$ = 1.6 Hz, 3H), 2.69 (s, 3H), 2.42 (s, 3H), 2.19 – 2.16 (m, 1H), 1.69 (dt, $J$ = 9.9, 6.4 Hz, 1H), 1.65 (s, 3H), 1.03 (t, $J$ = 7.3 Hz, 3H); $^{13}$C NMR (126 MHz, $CDCl_3$) δ 191.7, 175.5, 163.1, 139.1, 136.4, 134.2, 132.0, 131.1, 130.6, 130.1, 129.3, 129.2, 77.2, 59.5, 51.7, 49.7, 23.3, 14.6, 13.2, 11.9, 11.6; m/z (ES+): 437.2 $[M+H^+]^+$. HRMS (ES+) calculated for $[(C23H24N4O3S)+H^+]^+$ 437.16419 found 437.16505

**methyl (2*R*)-2-((6*S*)-2,3,9-trimethyl-4-(3-(oxiran-2-yl)phenyl)-6*H*-thieno[3,2-*f*][1,2,4]triazolo[4,3-*a*][1,4]diazepin-6-yl)butanoate (MR111)**

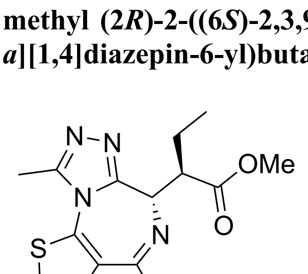

To a stirred solution of **MR108** (30 mg, 0.07 mmol) in dichloromethane (1 mL) at ambient temperature was added *m*-chloroperoxybenzoic acid (1 eq., 12 mg, 0.07 mmol) and the reaction stirred overnight. The mixture was diluted with dichloromethane (50 mL), washed with water (2 x 50 mL) and brine (2 x

50 mL), and the organic phase was dried over anhydrous magnesium sulfate and concentrated under reduced pressure. Purification by preparative reversed phase column chromatography, eluting with 5/95 to 95/5 MeCN/$H_2O$ (0.1% $NH_4OH$), afforded the title compound as a white solid (3 mg, 7%). $^1H$ NMR (500 MHz, $CDCl_3$) δ 7.27 – 7.18 (m, 4H), 4.20 – 4.14 (m, 1H), 3.98 – 3.89 (m, 1H), 3.82 – 3.75 (m, 4H), 3.09 – 3.02 (m, 1H), 2.72 – 2.66 (m, 1H), 2.65 – 2.58 (m, 3H), 2.34 (s, 3H), 2.15 – 2.06 (m, 1H), 1.65 – 1.52 (m, 4H), 0.95 (t, *J* = 7.4 Hz, 3H) ; m/z (ES+): 451.2 $[M+H^+]^+$. HRMS (ES+) calculated for $[(C24H26N4O3S)+H^+]^+$ 451.17984 found 451.18083

**Methyl (*R*)-2-((*S*)-4-(3-aminophenyl)-2,3,9-trimethyl-6H-thieno[3,2-f][1,2,4]triazolo[4,3-a][1,4]diazepin-6-yl)butanoate (Compound 8)**

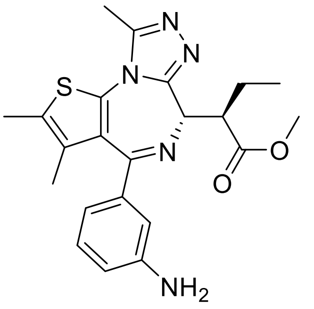

To a stirred solution of compound **6** (1.60 g, 3.61 mmol) in 1,4-dioxane (50 mL) was added benzophenone imine (0.79 g, 4.33 mmol) and potassium phosphate tribasic (1.92 g, 9.03 mmol), and the reaction mixture was degassed and back-filled with argon (g). *t*-BuXPhos (0.23 g, 0.54 mmol) and $Pd_2(dba)_3$ (0.17 g, 0.18 mmol) were added, and after further degassing with back-filling of argon (g), the reaction mixture was heated at 85 °C overnight. The reaction mixture was cooled to 55 °C and partially concentrated under reduced pressure, the subsequent residue being partitioned between water and ethyl acetate. The organic phase was separated, and the aqueous component was extracted with ethyl acetate. The combined organic extracts were washed with brine, dried over anhydrous magnesium sulfate and concentrated under reduced pressure to a brown oil. The crude imine was dissolved in THF (30 mL) and treated with 1M hydrochloric acid (30 mL), and the resulting solution was stirred at ambient temperature for 2 hours before being partially concentrated under reduced pressure. The aqueous residue was further diluted with water and washed with ethyl acetate. The aqueous phase was then adjusted to pH8-9 by slow addition of saturated aqueous sodium hydrogen carbonate solution and extracted with ethyl acetate (3 portions). These organic extracts were combined, dried over anhydrous magnesium sulfate and concentrated under reduced pressure. Purification by flash column chromatography, eluting with 0-2.5% 7M methanolic ammonia/dichloromethane, afforded a yellow foam. This process was repeated, followed by trituration with *tert*-butyl methyl ether/cyclohexane, affording the title compound as a yellow solid (0.76 g, 50%). $^1H$ NMR (400 MHz, $CDCl_3$): δ 7.10 (t, *J*= 7.8 Hz, 1H), 6.75-6.69 (m, 2H), 6.66 (d, *J*= 7.6 Hz, 1H), 4.22 (d, *J*= 11.0 Hz, 1H), 3.99 (td, *J*= 10.7, 3.7 Hz, 1H), 3.84 (s, 3H), 2.66 (s, 3H), 2.40 (s, 3H), 2.24-2.13 (m, 1H), 1.73-1.61 (m, 4H), 1.01 (t, *J*= 7.4 Hz, 3H); m/z (ES+): 446.2 $[M+Na^+]^+$.

**methyl (*R*)-2-((*S*)-4-(3-(2-chloroacetamido)phenyl)-2,3,9-trimethyl-6*H*-thieno[3,2-*f*][1,2,4]triazolo[4,3-*a*][1,4]diazepin-6-yl)butanoate (MR118)**

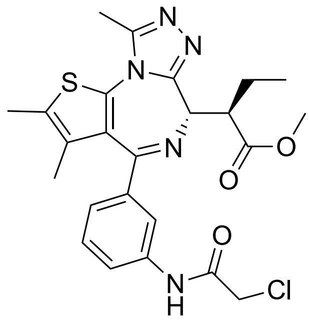

To a stirred solution of compound **8** (60 mg, 0.14 mmol) and *N*,*N*-diisopropylethylamine (250 µL, 0.14 mmol) in dichloromethane (1.4 mL) at ambient temperature was dropwise added 2-chloroacetyl chloride (430 µL, 0.53 mmol). The reaction mixture was stirred for 2 hours before being quenched with a few drops of methanol and then concentrated under reduced pressure. The residue was partitioned between dichloromethane and water, and the organic phase was separated. The organics were washed with water (2 x 50 mL) and brine (2 x 50 mL), dried over anhydrous magnesium sulfate, and concentrated under reduced pressure, affording the title compound as a white solid (50 mg, 71%). $^1$H NMR (500 MHz, $CDCl_3$) δ 8.46 (s, 1H), 7.66 (d, *J* = 9.5 Hz, 1H), 7.58 (s, 1H), 7.30 (t, *J* = 7.9 Hz, 1H), 7.11 (d, *J* = 7.7 Hz, 1H), 4.24 (d, *J* = 11.0 Hz, 1H), 4.17 (s, 2H), 3.98 (td, *J* = 10.7, 3.7 Hz, 1H), 3.84 (s, 3H), 2.66 (s, 3H), 2.41 (s, 3H), 2.16-2.10 (m, 1H), 1.73 – 1.59 (m, 4H), 1.00 (t, *J* = 7.4 Hz, 3H) ; $^{13}$C NMR (126 MHz, $CDCl_3$) δ 175.47, 164.05, 163.91, 154.50, 149.94, 139.12, 137.22, 132.13, 131.23, 130.84, 130.63, 129.28, 125.46, 122.33, 120.02, 59.48, 51.78, 49.80, 43.03, 23.34, 14.56, 13.28, 11.98, 11.77. m/z (ES+): 500.4 $[M+H^+]^+$; HRMS (ES+) calculated for $[(C24H26ClN5O3S)+H^+]^+$ 500.15176 found 500.15287

**methyl (*R*)-2-((*S*)-2,3,9-trimethyl-4-(3-(vinylsulfonamido)phenyl)-6*H*-thieno[3,2-*f*][1,2,4]triazolo[4,3-*a*][1,4]diazepin-6-yl)butanoate (MR117)**

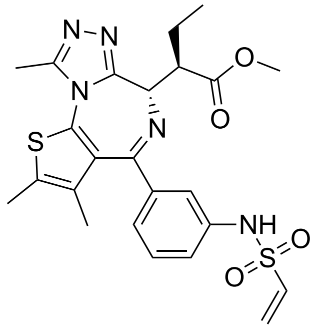

To a stirred solution of compound **8** (60 mg, 0.14 mmol) and pyridine (230 µL, 0.28 mmol) in dichloromethane (1.4 mL) at 0 °C was dropwise added vinyl sulfonyl chloride (120 µL, 0.12 mmol). The reaction mixture was stirred for 2 hours before being quenched with a few drops of methanol and then concentrated under reduced pressure. The residue was partitioned between dichloromethane and water, and the organic phase was separated. The organics were washed with water (2 x 50 mL) and brine (2 x 50 mL), dried over anhydrous magnesium sulfate, and concentrated under reduced pressure, affording the title compound as a white crystalline solid (43 mg, 54%). $^1$H NMR (500 MHz, $CDCl_3$) δ 7.43 (s, 1H), 7.31 – 7.20 (m, 2H), 7.08 (d, *J* = 7.1 Hz, 1H), 6.52 (dd, *J* = 16.5, 9.9 Hz, 1H), 6.25 (d, *J* = 16.6 Hz, 1H), 5.93 (d, *J* = 9.9 Hz, 1H), 5.29 (s, 1H), 4.25 (d, *J* = 11.0 Hz, 1H), 3.96 (td, *J* = 10.7, 3.7 Hz, 1H), 3.85 (s, 3H), 2.66 (s, 3H), 2.40 (s, 3H), 2.23 – 2.01 (m, 1H), 1.73 – 1.50 (m, 4H), 1.01 (t, *J* = 7.4 Hz, 3H); $^{13}$C NMR (126 MHz, $CDCl_3$) δ 175.27, 163.57, 154.36, 149.86, 139.35, 136.90, 134.95, 131.96, 131.02, 130.83, 130.51, 129.43, 128.85, 125.15, 122.32, 120.08, 59.36, 51.80, 49.78, 23.23,

14.37, 13.13, 11.83, 11.66; m/z (ES+): 514.0 $[M+H^+]^+$; HRMS (ES+) calculated for $[(C24H27N5O4S2)+H^+]^+$ 514.15772 176 found 514.15898

**methyl (*R*)-2-((*S*)-4-(3-acrylamidophenyl)-2,3,9-trimethyl-6*H*-thieno[3,2-*f*][1,2,4]triazolo[4,3-*a*][1,4]diazepin-6-yl)butanoate (MR119)**

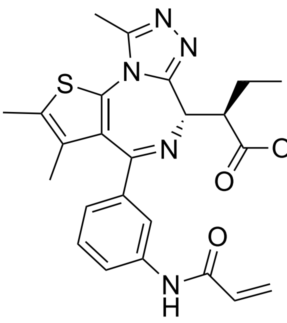

To a stirred solution of compound **8** (60 mg, 0.14 mmol) and triethylamine (240 µL, 0.20 mmol) in dichloromethane (1.4 mL) at ambient temperature was dropwise added acryloyl chloride (100 µL, 0.13 mmol). The reaction mixture was stirred for 2 hours before being quenched with a few drops of methanol and then concentrated under reduced pressure. The residue was partitioned between dichloromethane and water, and the organic phase was separated. The organics were washed with water (2 x 50 mL) and brine (2 x 50 mL), dried over anhydrous magnesium sulfate, and concentrated under reduced pressure. Purification by flash column chromatography, eluting with 0-5% methanol/dichloromethane, afforded the title compound as a yellow solid (40 mg, 59%). $^1$H NMR (500 MHz, $CDCl_3$) δ 7.70 – 7.63 (m, 2H), 7.51 (s, 1H), 7.22 (t, *J* = 7.9 Hz, 1H), 7.02 (d, *J* = 7.7 Hz, 1H), 6.34 (dd, *J* = 16.8, 1.3 Hz, 1H), 6.19 (dd, *J* = 16.9, 10.2 Hz, 1H), 5.68 (dd, *J* = 10.2, 1.3 Hz, 1H), 4.18 (d, *J* = 11.0 Hz, 1H), 3.91 (td, *J* = 10.7, 3.7 Hz, 1H), 3.77 (s, 3H), 2.59 (s, 3H), 2.34 (s, 3H), 2.17 – 2.05 (m, 1H), 1.64 – 1.56 (m, 4H), 0.94 (t, *J* = 7.4 Hz, 3H); m/z (ES+): 478.1 $[M+H^+]^+$. HRMS (ES+) calculated for $[(C25H27N5O3S)+H^+]^+$ 478.19074 found 478.19158

**methyl (*R*)-2-((*S*)-4-(4-((*tert*-butoxycarbonyl)amino)phenyl)-2,3,9-trimethyl-6*H*-thieno[3,2-*f*][1,2,4]triazolo[4,3-*a*][1,4]diazepin-6-yl)butanoate (MR126)**

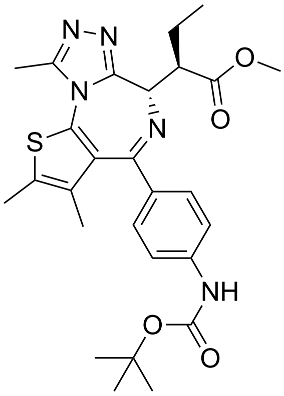

To a stirred solution of compound **7** (100 mg, 1 eq., 0.24 mmol) in methanol at ambient temperature (1.2mL) was added Boc anhydride (1.5 eq, 27 mg, 0.36 mmol) and TEA (2 eq, 66 µL, 0.47 mmol) and the reaction was stirred at 50 °C overnight. The reaction mixture was concentrated under reduced pressure and the residue was taken up in dichloromethane. The solution was washed with saturated aqueous sodium hydrogen carbonate solution (2x50 mL) and brine (2x50 mL), dried over anhydrous magnesium sulfate, and concentrated under reduced pressure. Purification by flash column chromatography, eluting with 0-5% methanol/dichloromethane, afforded the title compound as an off-white solid (90 mg, 73 %). $^1$H NMR (500 MHz, $CDCl_3$) 7.37 (d, *J* = 8.6 Hz, 2H), 7.32 (d, *J* = 8.7 Hz,

2H), 6.81 (s, 1H), 4.22 (d, *J* = 10.9 Hz, 1H), 4.0 (td, *J* = 10.8, 3.7 Hz, 1H), 3.85 (s, 3H), 2.68 (s, 3H), 2.41 (s, 3H), 2.24 – 2.13 (m, 1H), 1.87 – 1.81 (m, 1H), 1.70 (s, 3H), 1.52 (s, 9H), 1.03 (t, *J* = 7.4 Hz, 3H); $^{13}$C NMR (126 MHz, $CDCl_3$) δ 175.51, 163.45, 154.72, 152.31, 149.69, 140.63, 132.59, 131.76, 131.29, 130.97, 130.43, 129.49, 117.66, 80.97, 77.29, 77.23, 77.03, 76.78, 59.29, 51.52, 49.79, 28.31, 23.24, 14.48, 13.12, 11.86, 11.70; m/z (ES+): 524.5 $[M+H^+]^+$.

**methyl (*R*)-2-((*S*)-4-(4-((*tert*-Butoxycarbonyl)amino)phenyl)-2,3,9-trimethyl-6H-thieno[3,2-f][1,2,4]triazolo[4,3-a][1,4]diazepin-6-yl)butanoic acid (MR137)**

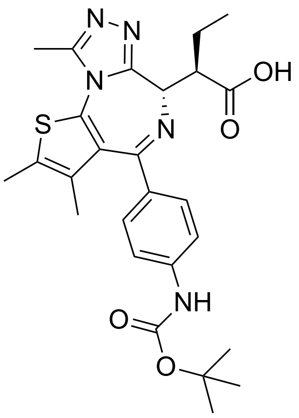


To a stirred solution of **MR126** (2.15 g (uncorr.), 1.51 g (corr.), 2.87 mmol (corr.)) in THF (35 mL) and methanol (8.5 mL) was added lithium hydroxide (21.00 mL, 0.65M aqueous solution), and the resulting orange solution was heated at 45 °C for 16 hours. The volatiles were removed under reduced pressure, and the remaining aqueous was diluted with water (40 mL) and adjusted to pH 1 by addition of 2M hydrochloric acid (20 mL). The solution was extracted with dichloromethane (3 x 100 mL), and the combined organic extracts were dried over anhydrous magnesium sulfate and concentrated under reduced pressure. Purification by flash column chromatography, eluting with 0-10% methanol in dichloromethane, followed by a trituration with dichloromethane/*tert*-butyl methyl ether /heptane (1:2:2; 50 mL), afforded the title compound as a white solid (1.09 g (uncorr.), 0.87 g (corr.), 59% (corr.)). $^{1}$H NMR (400 MHz, DMSO-d6) δ 12.41 (br. s, 1H), 9.58 (s, 1H), 7.48 (d, *J* = 8.9 Hz, 2H), 7.28 (d, *J* = 8.6 Hz, 2H), 4.06 (d, *J* = 10.7 Hz, 1H), 3.53 (td, *J* = 10.6, 3.5 Hz, 1H), 2.58 (s, 3H), 2.41 (d, *J* = 0.4 Hz, 3H), 2.01 – 1.90 (m, 1H), 1.63 (d, *J* = 0.4 Hz, 2H), 1.60 – 1.49 (m, 1H), 1.47 (s, 9H), 0.95 (t, *J* = 7.4 Hz, 3H); LCMS purity = 92.6%; m/z (ES+): 510.30 $[M+H^+]^+$.

**_tert_-butyl (4-((6_S_)-6-((3_S_,18_R_)-3-((2_S_,4_R_)-4-hydroxy-2-((4-(4-methylthiazol-5-yl)benzyl)carbamoyl)pyrrolidine-1-carbonyl)-2,2-dimethyl-5,17-dioxo-7,10,13-trioxa-4,16-diazaicosan-18-yl)-2,3,9-trimethyl-6_H_-thieno[3,2-_f_][1,2,4]triazolo[4,3-_a_][1,4]diazepin-4-yl)phenyl)carbamate (MR162)**

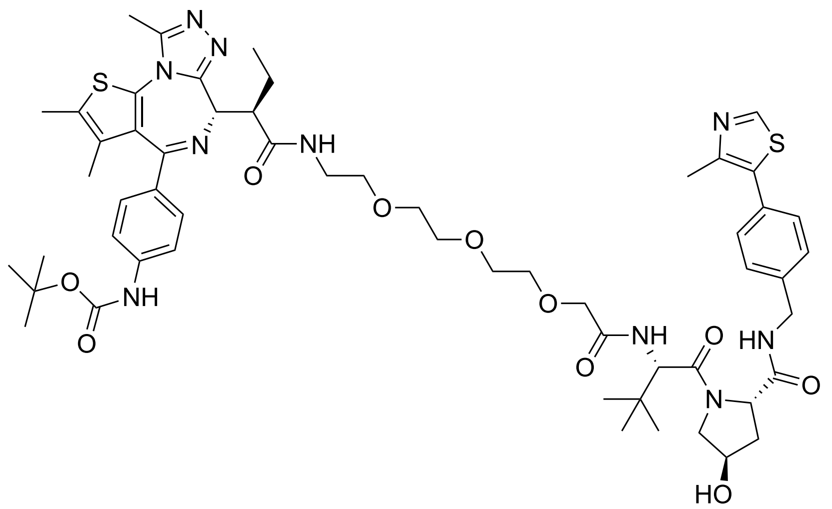

**MR137** (10 mg, 0.020 mmol) was dissolved in anhydrous dichloromethane and _N_,_N_-diisopropylethylamine (0.014 mL, 0.08 mmol) was added, followed by HATU (7.5 mg, 0.020 mmol) and the resulting mixture was stirred for 5 min before adding 4-((2-(2-(2-aminoethoxy)ethoxy)ethyl)amino)-2-(2,6-dioxopiperidin-3-yl)isoindoline-1,3-dione (9.5mg, 0.014 mmol). After stirring for 4 hours at ambient temperature, the reaction mixture was concentrated under reduced pressure. Purification by flash column chromatography, eluting with 0-10% methanol/dichloromethane, afforded the title compound as a white solid (15 mg, 76%). $^1$H NMR (500 MHz, $CDCl_3$) δ 8.63 (s, 1H), 8.51 (d, $J$ = 4.4 Hz, 1H), 8.20 (d, $J$ = 8.3 Hz, 1H), 8.02 (s, 1H), 7.28 (s, 1H), 7.23 (t, $J$ = 8.8 Hz, 4H), 7.17 (d, $J$ = 4.1 Hz, 4H), 6.66 (s, 1H), 4.77 (t, $J$ = 8.2 Hz, 1H), 4.69 (d, $J$ = 9.5 Hz, 1H), 4.43 (s, 1H), 4.35 (dd, $J$ = 15.6, 7.0 Hz, 1H), 4.17 (d, $J$ = 10.2 Hz, 1H), 4.09 (d, $J$ = 10.9 Hz, 1H), 4.00 (s, 1H), 3.76 – 3.51 (m, 14H), 3.42 (s, 2H), 2.56 (s, 3H), 2.45 (s, 3H), 2.31 (s, 3H), 2.21 – 2.07 (m, 3H), 1.89 – 1.80 (m, 1H), 1.54 (s, 3H), 1.43 (s, 8H), 1.37 – 1.31 (m, 1H), 0.98 – 0.86 (m, 12H). m/z (ES+): 556.9 $[M+2H^+]^{2+}$.

**(2*S*,4*R*)-1-((2*S*,17*R*)-17-((6*S*)-4-(4-acrylamidophenyl)-2,3,9-trimethyl-6*H*-thieno[3,2-*f*][1,2,4]triazolo[4,3-*a*][1,4]diazepin-6-yl)-2-(*tert*-butyl)-4,16-dioxo-6,9,12-trioxa-3,15-diazanonadecanoyl)-4-hydroxy-*N*-(4-(4-methylthiazol-5-yl)benzyl)pyrrolidine-2-carboxamide (MR170)**

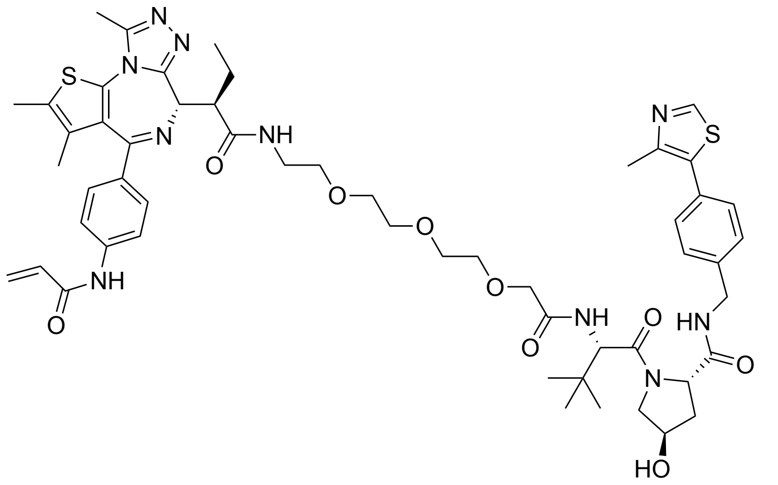

**MR162** (15 mg, 0.0135 mmol) was treated with a solution of 20% TFA in dichloromethane and then concentrated under reduced pressure. The residue was dissolved in dichloromethane (1 mL) and to this solution, *N*,*N*-diisopropylethylamine (0.004 mL, 0.022 mmol) was added followed by dropwise addition of a solution of acryloyl chloride (0.001 mL, 0.015 mmol) in dichloromethane (0.2 mL). After stirring for 30 minutes, the reaction was quenched by addition of a small amount of MeOH and then concentrated under reduced pressure. Purification by preparative reversed phase chromatography, using a gradient of 5/95 to 95/5 MeCN/$H_2O$ (0.1% formic acid), afforded the title compound as a white solid (5 mg, 34% over two steps). $^1$H NMR (500 MHz, $CDCl_3$) δ 8.60 (s, 1H), 7.90 (s, 1H), 7.77 (t, *J* = 5.5 Hz, 1H), 7.44 – 7.39 (m, 2H), 7.31 – 7.22 (m, 5H), 7.21 – 7.18 (m, 2H), 7.18 – 7.14 (m, 2H), 6.35 (dd, *J* = 16.8, 1.3 Hz, 1H), 6.17 (dd, *J* = 16.9, 10.2 Hz, 1H), 5.69 (dd, *J* = 10.2, 1.3 Hz, 1H), 4.74 – 4.64 (m, 2H), 4.43 – 4.30 (m, 1H), 4.15 (d, *J* = 9.9 Hz, 1H), 4.06 – 3.97 (m, 3H), 3.83 m, 1H), 2.11 – 2.01 (m, 1H), 1.90 – 1.81 (m, 1H), 1.62 – 1.52 (m, 4H), 0.97 – 0.88 (m, 12H). m/z (ES+): 1066.3 $[M+H^+]^+$.

***tert*-butyl (4-((6*S*)-6-((*R*)-4,21-dioxo-25-((3a*S*,4*S*,6a*R*)-2-oxohexahydro-1*H*-thieno[3,4-*d*]imidazol-4-yl)-8,11,14,17-tetraoxa-5,20-diazapentacosan-3-yl)-2,3,9-trimethyl-6*H*-thieno[3,2-*f*][1,2,4]triazolo[4,3-*a*][1,4]diazepin-4-yl)phenyl)carbamate (MR129)**

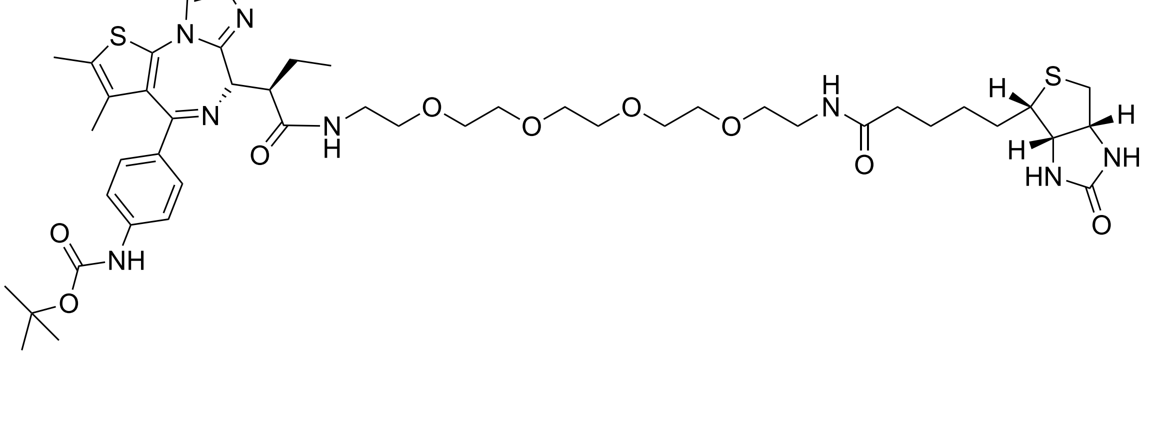

To a stirred solution of **MR137** (20 mg, 0.04 mmol) and *N,N*-diisopropylethylamine (205 µL, 0.12 mmol) in DMF (1 mL) at ambient temperature was added HATU (18 mg, 0.05 mmol) and the resulting mixture was stirred for 5 min before adding *N*-(14-amino-3,6,9,12-tetraoxatetradecyl)-5-((3a*S*,4*S*,6a*R*)-2-oxohexahydro-1*H*-thieno[3,4-*d*]imidazol-4-yl)pentanamide (36 mg, 0.08 mmol). After stirring for 16 hours, the reaction mixture was concentrated under reduced pressure. Purification of the residue by flash column chromatography, eluting with 0-10% methanol/dichloromethane, afforded the title compound as a white solid (30 mg, 80%). $^1$H NMR (500 MHz, $CDCl_3$) δ 7.51 (t, *J* = 5.4 Hz, 1H), 7.43 (s, 1H), 7.33 (d, *J* = 8.3 Hz, 2H), 7.25 (d, *J* = 8.4 Hz, 2H), 7.06 (t, *J* = 5.6 Hz, 1H), 6.14 (s, 1H), 5.51 (s, 1H), 4.29 (dd, *J* = 8.0, 4.9 Hz, 1H), 4.15 (d, *J* = 10.2 Hz, 1H), 4.10 (dd, *J* = 8.3, 4.6 Hz, 1H), 3.67 – 3.59 (m, 1H), 3.62 – 3.50 (m, 15H), 3.53 – 3.44 (m, 3H), 3.40 (s, 4H), 3.38 – 3.30 (m, 2H), 3.04 (q, *J* = 7.4 Hz, 0H), 2.96 (td, *J* = 7.3, 4.5 Hz, 1H), 2.74 – 2.68 (m, 1H), 2.59 (s, 3H), 2.57 – 2.51 (m, 1H), 2.32 (s, 3H), 2.10 (d, *J* = 7.3 Hz, 1H), 1.94 (ddd, *J* = 13.1, 6.5, 3.0 Hz, 1H), 1.67 – 1.60 (m, 1H), 1.59 (s, 3H), 1.55 (dt, *J* = 14.1, 6.5 Hz, 3H), 1.53 – 1.42 (m, 1H), 1.43 (s, 8H), 1.37 (s, 3H), 1.28 (p, *J* = 7.7 Hz, 2H), 0.95 (t, *J* = 7.4 Hz, 3H); m/z (ES+): 477.9 $[M+2H^+]^{2+}$.

**_N_-((17*R*)-17-((6*S*)-4-(4-acrylamidophenyl)-2,3,9-trimethyl-6*H*-thieno[3,2-*f*][1,2,4]triazolo[4,3-*a*][1,4]diazepin-6-yl)-16-oxo-3,6,9,12-tetraoxa-15-azanonadecyl)-5-((3a*S*,4*S*,6a*R*)-2-oxohexahydro-1*H*-thieno[3,4-*d*]imidazol-4-yl)pentanamide (MR169)**

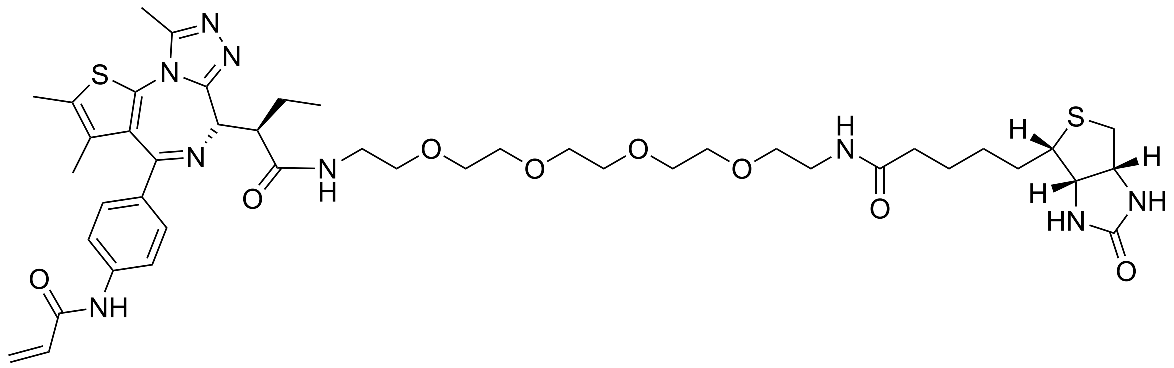

A solution of **MR129** (10 mg, 0.01 mmol) in 20% TFA/dichloromethane (1 mL) was stirred for 30 minutes and then concentrated under reduce pressure. The residue was re-dissolved in dichloromethane (1 mL) and the solution was treated with *N,N*-diisopropylethylamine (10 µL, 0.05 mmol) followed by acryloyl chloride (20 µL, 0.02 mmol). After stirring for 2 hours, the reaction mixture was quenched by addition of a few drops of methanol and then concentrated under reduced pressure. Purification by preparative HPLC afforded the title compound as a white solid (3 mg, 28%).$^1$H NMR (500 MHz, $CDCl_3$) δ 9.27 (s, 1H), 8.03 – 7.99 (m, 1H), 7.66-7.60 (m, 3H), 7.31 – 7.26 (m, 2H), 6.88 (s, 1H), 6.38 – 6.33 (m, 2H), 5.73 (s, 1H), 5.68-5.63 (m, 1H), 4.80 (s, 1H), 4.24 (s, 1H), 4.15 (d, *J* = 10.2 Hz, 1H), 3.87 (s, 1H), 3.68 – 3.49 (m, 14H), 3.57 (s, 5H), 3.52 – 3.43 (m, 2H), 3.36 – 3.25 (m, 1H), 2.86 – 2.78 (m, 1H), 2.74 – 2.67 (m, 1H), 2.59 (s, 3H), 2.57 – 2.51 (m, 1H), 2.32 (s, 3H), 2.06 – 2.00 (m, 2H), 1.94 – 1.88 (m, 2H), 1.51 – 1.42 (m, 1H), 1.35 – 1.29 (m, 2H), 1.25 – 1.16 (m, 4H), 0.95 (t, *J* = 7.3 Hz, 3H); m/z (ES+) : 908.5 $[M+H^+]^+$; HRMS (ES+) calculated for $[(C44H62N9O8S2)+2H^+]^{+2}$ 454.20759 found 454.71312

***tert*-butyl (4-((6*S*)-2,3,9-trimethyl-6-((*R*)-14-oxo-4,7,10-trioxa-13-azaheptadec-1-yn-15-yl)-6*H*-thieno[3,2-*f*][1,2,4]triazolo[4,3-*a*][1,4]diazepin-4-yl)phenyl)carbamate (MR152)**

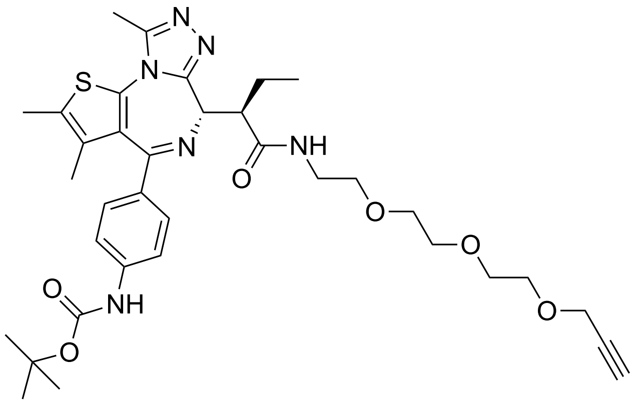

**MR137** (17 mg, 0.033 mmol) was dissolved in DMF and *N*,*N*-diisopropylethylamine (23 µL, 0.13 mmol) was added, followed by HATU (.7 mg, 0.0334 mmol) and the resulting mixture was stirred for 5 min before adding 2-(2-(2-(prop-2-yn-1-yloxy)ethoxy)ethoxy)ethan-1-amine (8.1 mg, 0.043 mmol). After stirring for 1 hour at ambient temperature, the reaction mixture was concentrated under reduced pressure. Purification by flash column chromatography, eluting with 0-10% methanol/dichloromethane, afforded the title compound as a yellow solid (10 mg, 44%), which was used directly in the subsequent stage. m/z (ES+): 679.1 $[M+H^+]^+$.

**(2*R*)-2-((6*S*)-4-(4-acrylamidophenyl)-2,3,9-trimethyl-6*H*-thieno[3,2-*f*][1,2,4]triazolo[4,3-*a*][1,4]diazepin-6-yl)-*N*-(2-(2-(2-(prop-2-yn-1-yloxy)ethoxy)ethoxy)ethyl)butanamide (MR155)**

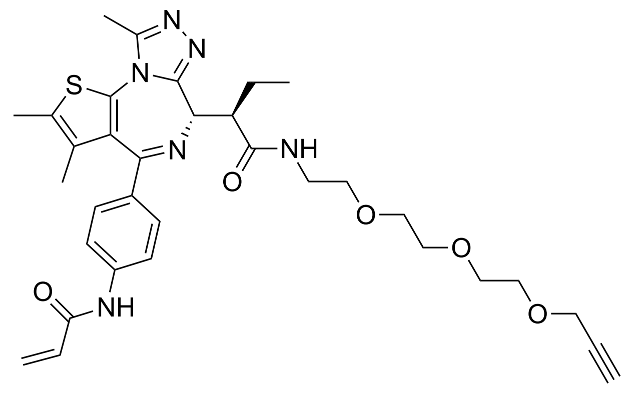

To a solution of **MR152** (10 mg, 0.017 mmol) was added a solution of 20% TFA in dichloromethane (1 mL) and stirred for 30 min. Once Boc group removal was confirmed by LC-MS, the reaction mixture was concentrated under reduced pressure. A mixture of acrylic acid (2.4 mg, 0.033 mmol), HATU (12.7 mg, 0.033 mmol) and *N*,*N*-diisopropylethylamine (15 µL, 0.083 mmol) in dichloromethane (0.5 mL) was stirred for 5 minutes before adding a solution of the crude aniline (0.5 equiv., 8 mg, 0.017 mmol) in dichloromethane (0.5 mL). After stirring for 1 hour, the reaction mixture was concentrated under reduced pressure. Purification by preparative reversed phase HPLC, using a gradient of 5/95 to 95/5 MeCN/$H_2O$ (0.1% formic acid), afforded the title compound as a white solid (3 mg, 15% over two steps). $^1H$ NMR (500 MHz, $CDCl_3$) δ 7.73 (s, 1H), 7.55 (d, *J* = 8.4 Hz, 2H), 7.36 – 7.31 (m, 2H), 6.72 (s, 1H), 6.36 (d, *J* = 16.8 Hz, 1H), 6.20 (dd, *J* = 16.8, 10.3 Hz, 1H), 5.69 (d, *J* = 10.1 Hz, 1H), 4.19 – 4.08 (m, 4H), 3.74 – 3.47 (m, 9H), 3.48 – 3.35 (m, 1H), 2.58 (s, 3H), 2.40 – 2.35 (m, 1H), 2.32 (s, 3H), 2.03 – 1.94 (m, 1H), 1.70 – 1.61 (m, 1H), 1.60 (s, 3H), 0.96 (t, *J* = 7.3 Hz, 3H); m/z (ES+): 633.4 $[M+H^+]^+$.

**5-((2,2-dimethyl-4-oxo-3,8,11,14-tetraoxa-5-azahexadecan-16-yl)carbamoyl)-2-(6 (dimethylamino)-3-(dimethyliminio)-3*H*-xanthen-9-yl)benzoate (MR185)**

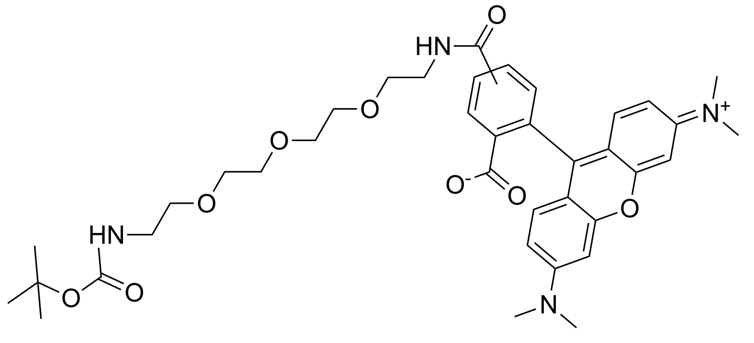

To a solution of 5(6)-TAMRA (23 mg, 0.05 mmol) in dichloromethane (1 mL), *N,N*-diisopropylethylamine (0.016 mL, 0.0924 mmol) was added followed by HATU (24.4 mg, 0.0254 mmol) and the reaction was stirred for 5 min before adding a mixture of *Boc*-amino-PEG3-amine (23.4 mg, 0.046 mmol) and *N,N*-diisopropylethylamine (1.5 eq, 0.37 mL, 0.21 mmol ). The mixture was stirred at room temperature for 1 h. The solvent was evaporated under reduced pressure and the product was purified by flash column chromatography, eluting with 0-5% methanol/dichloromethane, affording the title compound as a pink solid (20 mg,0.02 mmol, 61.9 % yield). m/z (ES+): 705.4 $[M+H^+]^+$.

***N*-((17*R*)-17-((6*S*)-4-(4-acrylamidophenyl)-2,3,9-trimethyl-6*H*-thieno[3,2-*f*][1,2,4]triazolo[4,3-*a*][1,4]diazepin-6-yl)-16-oxo-3,6,9,12-tetraoxa-15-azanonadecyl)-5-((3a*S*,4*S*,6a*R*)-2-oxohexahydro-1*H*-thieno[3,4-*d*]imidazol-4-yl)pentanamide (MR202)**

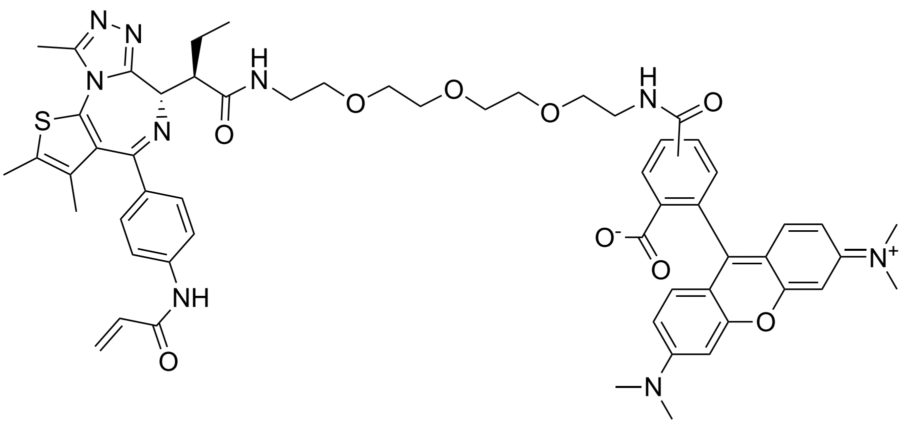

**MR185** (11 mg, 0.018 mmol) was dissolved in 20% TFA in DCM and the reaction stirred for one hour. After one hour, the solvent was evaporated and to the crude amine, a solution of **MR137** (10 mg, 0.18 mmol) and *N*,*N*-diisopropylethylamine (205 µL, 0.12 mmol) in DMF (1 mL) at ambient temperature was added, followed by HATU (12 mg, 0.018 mmol) and the resulting mixture . After stirring for 16 hours, the reaction mixture was concentrated under reduced pressure. The resulting Boc-aniline precursor (10 mg (uncorr.), 0.18 µmol (corr.)) was dissolved in a solution of 20% TFA in dichloromethane (2 mL) and the reaction mixture was stirred at room temperature for 1 hour. The solvent was evaporated, and the resulting crude resuspended in dichloromethane. The stirred solution was treated with triethylamine (0.90 mL) followed by the portion-wise addition of acryloyl chloride diluted in dichloromethane (440 µL of acryloyl chloride dissolved in 1 mL of dichloromethane, 0.6 mL added), and the reaction mixture was stirred at ambient temperature for 4 hours before being concentrated under reduced pressure. Purification by preparative reversed phase HPLC, using a gradient of 5/95 to 95/5 MeCN/$H_2O$ (0.1% formic acid), afforded the title compound as a white solid (10 mg, 53% over two steps).$^1$H NMR (500 MHz, $CDCl_3$) δ 9.20 (s, 1H), 8.10 (d, $J$ = 8.1 Hz, 1H), 8.02 (dd, $J$ =

8.1, 1.6 Hz, 1H), 7.70 (d, $J$ = 1.5 Hz, 1H), 7.65 (s, 1H), 7.60 (d, $J$ = 8.4 Hz, 2H), 7.28 (s, 1H), 7.01 (s, 1H), 6.88 – 6.83 (m, 1H), 6.79 (d, $J$ = 9.1 Hz, 1H), 6.52 – 6.46 (m, 2H), 6.44 (dd, $J$ = 9.2, 2.5 Hz, 1H), 6.40 (s, 1H), 6.30 – 6.17 (m, 2H), 5.49 (dd, $J$ = 9.0, 2.8 Hz, 1H), 4.18 (s, 1H), 3.68 – 3.62 (m, 3H), 3.61 (q, $J$ = 6.4 Hz, 4H), 3.56 (dt, $J$ = 10.3, 5.4 Hz, 5H), 3.53 (s, 4H), 3.02 (s, 6H), 2.96 (s, 7H), 2.64 (s, 3H), 2.39 (s, 3H), 2.38 (d, $J$ = 8.3 Hz, 1H), 1.93 (s, 1H), 1.64 (s, 3H), 1.62 (s, 1H), 0.98 (t, $J$ = 7.3 Hz, 3H). m/z (ES+): 526.1 $[M+2H^+]^{2+}$ HRMS (ES+) calculated for $[(C57H64N9O9S)+2H^+]^{+2}$ 525.73075 found 525.73139

**4-((2-(2-(2-(2-Aminoethoxy)ethoxy)ethoxy)ethyl)carbamoyl)-2-(3-(3-fluoroazetidin-1-ium-1-ylidene)-7-(3-fluoroazetidin-1-yl)-5,5-dimethyl-3,5-dihydrodibenzo[b,e]silin-10-yl)benzoate**

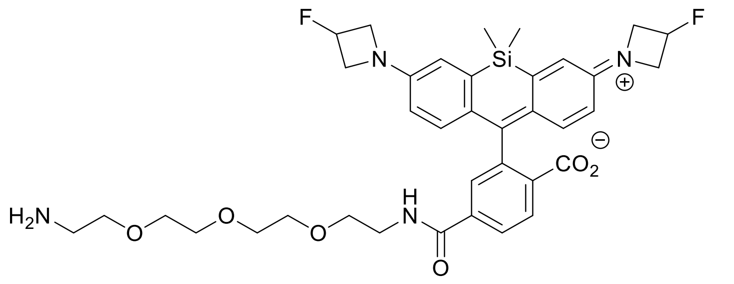

To a stirred solution of Amino-PEG3-amine (450 mg, 0.29 mmol) in dichloromethane (3 mL) at ambient temperature was added a solution of Janelia Fluor® 635, NHS ester (200 mg (uncorr.), 184 mg (corr.), 0.29 mmol (corr.)) in dichloromethane (5 mL), and the reaction mixture was stirred for 2 hours before being concentrated under reduced pressure. Purification by flash column chromatography, eluting with 0-20% methanol in dichloromethane, afforded the title compound as a pale-yellow solid (175 mg (uncorr.), 166 mg (corr.), 81% (corr.)). $^1$H NMR (400 MHz, DMSO-d6): δ 8.85 (t, $J$ = 5.5 Hz, 1H), 8.08 (dd, $J$ = 8.0, 1.3 Hz, 1H), 8.06 – 8.01 (m, 1H), 7.68 – 7.64 (m, 1H), 6.82 (d, $J$ = 2.6 Hz, 2H), 6.66 (d, $J$ = 8.7 Hz, 2H), 6.42 (dd, $J$ = 8.8, 2.7 Hz, 2H), 5.47 (dtt, $J$ = 57.3, 5.8, 3.0 Hz, 2H), 4.25 – 4.10 (m, 4H), 3.98 – 3.84 (m, 4H), 3.53 – 3.40 (m, 10H), 3.40 – 3.34 (m, 2H), 3.31 (t, $J$ = 5.7 Hz, 2H), 2.61 (t, $J$ = 5.7 Hz, 2H), 2.37 (br. s, $J$ = 17.3 Hz, 2H), 0.63 (s, 3H), 0.52 (s, 3H); LCMS purity = 98.7%; m/z (ES+): 707.40 $[M+H^+]^+$.

**4-(((*R*)-14-((*S*)-4-(4-Acrylamidophenyl)-2,3,9-trimethyl-6H-thieno[3,2-f][1,2,4]triazolo[4,3-a][1,4]diazepin-6-yl)-13-oxo-3,6,9-trioxa-12-azahexadecyl)carbamoyl)-2-(3-(3-fluoroazetidin-1-ium-1-ylidene)-7-(3-fluoroazetidin-1-yl)-5,5-dimethyl-3,5-dihydrodibenzo[b,e]silin-10-yl)benzoate (C10852S)**

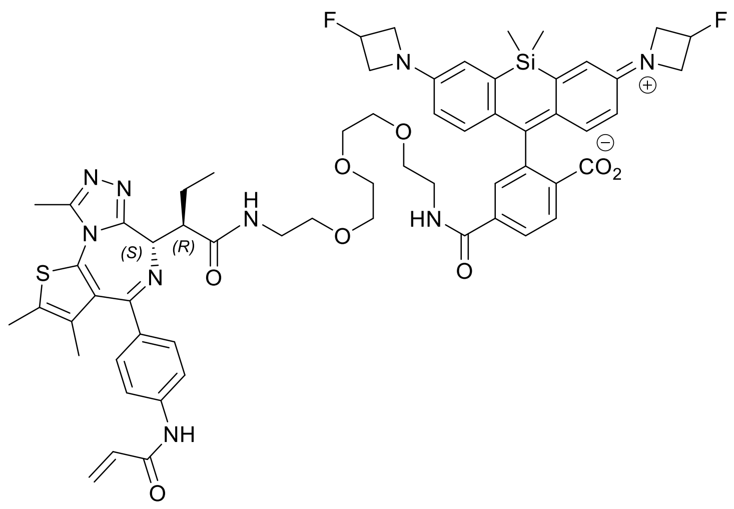

To a stirred solution of (2*R*)-2-[(9*S*)-7-[4-(tert-butoxycarbonylamino)phenyl]-4,5,13-trimethyl-3-thia-1,8,11,12-tetrazatricyclo[8.3.0.02,6]trideca-2(6),4,7,10,12-pentaen-9-yl]butanoic acid (**MR137**) (50 mg (uncorr.), 43 mg (corr.), 83.40 µmol (corr.)) in dichloromethane (3.0 mL) at ambient temperature was sequentially added bis(2,5-dioxopyrrolidin-1-yl) carbonate (43 mg, 167.86 µmol), *N,N*-dimethylpyridin-4-amine (2 mg, 16.37 µmol) and triethylamine (60 µL, 430.48 µmol) and the reaction mixture was stirred at ambient temperature for 45 minutes. After this time, a solution of 4-((2-(2-(2-(2-aminoethoxy)ethoxy)ethoxy)ethyl)carbamoyl)-2-(3-(3-fluoroazetidin-1-ium-1-ylidene)-7-(3-fluoroazetidin-1-yl)-5,5-dimethyl-3,5-dihydrodibenzo[b,e]silin-10-yl)benzoate (75 mg (uncorr.), 72 mg (corr.), 100.80 µmol (corr.)) in dichloromethane (2.0 mL) was added and the reaction mixture was stirred at ambient temperature for 20 hours before being concentrated under reduced pressure. Purification by flash column chromatography, eluting with 0-4% methanol in dichloromethane, afforded the Boc-protected precursor as a white solid (72 mg (uncorr.), 65 mg (corr.), 65% (corr.)). LCMS purity = 96.5%; m/z (ES+): 1198.30 $[M+H^+]^+$.

To a stirred solution of the Boc-precursor (72 mg (uncorr.), 54.07 µmol (corr.)) in dichloromethane (8 mL) was added TFA (0.40 mL) and the reaction mixture was stirred at 39 °C for 18 hours. After cooling to ambient temperature, the solution was treated with triethylamine (0.90 mL) followed by the portion-wise addition of acryloyl chloride diluted in dichloromethane (440 µL of acryloyl chloride dissolved in 1 mL of dichloromethane, 0.60 mL added), and the reaction mixture was stirred at ambient temperature for 4 hours before being concentrated under reduced pressure and reconcentrated from heptane. Purification by reversed phase chromatography, eluting with 10-70% acetonitrile (0.1% formic acid) in water (0.1% formic acid), was followed by conversion of the resulting formate salt to the free base by extractive isolation (dichloromethane and $NaHCO_3$ (aq.)) before lyophilisation from acetonitrile/water (1:1) afforded the title compound as an off-white solid (37 mg, 56%). $^1$H NMR (400 MHz, DMSO-d6): δ 10.31 (s, 1H), 8.80 (t, *J* = 5.5 Hz, 1H), 8.35 (t, *J* = 5.6 Hz, 1H), 8.08 (dd, *J* = 8.1, 1.2 Hz, 1H), 8.02 (d, *J* = 8.1 Hz, 1H), 7.68 (d, *J* = 8.9 Hz, 2H), 7.66 – 7.66 (m, 1H), 7.33 (d, *J* = 8.5 Hz, 2H), 6.81 (d, *J* = 2.6 Hz, 2H), 6.66 (d, *J* = 8.7 Hz, 2H), 6.46 – 6.38 (m, 3H), 6.25 (dd, *J* = 17.0, 2.0 Hz, 1H), 5.75 (dd, *J* = 10.1, 2.0 Hz, 1H), 5.46 (dtt, *J* = 57.7, 5.7, 3.0 Hz, 2H), 4.23 – 4.10 (m, 4H), 4.06 (d, *J* = 10.8 Hz, 1H), 3.97 – 3.83 (m, 4H), 3.56 – 3.44 (m, 13H), 3.40 – 3.33 (m, 4H), 2.58 (s, 3H), 2.40 (s, 3H), 1.92 – 1.80 (m, 1H), 1.63 (s, 3H), 1.48 – 1.35 (m, 1H), 0.88 (t, *J* = 7.4 Hz, 3H), 0.63 (s, 3H), 0.51 (s, 3H); $^{13}$C NMR (100 MHz, DMSO-d6): δ 172.88, 169.20, 164.88, 163.34, 162.36, 154.69, 154.60, 150.02, 150.01, 149.55, 140.99, 139.96, 135.75, 132.82, 132.18, 131.63, 130.73, 130.23, 129.95, 129.12, 128.28, 127.47, 127.36, 127.09, 125.54, 122.74, 118.62, 116.22, 113.46, 90.84, 83.48 (d, $J_{CF}$ = 200.1 Hz), 69.72, 69.61, 69.51, 69.41, 68.64, 59.24 (d, $J_{CF}$ = 23.4 Hz), 59.06, 49.34, 39.44, 38.49, 22.46, 14.06, 12.70, 11.59, 11.25, -0.10, -1.31; $^{19}$F NMR (376 MHz, DMSO-d6): δ -179.1; LCMS purity = 96.6%; m/z (ES+): 1152.30 $[M+H^+]^+$; HRMS (ES+) calculated for $[(C61H67F2N9O8SSi)+H^+]^+$ 1152.4649, found 1152.4647.

**FULL GELS and Western Blots**

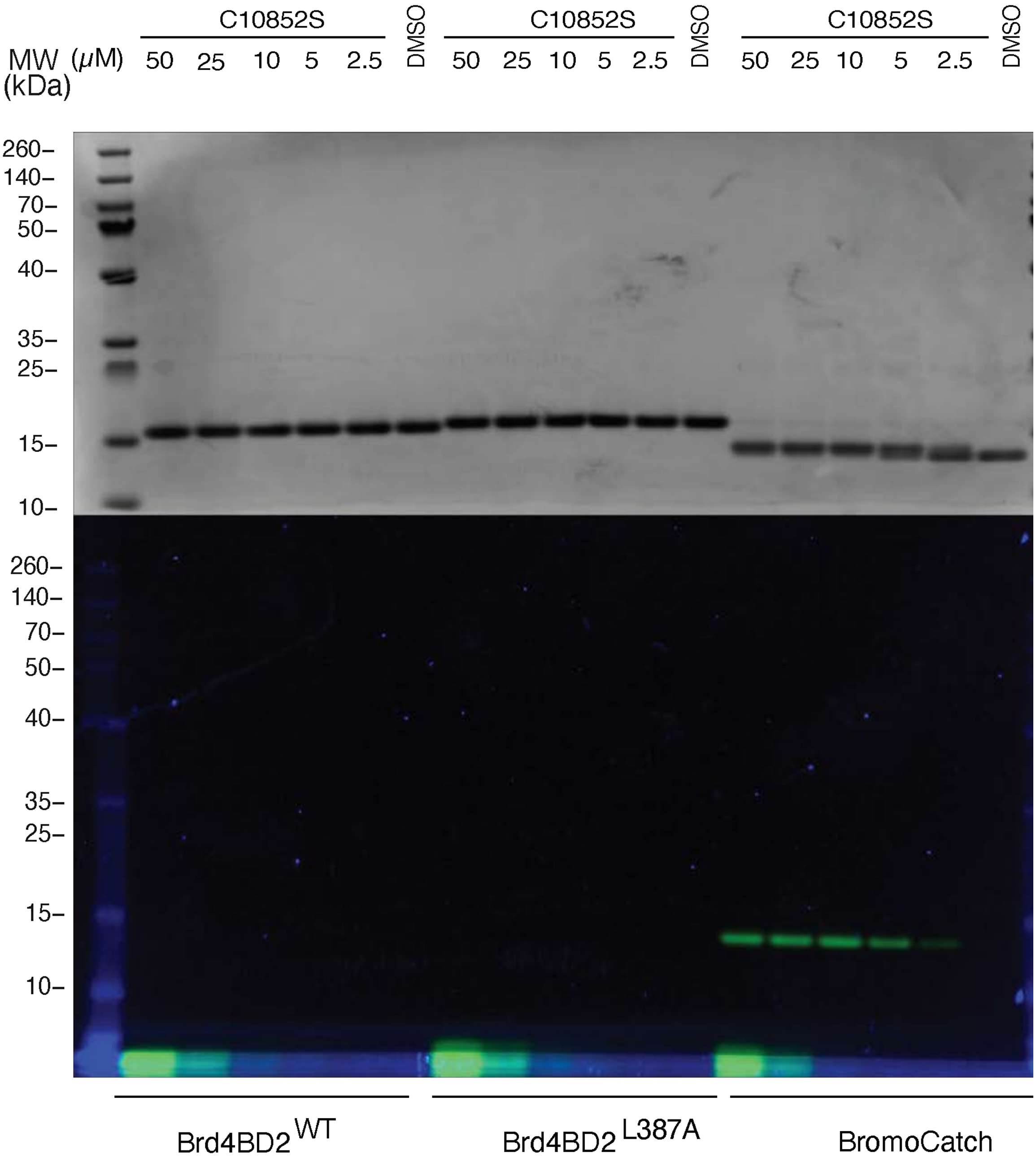


**Supplementary Figure 15.** Full length SDS gel of MR202 titration corresponding to Figure 6 in the main manuscript.

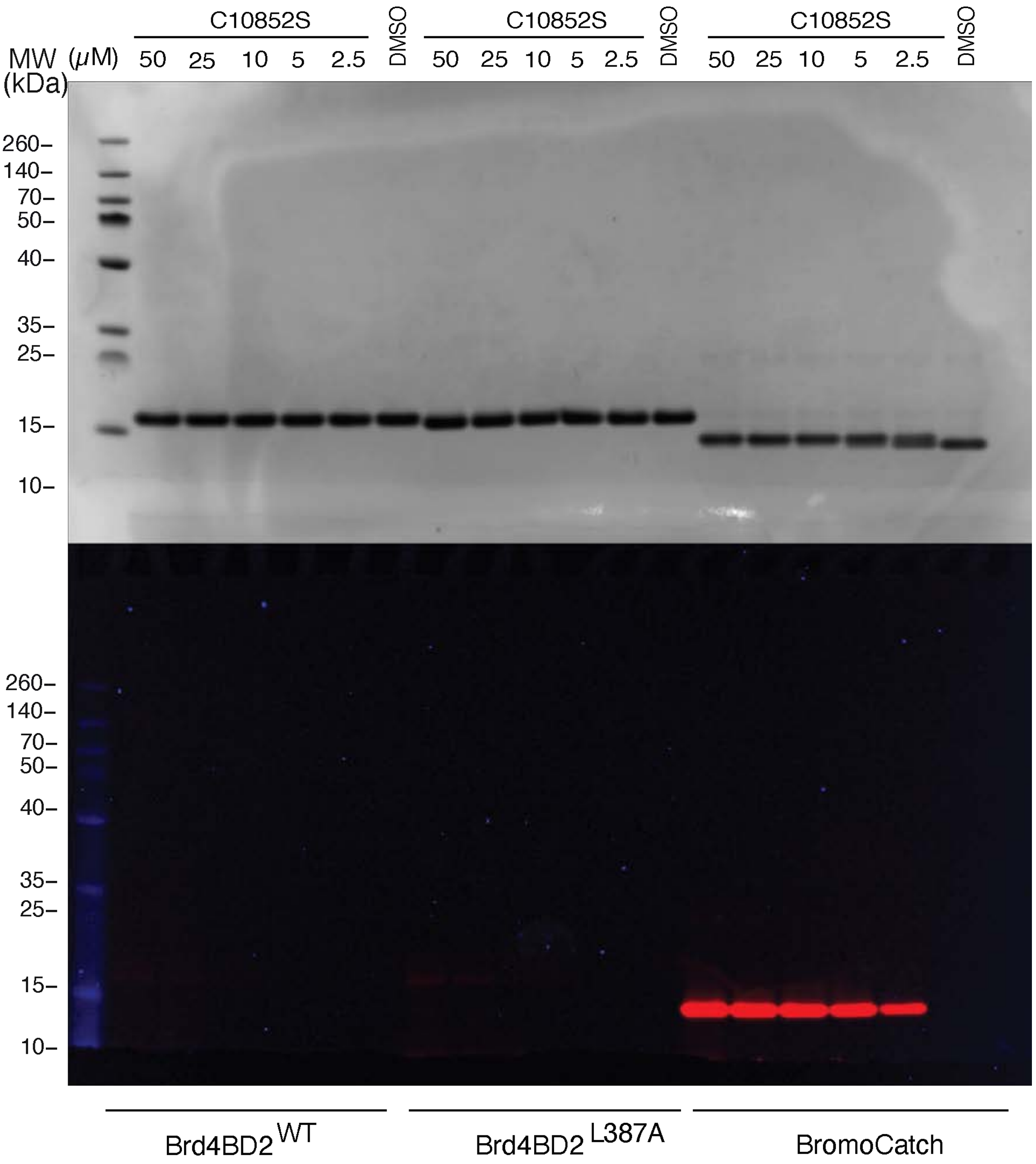


**Supplementary Figure 16.** Full length SDS gel of C18052S titration corresponding to Figure 7 in the main manuscript.

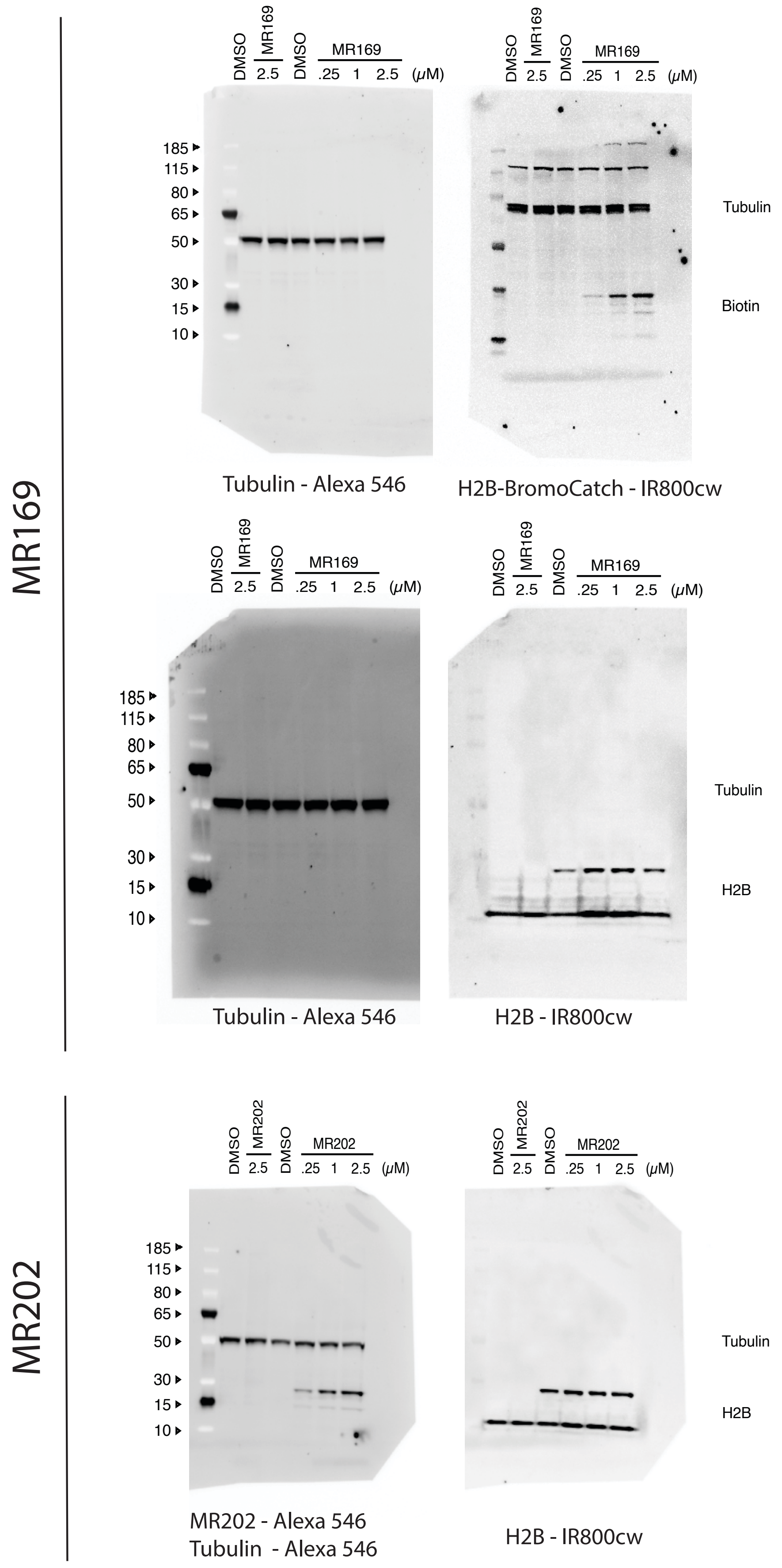


**Supplementary Figure 17.** Full length western blot membrane corresponding to Figure 6 in the main manuscript.

### HPLC-UV / LC-MS

#### MR155 – alkyne probe

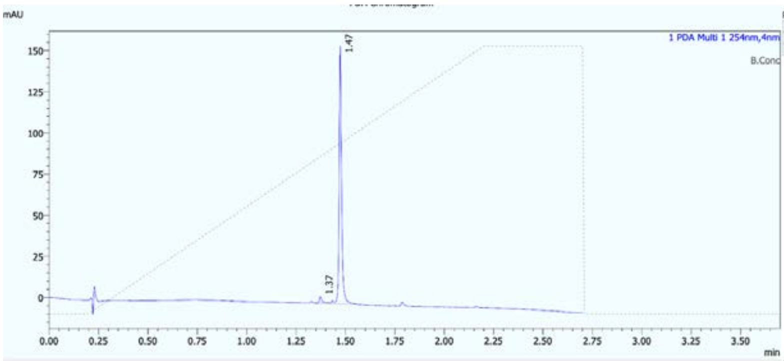


PDA Ch1 254nm

| Pk# | R.T. | Area | Height | Area% | UV max | S/N |
|---|---|---|---|---|---|---|
| 1 | 1.37 | 3865 | 3552 | 2.8 | 267/308/359 | 0.68 |
| 2 | 1.47 | 136237 | 148359 | 97.2 | 302/263 | 28.36 |

TIC(1-1)

| Pk# | R.T. | Area | Bs. Pk Int. | Area% | M/z | S/N |
|---|---|---|---|---|---|---|
| 2 | 1.47 | 10492143 | 2664114 | 100.000 | 633.5 | 10.1 |

TIC(1-2)

| Pk# | R.T. | Area | Bs. Pk Int. | Area% | M/z | S/N |
|---|---|---|---|---|---|---|
| 2 | 1.47 | 2103448 | 441746 | 100.000 | 631.2 | 7.0 |

#### MR170 – VHL covalent PROTAC

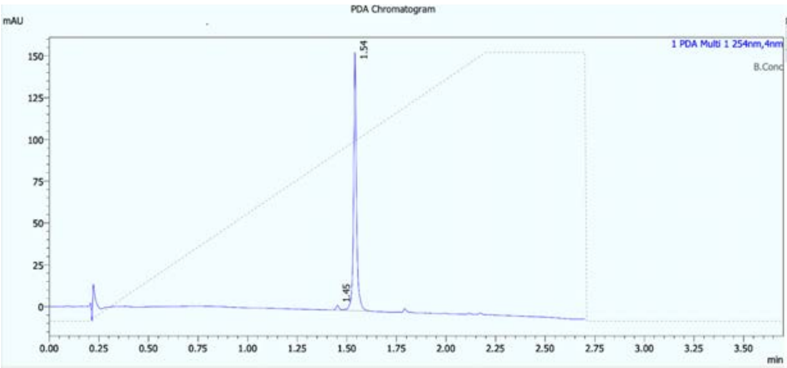


PDA Peak Table

PDA Ch1 254nm

| Pk# | R.T. | Area | Height | Area% | UV max | S/N |
|---|---|---|---|---|---|---|
| 1 | 1.45 | 3269 | 2772 | 2.0 | 268 | 0.42 |
| 2 | 1.54 | 162748 | 144254 | 98.0 | 267 | 21.86 |

TIC(1-1)

| Pk# | R.T. | Area | Bs. Pk Int. | Area% | M/z | S/N |
|---|---|---|---|---|---|---|
| 2 | 1.54 | 12690074 | 1316808 | 100.000 | 1065.7 | 18.8 |

TIC(1-2)

| Pk# | R.T. | Area | Bs. Pk Int. | Area% | M/z | S/N |
|---|---|---|---|---|---|---|
| 2 | 1.54 | 1349209 | 295332 | 100.000 | 1063.9 | 5.2 |

#### MR202 – TMR probe

UV

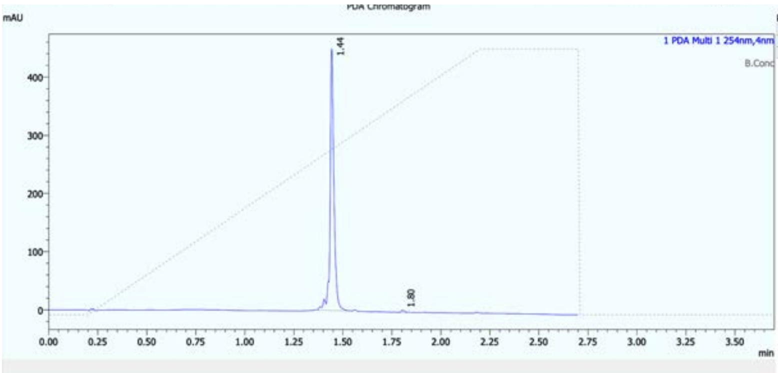


PDA Ch1 254nm

| Pk# | R.T. | Area | Height | Area% | UV max | S/N |
|---|---|---|---|---|---|---|
| 1 | 1.44 | 620390 | 444275 | 99.3 | 556/256/285/195/352 | 194.52 |
| 2 | 1.80 | 4125 | 3635 | 0.7 | 259/552/630/660 | 1.59 |

MS

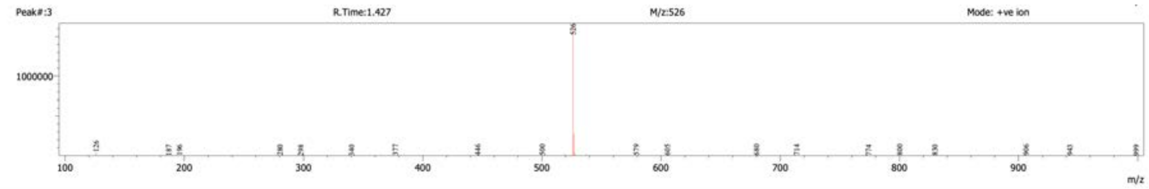

**MR169 – Biotin probe**

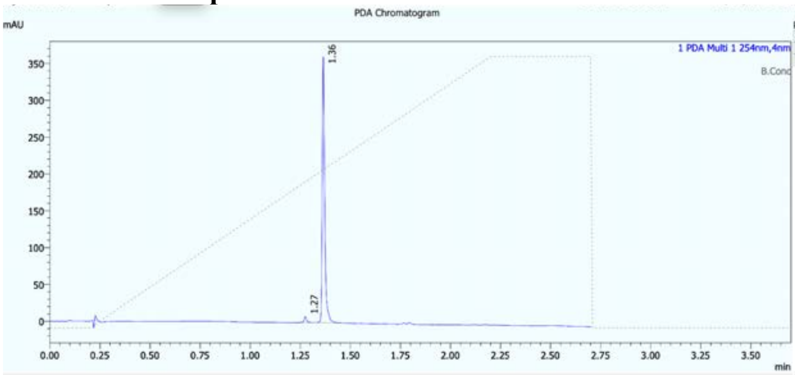


PDA Peak Table

PDA Ch1 254nm

| Pk# | R.T. | Area | Height | Area% | UV max | S/N |
|---|---|---|---|---|---|---|
| 1 | 1.27 | 6706 | 7961 | 2.1 | 267/307/357 | 1.51 |
| 2 | 1.36 | 311321 | 329828 | 97.9 | 302/263/231 | 62.70 |

TIC(1-1)

| Pk# | R.T. | Area | Bs. Pk Int. | Area% | M/z | S/N |
|---|---|---|---|---|---|---|
| 2 | 1.37 | 28410666 | 3774974 | 100.000 | 908.5 | 14.3 |

TIC(1-2)

| Pk# | R.T. | Area | Bs. Pk Int. | Area% | M/z | S/N |
|---|---|---|---|---|---|---|
| 2 | 1.36 | 5157416 | 714826 | 100.000 | 906.5 | 14.4 |

**C10852S – Janelia Fluor® 635 probe**

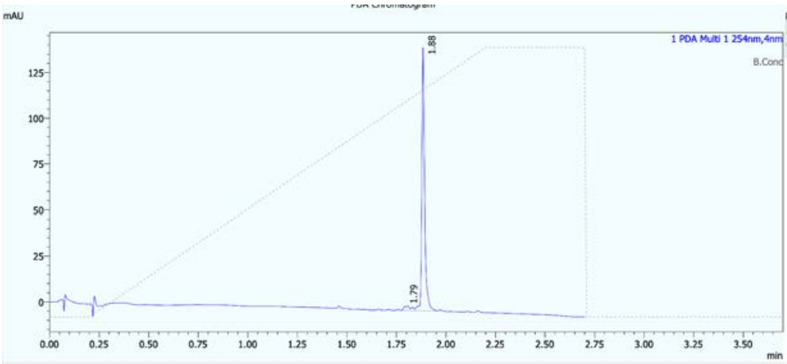


PDA Ch1 254nm

| Pk# | R.T. | Area | Height | Area% | UV max | S/N |
|---|---|---|---|---|---|---|
| 1 | 1.79 | 6136 | 2119 | 4.1 | 258/345 | 0.60 |
| 2 | 1.88 | 143051 | 136651 | 95.9 | 267 | 38.53 |

TIC(1-1)

| Pk# | R.T. | Area | Bs. Pk Int. | Area% | M/z | S/N |
|---|---|---|---|---|---|---|
| 2 | 1.88 | 2842961 | 1418341 | 100.000 | 577.2 | 4.4 |
